# Rescuing Neurodevelopmental Deficits in AMPA Receptor Gain-of-Function Mutant

**DOI:** 10.64898/2025.12.30.697121

**Authors:** Chih-Ming Chen, Yu-Min Huang, Chih-Ching Chung, Richard C. Johnson, Cheng-Ya Tsai, Yu-Han Chen, Han L. Tan, Fu-Yun Hsiao, Richard L. Huganir, Shu-Ling Chiu

## Abstract

AMPA receptors (AMPARs) mediate fast excitatory synaptic transmission and are essential for neuronal development and brain function. We investigated the role of a recurrent variant in the AMPAR GluA1 subunit *(GRIA1* p.A636T) identified in individuals with autism spectrum disorder (ASD) and intellectual disability (ID). To test causality and mechanism, we generated a *Gria1*-A636T knock-in mouse model. Mutant mice exhibited core ASD/ID-like behaviors and a selective hippocampal vulnerability characterized by progressive dendritic atrophy and neuronal loss. Despite reduced GluA1-containing complexes, AMPARs displayed synaptic hyperexcitability and failed to undergo the normal postnatal transition to calcium-impermeable AMPARs, resulting in persistent excitotoxicity. To explore therapeutic intervention, we designed an allele-specific antisense oligonucleotide to specifically silence the mutant transcript. A single neonatal administration of the antisense oligonucleotide entirely prevented hippocampal pathology and ameliorated behavioral deficits. These findings establish *GRIA1*-A636T as a gain-of-function mutation that drives developmental excitotoxicity and highlight the potential of RNA-targeted precision medicine for neurodevelopmental disorders.

## Introduction

Neurodevelopmental disorders (NDDs), including autism spectrum disorder (ASD) and intellectual disability (ID), profoundly affect neural circuit function and cognition and often lead to lifelong disability. Despite their heterogeneous and complex etiology, advances in human genetics have identified numerous rare and recurrent genetic variants that are strongly associated with NDD pathology. Among these, mutations affecting glutamatergic synapse function, particularly those affecting AMPA-type glutamate receptors (AMPARs), have emerged as important contributors, given the central role of AMPARs in excitatory neurotransmission and synaptic plasticity during neurodevelopment^1–5^.

AMPARs are tetrameric glutamate-gated ion channels composed of four subunits (GluA1-4) that mediate the majority of fast excitatory synaptic transmission across brain regions critical for cognition. Dysfunction of AMPAR has been implicated in diverse neuropsychiatric and cognitive disorders, highlighting the importance of precise regulation of AMPAR activity for proper brain functions^2–4^. Recently, a recurrent *de novo* variant (c.1906 G>A, resulting in p.Ala636Thr [A636T]) in the *GRIA1* gene encoding the GluA1 subunit, was identified as a hotspot mutation in a large-scale genetic screen of individuals with NDDs^6^. Several independent clinical studies have also reported its link to ASD and ID^4,6,7^. Notably, AMPARs are expressed at particularly high density in the hippocampus, where GluA1 is the predominant AMPA subunit relative to other subunits^5^, suggesting that hippocampal circuit may be especially vulnerable to GluA1 dysfunction and contribute to NDD-associated behavior phenotypes. However, it remains unclear whether *GRIA1-*A636T alone is sufficient to drive NDD pathogenesis. In addition, the molecular and cellular mechanisms underlying its effects on neural function and behavior remain unknown.

The *Gria1-*A636T mutation is located within a highly conserved gating motif in the pore-forming transmembrane domain 3 of the GluA1 (Fig. 1a, b). This motif is homologous across species and among ionotropic glutamate receptor subtypes, including AMPA, NMDA (*GRIN1*), and kainate (*GRIK1*) receptors, as well as the glutamate receptor-like gene *GRID2* (Fig. 1a). Notably, an analogous mutation (A654T) in GluD2, known as the “Lurcher” mutation, has been shown to cause constitutive channel opening, severe Purkinje cell neurodegeneration, and profound ataxia in mice^8,9^. Although prior heterologous expression studies suggest that A636T can alter AMPAR gating properties^7,10,11^, its impact on synaptic transmission, neuronal structure, circuit development and behavior *in vivo*, and its relevance to human NDD pathology, remains unclear.

**Fig. 1.**
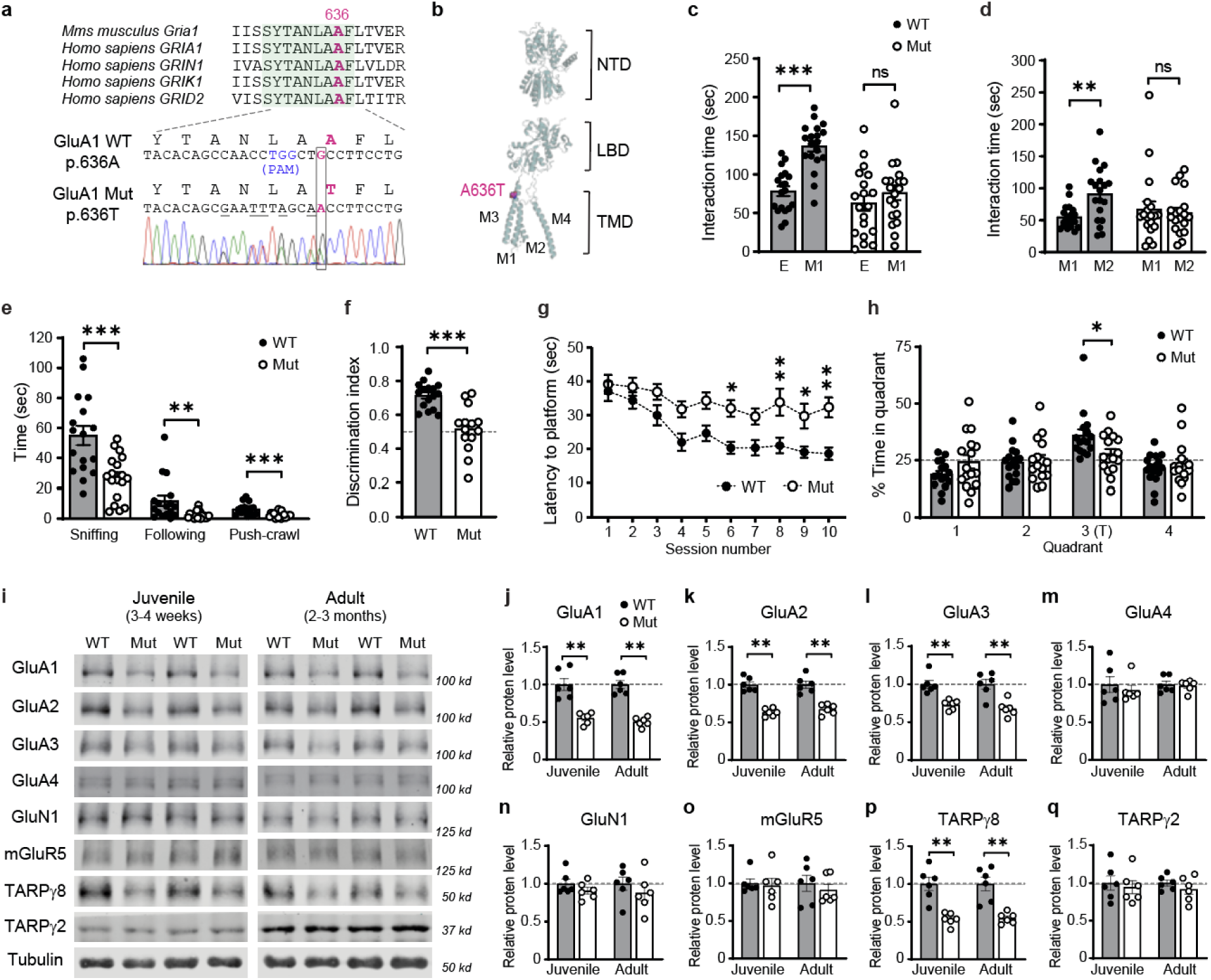
*Gria1*-A636T heterozygous (*Gria1* Mut) mice exhibit social-cognitive deficits and reduced AMPAR expression. **a**, Sanger sequencing confirmed the heterozygous c.1906 G>A (p.A636T) substitution in *Gria1*. The variant (magenta) is located within a conserved gating motif (highlighted in green) across species and glutamate receptor family members. The protospacer adjacent motif (PAM, blue) and CRISPR-targeted restriction sites (underlined) are also indicated. **b**, GluA1 subunit schematic showing the A636T residue within the M3 helix. NTD: N-terminal domain, LBD: ligand-binding domain, TMD: transmembrane domain. M1-4: transmembrane helices. **c**, **d**, In the three-chamber social interaction test, WT mice showed a significant preference for a conspecific (M1) over an empty cup (E) (P<0.0001), and for a novel mouse (M2) over a familiar one (M1) (P=0.0037). In contrast, Mut mice showed no preference in either case, indicating impaired sociability and social novelty recognition. **e**, In the reciprocal social interaction test, Mut mice showed reduced sniffing, following, and push-crawl behaviors (All P<0.005). **f**, In the Y-maze, WT mice preferred the novel arm, whereas Mut mice did not. The discrimination index was significantly lower in Mut mice (P<0.0001). The dashed line indicates chance. **g**, In the Morris water maze (MWM) training, Mut mice showed impaired spatial learning, with longer latency to locate the platform at later sessions (P<0.05). **h**, In the MWM probe trial, WT mice spent more time in the target (T) quadrant, whereas Mut mice did not (P=0.027). The dashed line indicates chance. **i**, Representative Western blots of hippocampal glutamate receptor subunits and auxiliary proteins from juvenile and adult mice. **j-q**, Quantification of (**i**) revealed reduced GluA1-3, and TARPγ8 levels in Mut mice at both ages (P<0.005). No changes in GluA4, GluN1, mGluR5, or TARPγ2 were detected. Dashed lines indicate normalized WT levels (=1). Data are presented as mean ± s.e.m. *P < 0.05, **P < 0.01, ***P < 0.001. Exact P values, sample sizes, and statistical tests are provided in the Supplementary Table.

To address these questions, we developed a mouse model carrying the clinically identified *Gria1*-A636T mutation. We demonstrate that this mutation alone is sufficient to induce core behavioral phenotypes resembling human ASD and ID. Moreover, we show that *Gria1-*A636T profoundly alters AMPAR gating, resulting in enhanced excitatory synaptic currents and persistent calcium-permeable AMPAR activity. These changes lead to significant dendritic remodeling, excitotoxic cell death, neuroinflammation, and progressive hippocampal atrophy, highlighting a selective vulnerability of the hippocampal circuit.

Furthermore, we developed a targeted therapeutic strategy utilizing antisense oligonucleotides (ASOs) specifically designed to reduce the mutant *Gria1* transcripts. ASO therapies have emerged as precise and stable modulators of gene expression with significant clinical promise for treating genetic and neurological disorders^12–14^. Remarkably, a single ASO administration effectively prevented neuronal loss, resolved neuroinflammation, restored dendritic architecture, and recovered the social and cognitive behaviors of *Gria1*-A636T mice, illustrating the therapeutic potential of this strategy.

Together, our findings establish *Grai1*-A636T as a causal, gain-of-function mutation that drives severe ASD/ID pathology through AMPAR-mediated excitotoxicity. Our study elucidates critical molecular mechanisms underlying AMPAR-related NDDs and highlights ASO-based precision medicine as a promising therapeutic strategy for individuals carrying this mutation.

### *Gria1*-A636T mice recapitulate core ASD/ID-like behavioral traits

To determine whether the human variant *GRIA1*-A636T (c.1906 G>A; p.A636T) identified in individuals with NDDs, is sufficient to drive disease-related phenotypes, we generated a mouse model harboring the identical mutation using CRISPR/Cas9 genome editing (Fig. 1a, b). Heterozygous *Gria1*-A636T (*Gria1* Mut) mice were viable, exhibited normal Mendelian inheritance, and displayed typical physical development. However, homozygous mutants were lethal, indicating that the A636T mutation exerts a dose-dependent effect on viability. All subsequent experiments were performed with *Gria1* Mut mice to model the *de novo* heterozygous mutation found in patients.

To assess ASD-like behaviors, we first used the three-chamber test, a standard assay for measuring social preference in rodents. *Gria1* Mut mice exhibited impaired social preference, spending comparable time interacting with a novel mouse and an object (Fig. 1c), and failed to show social novelty recognition, interacting equally with a novel and a familiar mouse (Fig. 1d). To validate these findings in a more natural setting, we employed reciprocal social interaction assays to access social behavior between two freely moving mice. Consistently, Mut mice exhibited significantly reduced social behaviors, including sniffing, following and push-crawl interactions (Fig. 1e), confirming impaired social function.

We next evaluated cognitive impairments associated with ID using spatial memory tasks. In the Y-maze test, Mut mice failed to show a preference for the novel arm, indicating a deficit in short-term spatial memory (Fig. 1f). In the Morris water maze, wild-type (WT) mice rapidly learned the platform location and reached a plateau performance by the sixth session. In contrast, Mut mice showed minimal improvement across training sessions (Fig. 1g), indicating an impairment in spatial learning. During the probe trial, WT mice spent significantly more time in the target quadrant, whereas Mut mice showed no quadrant preference, indicating impaired memory retention (Fig. 1h). Mut mice also performed poorly in the reversal learning phase, failing to learn the new platform location and retain spatial memory (Extended Data Fig. 1a, b).

Beyond major ASD/ID traits, we assessed repetitive behaviors, motor function, and anxiety-like phenotypes. Mut mice showed increased repetitive self-grooming activities (Extended Data Fig. 1c), a hallmark of stereotypic behaviors in ASD. Anxiety-like behaviors were also evident, with Mut mice spending more time in the periphery during the open field test (Extended Data Fig. 1f) and preferring the closed arms in the elevated plus maze (Extended Data Fig. 1g). Locomotor activity was comparable between genotypes, excluding gross motor deficits (Extended Data Fig. 1d, e).

Together, these behavioral findings establish *Gria1*-A636T as a pathogenic mutation capable of recapitulating ASD and ID-like phenotypes, including social deficits, repetitive behaviors, learning and memory impairments, and increased anxiety.

### *Gria1*-A636T mutation selectively reduces GluA1-associated AMPAR complexes

We examined molecular alterations that may underlie these behavioral phenotypes by assessing AMPAR protein expression across brain regions implicated in ASD and ID. Western blot analyses revealed a marked reduction in GluA1 protein in Mut mice, with the hippocampus showing the strongest effect (a 45% reduction in juveniles and 50% reduction in adults; Fig. 1i, j). Moderate reductions were also observed in other regions relevant to ASD/ID, including the prefrontal cortex, somatosensory cortex, thalamus, and cerebellum (20-36% in juveniles; 29-38% in adults; Extended Data Fig. 2).

Consistent with the critical role of GluA1 in assembling functional AMPARs, hippocampal levels of GluA2, GluA3, and the auxiliary protein TARPγ8 were also selectively reduced in Mut mice (Fig. 1k, l, p). In contrast, GluA4 and TARPγ2, which are typically incorporated into distinct AMPAR complexes or are enriched in non-hippocampal regions, were unaffected (Fig. 1m, q). Notably, RT-qPCR analysis showed no changes in mRNA levels for AMPAR subunits or auxiliary proteins in both juvenile and adult stages (Extended Data Fig. 2u, v), suggesting that the observed protein reductions arise from post-translational mechanisms rather than altered transcription.

Notably, the expression levels of other glutamate receptor components, including GluN1 (the obligate NMDAR subunit) and mGluR5 (a major metabotropic glutamate receptor), remained largely unchanged in the Mut hippocampus (Fig. 1n, o). Together, these biochemical analyses demonstrate that the *Gria1*-A636T selectively disrupts GluA1-containing AMPAR complexes, particularly in the hippocampus.

### *Gria1-A636T* mutation selectively disrupts hippocampal structure and neuronal morphology

To identify brain regions affected by the *Gria1-A636T* mutation, we also performed non-invasive magnetic resonance imaging (MRI) to assess brain structure in live *Gria1* mice at juvenile and adult stages. MRI-based segmentation and volumetric analysis revealed a significant and selective reduction in hippocampal volume, with an approximately 21% reduction observed at the juvenile stage (Fig. 2a, c). In contrast, the volumes of the neocortex (Fig. 2d), thalamus, striatum, ventricles and total brain remained unchanged at this stage (Extended Data Fig. 3).

**Fig. 2.**
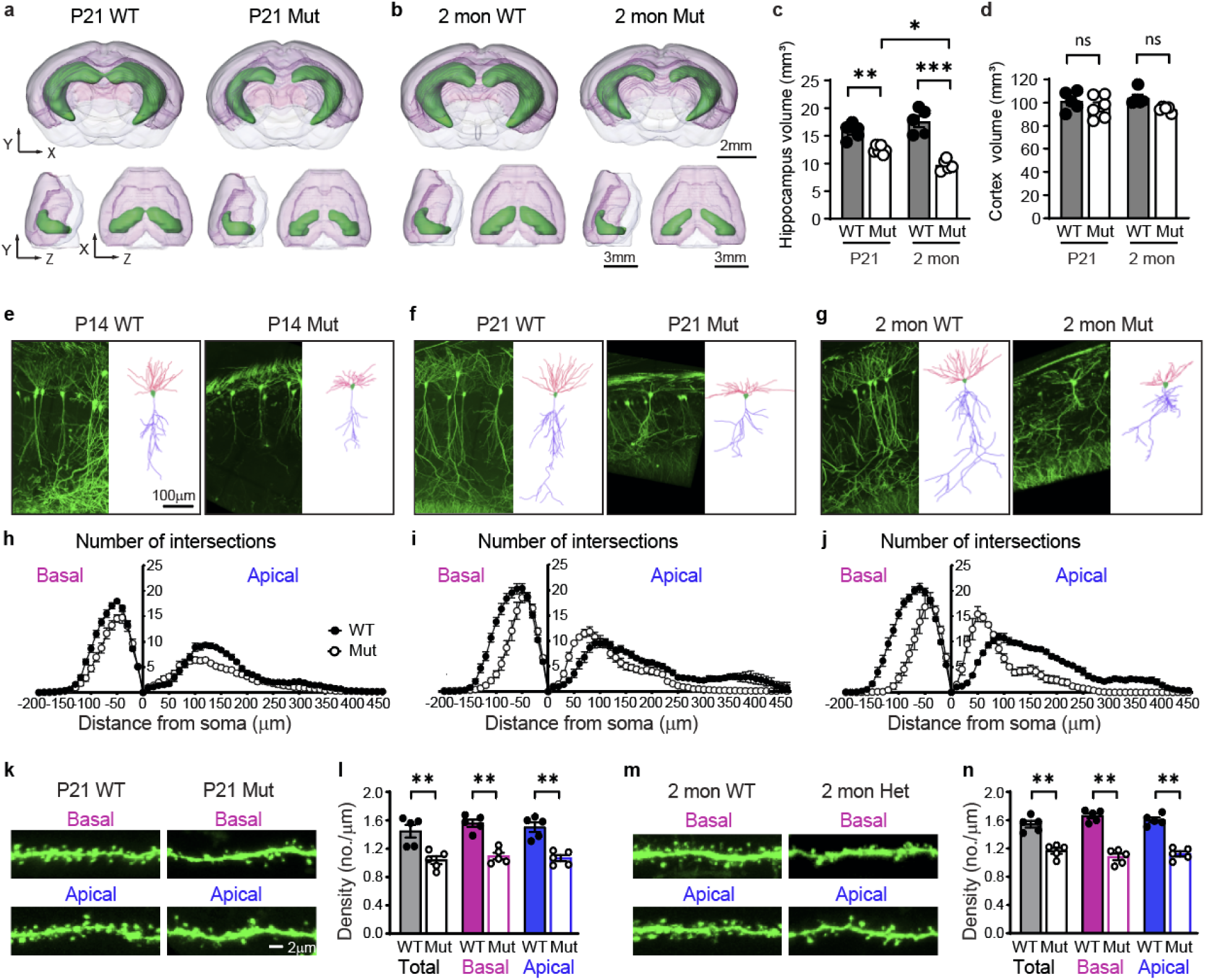
Selective hippocampal atrophy and dendritic remodeling in *Gria1* Mut mice. **a, b**, Coronal, sagittal and axial MRI images showing 3D brain reconstructions of WT and Mut mice at juvenile (**a,** P21) and adult (**b,** 2 months) stages. **c, d,** Quantification of brain volume reveals progressive hippocampal atrophy in Mut mice compared to WT littermate controls (∼21% reduction at juvenile, ∼45% reduction at adult, P<0.005), while cortical volume remained unchanged. **e-g,** Maximum-intensity projections and 3D reconstructions of GFP-labeled CA1 pyramidal neurons at P14 (**e**), P21 (**f**), and 2 months (**g**). **h-j,** Sholl analyses showed progressively reduced dendritic arbor complexity and field coverage in Mut mice. Significant genotype differences were observed at multiple radial distances (n=5 neurons/animal, 5 animals/group, detailed statistics in Supplementary Table). **k-n,** High magnification images and spine density quantification of basal and apical dendrites at P21 (k, l) and 2 months (m, n) showed significantly reduced spine density in Mut neurons. (P=0.008, n=5 animals/group; 5 neurons/animal; 5 dendrites/neuron). Data are presented as mean ± s.e.m. *P < 0.05, **P < 0.01, ***P < 0.001. Exact P values, sample sizes, and statistical tests are provided in the Supplementary Table.

As mice matured, hippocampal atrophy progressively worsened, reaching a volume reduction of ∼45% by adulthood (Fig. 2b, c). This was accompanied by a modest reduction in total brain volume (Extended Data Fig. 3c), while the sizes of other brain regions analyzed remained largely unaffected (Fig. 2d, Extended Data Fig. 3d-f). These MRI imaging findings highlight the hippocampus as a region of increased vulnerability to the *Gria1-*A636T mutation and demonstrate a progressive, age-dependent deterioration that parallels the molecular abnormalities observed in Mut mice.

To further elucidate the cellular basis of hippocampal atrophy, we crossed *Gria1*-A636T mice with Thy1-GFP transgenic mice to sparsely label individual pyramidal neurons for morphological analysis across developmental stages. Starting at postnatal day 14 (P14), the earliest time point at which sufficient GFP signal could be detected, CA1 pyramidal neurons with clearly distinguishable dendritic arbors were reconstructed from full z-stack confocal images. Morphological analysis revealed a progressive reduction in dendritic field size in Mut mice compared to WT littermates at pre-weaning (P14), weaning (P21), and early adulthood (2 months). Sholl analysis showed that the dendritic arbors in Mut neurons were more compact and shifted toward the soma (Fig. 2e-g). Concurrently, the total dendritic branch length was significantly reduced by approximately 20% at P14, 23% at P21, and 33% at 2 months (Extended Data Fig. 4a-d), indicating an age-dependent deterioration in dendritic architecture. Notably, these deficits were confined to the hippocampus, as pyramidal neurons in the somatosensory cortex remained unaffected (Extended Data Fig. 5).

Given the enrichment of AMPARs, particularly GluA1, at postsynaptic sites and dendritic spines, we further examined spine density in CA1 pyramidal neurons. Spine quantification revealed a significant reduction in both apical and basal dendrites of Mut mice (Fig. 2h-i), suggesting a global loss of excitatory synaptic connectivity.

Collectively, these results demonstrate that dendritic remodeling and spine loss are key morphological correlates of hippocampal atrophy in *Gria1*-A636T mice. These progressive structural changes likely reflect disrupted neuronal development and persistent synaptic dysfunction, contributing to the selective vulnerability of the hippocampus.

### *Gria1*-A636T mutation enhances AMPAR-mediated excitatory transmission

To assess the impact of the A636T mutation on AMPAR function, we performed whole-cell patch-clamp recordings of CA1 pyramidal neurons in acute hippocampal slices derived from juvenile *Gria1* WT and Mut littermates (Fig. 3a). Because the *Gria1*-A636T substitution is homologous to the constitutively active *Griδ2*-Lurcher mutation^8,9^, we first tested whether *Gria1*-A636T produces basal AMPAR leak currents. Holding currents and membrane resistance were unchanged in Mut neurons (Fig. 3b, 3c), and AMPAR blockade with NBQX had no effect on resting holding currents (Fig. 3e), indicating the absence of constitutive AMPAR conductance. Mut neurons, however, exhibited reduced capacitance (Fig. 3d), consistent with the apparent dendritic atrophy observed morphologically.

**Fig. 3.**
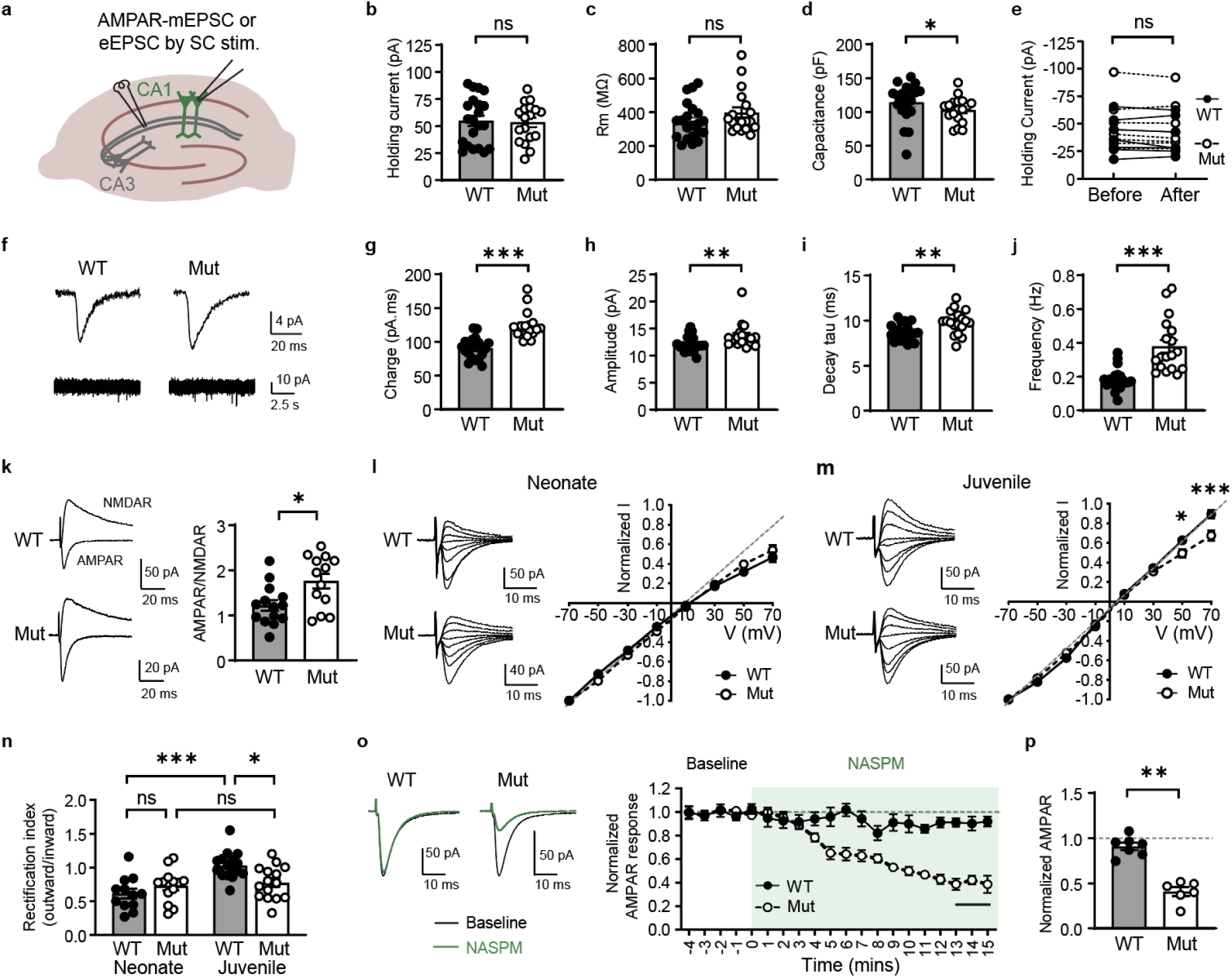
Enhanced AMPAR-mediated synaptic transmission and calcium permeability in *Gria1* Mut mice. **a**, Schematic of patch clamp setup for recording AMPAR-mEPSCs and eEPSCs from CA1 pyramidal neurons. SC: Schaffer collateral. ***b-d,*** Baseline membrane properties of CA1 neurons from juvenile mice. No differences in holding current (b, P=0.900) or membrane resistance (c, P=0.477) were observed, indicating no AMPAR-mediated leak in Mut neurons. Cell capacitance was modestly reduced in Mut mice (d, P=0.046). **e,** NBQX application had no effect on resting current, confirming the absence of basal AMPAR leak in Mut neurons. (P>0.999). **f-j**, AMPAR-mEPSC recordings showed increased charge transfer (**g,** P<0.0001), amplitude (**h,** P = 0.005), decay time (**i,** P=0.004), and frequency (**j,** P < 0.0001) in Mut neurons. **k,** AMPAR/NMDAR eEPSC ratio was increased in Mut neurons (P=0.015), indicating enhanced AMPAR-mediated synaptic drive. **l,** Normalized current-voltage (I-V) relationships of AMPA eEPSC in neonatal neurons. Both WT and Mut neurons showed inward rectification, consistent with the normal early developmental expression of CP-AMPARs at this stage. **m,** At the juvenile stage, WT neurons exhibited near-linear I-V relationships, whereas Mut neurons retained inward rectification, reflecting persistent AMPAR calcium permeability (+50 mV, P=0.012; +70mV, P<0.0001). **n,** Rectification index (RI) increased from neonate to juvenile stages in WT mice (P=0.0001), but remained consistently low in Mut mice (P>0.999), indicating persistent CP-AMPAR expression in Mut mice at the juvenile stage. **o,** Representative traces and time course of AMPAR eEPSC peak amplitude during NASPM application in juvenile CA1 neurons. The final stable phase (last 3 min) is indicated. **p,** Quantification of the NASPM stable phase (**o**) showed a significant reduction of AMPAR eEPSCs in Mut, but not WT neurons (P=0.001). Data are presented as mean ± s.e.m. *P < 0.05, **P < 0.01, ***P < 0.001. Exact P values, sample sizes, and statistical tests are provided in the Supplementary Table.

Miniature AMPAR-mediated excitatory postsynaptic current (mEPSC) recordings revealed increased charge transfer, amplitude, prolonged decay kinetics, and elevated mEPSC frequency compared to WT controls (Fig. 3f-j). These changes demonstrate gain-of-function AMPAR activity and are consistent with prior *in vitro* studies showing that the A636T mutation slows AMPAR deactivation, prolongs synaptic responses and increases glutamate sensitivity^7,10,11^. Additionally, the AMPAR/NMDAR current ratio was significantly elevated by 1.44-fold in Mut neurons (Fig. 3k), consistent with increased AMPAR synaptic drive in the juvenile hippocampus.

To determine whether these synaptic alterations emerge early in development, we recorded AMPAR-mEPSCs and eEPSC from neonatal CA1 neurons (∼P7, Extended Data Fig. 6). Baseline membrane properties, including holding current, membrane resistance, and capacitance, were comparable between Mut and WT neurons (Extended Data Fig. 6b-d), confirming the absence of early AMPAR leak currents. Nevertheless, Mut neurons already exhibited increased charge transfer, prolonged decay kinetics and elevated mEPSC frequency at this stage (Extended Data Fig. 6e, g, h), indicating early excitatory synaptic hyperactivity. In contrast, AMPAR-mEPSC amplitude and AMPAR/NMDAR ratios remained unchanged in neonatal Mut neurons (Extended Data Fig. 6f, i), suggesting that additional AMPAR regulatory mechanisms emerge later during postnatal maturation.

### *Gria1*-A636T mutation disrupts AMPAR calcium regulation during development

In the hippocampus, AMPARs are predominantly assembled as calcium-impermeable heterotetramers, most commonly GluA1/GluA2 or GluA2/GluA3 heteromers, which conduct monovalent ions^15^. A smaller population of GluA2-lacking calcium-permeable (CP) AMPARs, which are typically GluA1 homomers or GluA1/GluA3 heteromers, is expressed during early postnatal development^16–18^, or transiently during specific forms of synaptic plasticity^16,19^. While physiological calcium influx through CP-AMPARs is crucial for calcium-dependent signaling and synaptic plasticity underlying cognitive function^20–23^, prolonged or excessive CP-AMPAR activation can trigger excitotoxicity and leading to neurological disorders^16,24,25^.

Because CP-AMPARs exhibit higher conductance and reduced voltage-dependent polyamine block^26,27^, their persistence beyond early development would be expected to enhance AMPAR-mediated synaptic currents at the juvenile stage. We therefore tested whether the *Gria1*-A636T disrupts the normal developmental reduction of AMPAR calcium permeability. AMPAR calcium permeability was assessed by measuring voltage-dependent inward rectification, a biophysical signature of CP-AMPARs resulting from intracellular polyamine block.

At neonatal stages (P7), both WT and Mut neurons exhibited comparable inwardly rectifying AMPAR currents, characterized by reduced current at positive potentials and similar rectification indices (Fig. 3l, n), consistent with normal developmental expression of CP-AMPARs at this stage. By the juvenile stage, WT neurons displayed near-linear current–voltage (I-V) relationships, indicative of dominant calcium impermeable-AMPAR expression (rectification index [RI] =1.07±0.06; Fig. 3m, n). In contrast, Mut neurons failed to undergo this developmental transition and maintained pronounced inward rectification across development (RI= 0.78±0.06; Fig. 3m, n), indicating persistent AMPAR calcium permeability.

To directly assess the functional contribution of CP-AMPARs in the juvenile hippocampal circuit, we recorded AMPAR-eEPSC at Schaffer collateral-CA1 synapses with the application of NASPM, a selective, use-dependent CP-AMPAR antagonist. NASPM had minimal effects on AMPA-eEPSC amplitudes in WT neurons, but significantly reduced eEPSC amplitudes by ∼56% in Mut neurons (Fig. 3o-p). Together, these results provide functional evidence that the *Gria1*-A636T prevents the normal developmental attenuation of synaptic AMPAR calcium permeability, resulting in a substantial CP-AMPAR component to excitatory synaptic transmission at the juvenile stage.

### *Gria1*-A636T mutation promotes calcium influx and excitotoxic vulnerability

To determine whether the *Gria1-*A636T mutation is sufficient to drive calcium entry through AMPARs in a cell-autonomous manner, we transfected cultured hippocampal neurons with WT or A636T *Gria1* constructs together with GCaMP8s to monitor AMPAR-mediated calcium signals and mCherry to visualize dendritic and spine morphology. Following a brief 5-second pulse of 0.5 μM AMPA in ACSF containing TTX and APV to isolate AMPAR-mediated responses, neurons expressing GluA1-WT showed minimal or undetectable GCaMP responses, consistent with the modest amplitude and fast kinetics of AMPAR-mediated calcium influx under these conditions (Fig. 4a). In contrast, neurons expressing GluA1-A636T exhibited a robust (∼4-fold) transient calcium rise that returned to baseline after AMPA washout (Fig. 4a, c). This calcium signal was abolished by NBQX, demonstrating that A636T-containing AMPARs are sufficient to drive large AMPAR-dependent calcium responses (Fig. 4a, c, e).

**Fig. 4.**
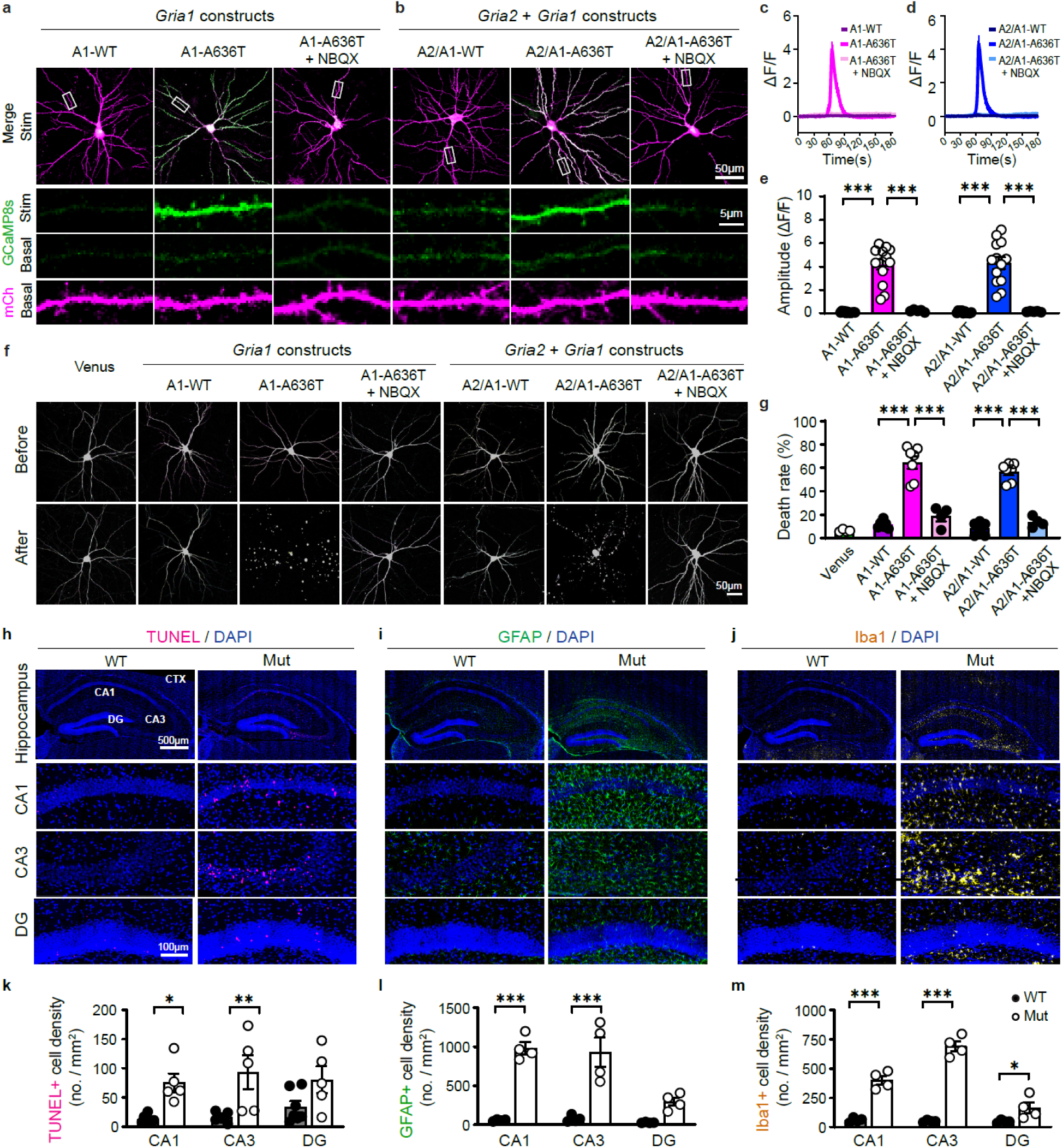
*Gria1*-A636T increases calcium permeability and induces excitotoxic neuronal death and neuroinflammation. **a-b**, Representative images of cultured hippocampal neurons co-transfected with GCaMP8s, mCherry and *Gria1* constructs (A1-WT or A1-A636T), or *Gria2* + *Gria1* constructs (A2/A1-WT or A2/A1-A636T), with or without NBQX co-application during AMPA stimulation. **c-d,** Averaged traces of AMPA-evoked calcium transient in respond to a brief AMPA pulse and washout. **e,** Quantification of peak calcium amplitudes showing markedly elevated responses in A1-A636T-expressing neurons, irrespective of GluA2 co-expression, which were abolished by NBQX. **f,** Representative images of Venus-labeled neurons before and after 16-hr AMPA exposure. Expression of A1-A636T or A2/A1-A636T induced dendritic fragmentation and neuronal loss that was prevented by NBQX. **g,** Quantification of neuron death rates in (**f**). **h,** Representative TUNEL staining images showing increased cell death in the hippocampus of *Gria1* Mut mice, with magnified views of the CA1, CA3, and DG subregions. DG: dentate gyrus; CTX: cortex. **i,** GFAP immunostaining showed increased astrogliosis in the Mut hippocampus. **j,** Iba1 immunostaining showed increased microglia activation in the Mut hippocampus. **k,** Quantification of TUNEL+ cell density at P21 showed significantly increased cell death in CA1 (P=0.014) and CA3 (P=0.002), but not in DG (P=0.21). **l-m,** Quantification of GFAP+ astrocytes (**l**), and Iba1+ microglia (**m**) densities showed marked neuroinflammation in CA1 and CA3 of Mut mice (all P<0.0001). Data are presented as mean ± s.e.m. *P < 0.05, **P < 0.01, ***P < 0.001. Exact P values, sample sizes, and statistical tests are provided in the Supplementary Table.

AMPAR calcium permeability is normally restricted by GluA2 incorporation, as RNA editing at the Q/R site in GluA2 introduces a positive charge within the channel pore that repels calcium and prevent it from entry^15,24,27^. As expected, neurons expressing GluA1-WT and GluA2 showed no detectable calcium responses. Unexpectedly, neurons co-transfected with GluA1 A636T with GluA2 still elicited large calcium transients (Fig. 4b & d), comparable to GluA1-A636T homomers. Thus, A636T confers dominant calcium permeability, overriding the normal inhibitory effect of GluA2 Q/R editing on AMPAR calcium regulation (Fig. 4b, d, e).

Since sustained calcium entry is a well-documented trigger of neuronal excitotoxicity^16,24,25^, we investigated whether the *Gria1*-A636T promotes cell death under excitatory stress. Cultured hippocampal neurons expressing WT or A636T GluA1, with or without GluA2, were challenged with 0.5 μM AMPA for 16 hours. Neuronal viability was assessed by longitudinal imaging of Venus-labeled neurons before and after AMPA treatment. Neurons that maintained intact morphology and fluorescence were classified as viable, whereas loss of Venus signal or neurite fragmentation indicated cell death. Under these conditions, neurons expressing GluA1-WT (alone or with GluA2) remained largely healthy, whereas neurons expressing GluA1-A636T (alone or with GluA2) showed extensive degeneration or loss of Venus fluorescence (Fig. 4f, g). Notably, NBQX fully prevented this cell death, identifying AMPAR overactivation as the direct cause of excitotoxicity, independent of GluA2 expression (Fig. 4f, g).

### *Gria1*-A636T mutation induces hippocampal neuronal death and neuroinflammation

Building on the persistent CP-AMPAR activity in Mut mice and the heightened excitotoxicity in GluA1-A636T-expressing neurons, we next examined whether this mutation compromises neuronal viability *in vivo*. To directly assess cell death, we performed TUNEL assays in juvenile mice to detect DNA fragmentation. Mut mice exhibited a pronounced increase in TUNEL-positive cells compared to WT littermates. Cell death was predominantly confined to the hippocampus, with prominent labeling in CA1 and CA3, a trend toward an increase in the DG, and minimal labeling in adjacent cortical region (Fig. 4h, k). These findings reveal a selective vulnerability of the hippocampus to A636T-induced excitotoxic degeneration.

Accompanying cell death, we also observed robust neuroinflammation in the hippocampal regions. Prominent astrogliosis, marked by an increased GFAP staining (Fig. 4i, l), and microglial activation, marked by increased Iba1 staining and amoeboid morphology (Fig. 4j, m), were observed in CA1 and CA3, with modest increases in the DG, and negligible labeling in the cortex. These glial responses likely represent reactive processes in response to neuronal injury, mediating debris clearance and modulating the inflammatory environment^28,29^. Together, these findings suggest that persistent CP-AMPAR activity and heightened excitatory drive render CA1 and CA3 neurons particularly vulnerable to excitotoxic degeneration.

Developmental analysis showed that the TUNEL labeling during the first postnatal week was comparable between genotypes. Increased cell death first emerged in CA1 at P14 and subsequently in CA3 by P21 (Extended Data Fig. 7). This temporal progression aligns with the sustained CP-AMPAR activity and synaptic hyperexcitability between P7 and P21 in Mut neurons. Moreover, elevated cell death persisted into adulthood, consistent with the progressive hippocampal atrophy. These findings support a mechanistic link between dysregulated AMPAR calcium permeability and progressive hippocampal degeneration.

### ASO therapy rescues neuronal loss and behavioral deficits in *Gria1*-A636T mice

The developmental onset of hippocampal cell death and neuroinflammation in *Gria1-*A636T mice reveals a critical vulnerability driven by AMPAR-mediated hyperexcitability that worsens with age. To explore potential therapeutic interventions, we first evaluated pharmacological approaches aimed at suppressing AMPAR activity. Specifically, we assessed perampanel (PER), a FDA-approved AMPAR antagonist^30^, and JNJ-5551118 (JNJ), a selective inhibitor targeting hippocampus-enriched TARPγ8-bound AMPARs^31^. Both treatments significantly reduced the number of TUNEL-positive cells compared to vehicle-treated Mut mice. However, cell death remained significantly elevated relative to WT controls (Extended Data Fig. 8a, b, c, e). Moreover, neither treatment restored the thickness of the CA1 pyramidal cell layer, which remained significantly reduced in treated Mut mice (Extended Data Fig. 8d, f), which is consistent with the persistent loss of pyramidal neurons. Notably, PER at an effective dose produced severe motor side effects. These results highlight the limitations of general AMPAR antagonists and the need for mutation-specific precision therapies.

To achieve selective suppression of mutant receptor expression, we developed an antisense oligonucleotide (ASO) therapy targeting the *Gria1*-A636T transcript. ASOs are short synthetic nucleic acids that bind to complementary mRNA sequences, enabling allele-specific transcript degradation. This technology has demonstrated therapeutic efficacy and is FDA-approved for multiple neurological disorders^12,13^. Gapmer-type ASOs were delivered by intracerebroventricular (ICV) injection into *Gria1* pups at the early neonatal stage (P2-P3), and their efficacy and specificity were evaluated in hippocampal tissues collected at the juvenile stage (Fig. 5a). Allele-specific RT-qPCR reliability distinguished between WT and Mut *Gria1* transcripts, and accurately detected the expected WT and Mut mRNA levels in the WT or Mut *Gria1* mice (Fig. 5b, c). Among the candidates tested, ASO2 produced the most potent and selective knockdown of Mut mRNA, reducing mutant transcript levels to ∼40% of vehicle-treated controls without affecting WT expression (Fig. 5d, e). However, ASO2 administration was associated with high mortality (< 50% survival). Incorporating cytosine methylation at CpG motifs in ASO2 (ASO2^mC^), a modification known to reduce ASO toxicity^14,32^, not only restored the survival to control levels but also further reduced mutant transcripts to ∼26% of the control (Fig. 5d).

**Fig. 5.**
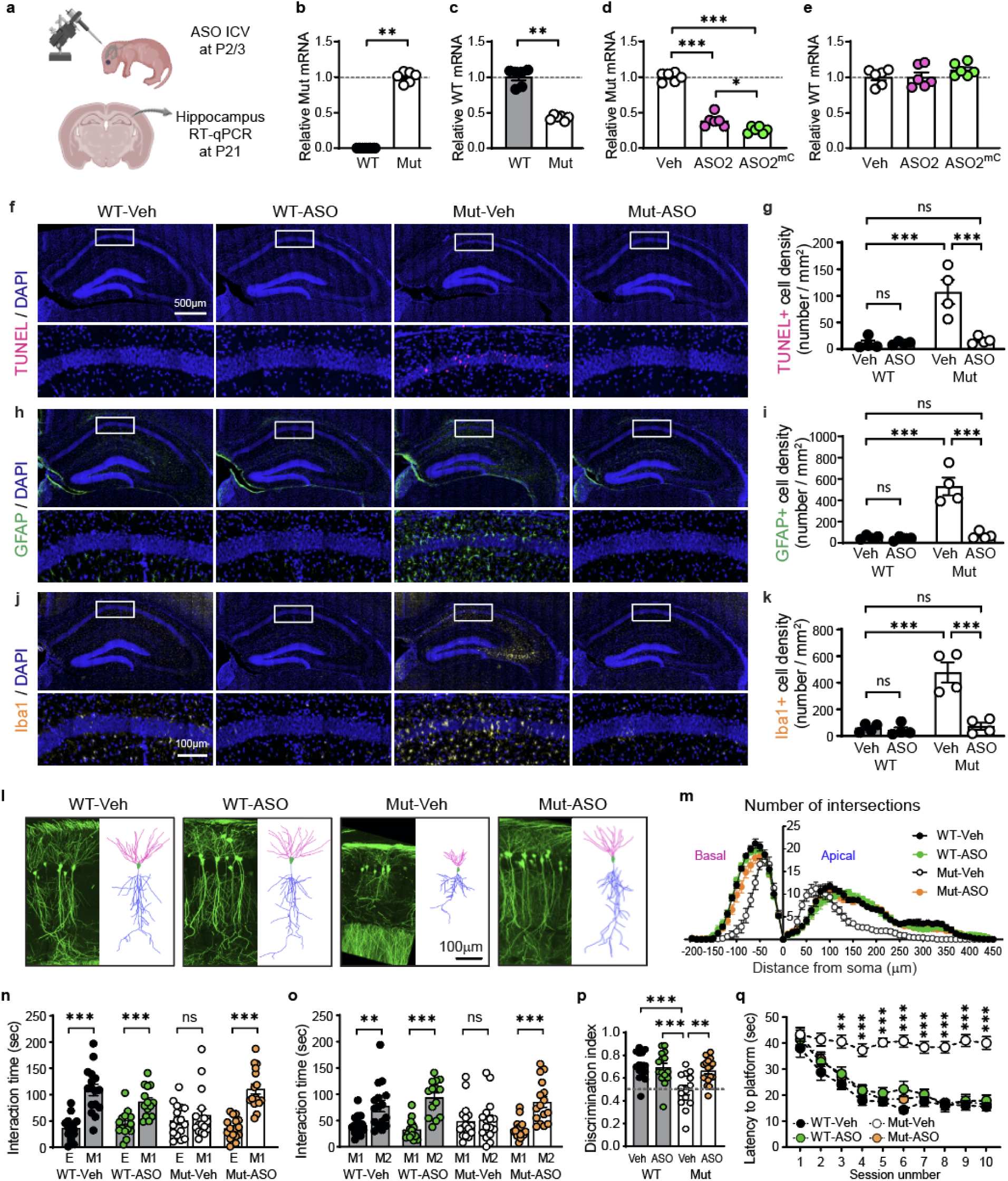
Allele-specific ASO therapy rescues hippocampal pathology and behavioral deficits in *Gria1* Mut mice. **a,** Schematic of ASO administration and efficacy assessment following intracerebroventricular (ICV) injection. **b-e,** Allele-specific RT-qPCR analysis of hippocampal tissue at P21. **b,** Mutant transcripts were detected exclusively in Mut mice (P=0.002). **c,** WT transcript levels were reduced by half in Mut mice (P=0.002). **d,** ASO2 and ASO2^mC^ significantly reduced mutant transcript levels compared to vehicle (Veh) controls (Veh vs. ASO2, P < 0.0001; Veh vs. ASO2^mC^, P < 0.0001; ASO2 vs. ASO2^mC^, P = 0.016). **e,** WT transcript levels remained unchanged across treatments. Dashed lines indicate normalized level (=1). **f-g,** Representative TUNEL staining (whole hippocampus and magnified CA1) and quantification of TUNEL+ cell density in CA1 showing that ASO treatment significantly rescued neuronal survival (Mut-Veh vs. Mut-ASO, P=0.0009), and reduced cell death to WT levels (Mut-ASO vs. WT-Veh, P>0.0000). **h-k,** Representative GFAP (**h**) and Iba1 (**j**) immunostaining and quantifications (**i, k**) showing that ASO treatment markedly rescued neuroinflammation in the CA1 (Mut-Veh vs. Mut-ASO: GFPA, P < 0.0001; Iba1, P=0.0001). **l-m,** Representative images and Sholl analysis of GFP-labeled CA1 pyramidal neurons showing that ASO treatment rescued dendritic arbor complexity in Mut mice. Significant treatment effects were observed across multiple distances from the soma (see Supplementary table for full statistics). **n-q,** Behavioral recovery following neonatal ASO therapy. ASO-treated Mut mice showed normalized sociability (**n**) and social novelty preference (**o**) in the three-chamber test, improved spatial memory in the Y-maze (**p**) and enhanced learning performance in the MWM (**q**). The dashed line in (**p**) indicates chance. (Statistical details are provided in Supplementary table). Data are presented as mean ± s.e.m. *P < 0.05, **P < 0.01, ***P < 0.001. Exact P values, sample sizes, and statistical tests are provided in the Supplementary Table.

Remarkably, a single neonatal administration of ASO2 or ASO2^mC^ produced a robust rescue of hippocampal pathology. TUNEL-positive cells (Fig. 5f, g), and neuroinflammatory markers, including GFAP (Fig. 5h, i) and Iba1 (Fig. 5j, k), were markedly reduced at the juvenile stage. At the cellular level, Sholl analysis showed that the dendritic architecture of pyramidal neurons was also fully restored (Fig. 5l, m). Notably, ASO-treated WT mice showed no evidence of neuronal toxicity or inflammation (no increase in TUNEL, GFAP or Iba1 signals, Fig. 5f-k), confirming the therapeutic specificity and safety of our ASO therapy. Thus, a single neonatal injection provided sustained protection for at least three weeks, demonstrating long-lasting efficacy against excitotoxic degeneration *in vivo*.

To determine whether ASO therapy also ameliorates behavioral deficits, we assessed the social and cognitive performance in *Gria1* mice following neonatal treatment. In the three-chamber test, vehicle-treated Mut mice failed to show normal sociability or social novelty preference, whereas ASO-treated Mut mice displayed significant restoration of both social behaviors (Fig. 5n, o). ASO treatment also rescues additional ASD-associated behaviors, including reciprocal social interactions, grooming, and anxiety (Extended Data Fig. 9-c). Moreover, ASO treatment restored discrimination index in the Y-maze, indicating recovery of short-term spatial memory (Fig. 5p). In the MWM learning test, vehicle-treated Mut mice exhibited prolonged latency to find the hidden platform across training sessions, whereas ASO-treated Mut mice performed comparably to WT controls, reflecting the restoration of spatial learning (Fig. 5q). Therefore, early allele-specific ASO intervention not only prevents hippocampal pathology but also restores social and cognitive function in this NDD mouse model.

In summary, ASO-mediated knockdown of the mutant *Gria1* provides a precise and effective therapeutic strategy that far exceeds general AMPAR antagonism. These findings establish a direct mechanistic link between AMPAR dysregulation and excitotoxic vulnerability in *Gria1-*A636T mice and highlight precision RNA-targeted therapy as a promising strategy for treating mutation-associated neurodevelopmental and neurodegenerative disorders.

## Discussion

Neurodevelopmental disorders often cause lifelong impairments, yet effective therapies remain limited. Although recent genetic studies implicate *de novo* mutations in glutamate receptor genes as major contributors to NDDs^6,33^, their pathogenic mechanisms and therapeutic strategies remain largely unexplored. Here, we establish a direct causal link between a clinically identified *Gria1*-A636T mutation and ASD/ID-like behavioral deficits in mice. This mutation confers a selective vulnerability of the hippocampus, characterized by severe dendritic atrophy, neuronal death, and neuroinflammation, driven by heightened excitatory synaptic transmission and aberrant calcium permeability. Notably, allele-specific ASO therapy fully rescued both hippocampal pathology and behavioral deficits, providing strong preclinical support for precision RNA-targeted interventions in mutation-associated NDDs.

Despite the broad distribution of AMPARs throughout the CNS, *Gria1-*A636T mice exhibited a more pronounced reduction in GluA1 (∼50%) and associated proteins in the hippocampus compared to other brain regions, likely due to the high expression of GluA1 in this region^5,34^. Interestingly, *Gria1* knockout mice, which also lack GluA1, exhibit relatively mild phenotypes, including preserved hippocampal structure, normal calcium transients, comparable synaptic ultrastructure and largely intact learning and memory^34,35^. In contrast, *Gria1-*A636T mice exhibit a pronounced gain-of-function phenotype, marked by elevated excitatory drive and persistent CP-AMPAR activity, resulting in progressive neuronal degeneration and ASD/ID-like behavioral deficits. Supporting a dominant gain-of-function mechanism, cultured neurons expressing GluA1-A636T showed large NBQX-sensitive calcium transients and underwent AMPAR-dependent neuronal death. These effects persisted even with GluA2 co-expression, indicating GluA1-A636T functions as a dominant effector of calcium permeability despite the normally protective GluA2 Q/R editing. Thus, the pathology arises not from the loss of GluA1 per se, but from the aberrant channel function.

Although the *Gria1*-A636T mutation is homologous to the *Griδ2*-Lurcher mutation, their pathogenic mechanisms are fundamentally distinct. The Lurcher mutation causes constitutive GluD2 channel opening, leading to large sodium leak and chronic depolarization even at the resting state^9^. In contrast, *Gria1*-A636T-containing AMPARs remain ligand-gated but exhibit enhanced calcium permeability and excitatory drive without basal leak currents or resting depolarization. This difference results in activity-dependent, rather than constitutive, excitotoxic stress, highlighting the mechanistic novelty of the *Gria1*-A636T mutation.

The A636 residue of GluA1 is located within the conserved “SYTANLAAF” gating motif in the M3 transmembrane helix, adjacent to the ligand-binding domain (LBD). Upon glutamate binding, closure of the LBD induces conformational changes that relieve gating constraints and allow the channel to open^36^. Structural modeling suggests that A636 normally stabilizes hydrophobic interactions with adjacent subunits, particularly residues Leu638 and Thr639, to maintain the receptor in a closed or desensitized state^7^. The A636T mutation likely weakens these interactions, favoring channel opening and delaying desensitization. Moreover, the A636T mutation may also introduce a new hydrogen bond with neighboring N633, generating an outward kink in the M3 helix and localized dilation of the upper gate. This rearrangement may enhance the coupling between ligand binding and gate opening, thereby lowering the activation threshold^37^. Furthermore, a novel calcium-binding site within the SYTANLAAF motif has recently been identified^38,39^, and the mutation at the residue homologous to N633 in GluA1 has been shown to disrupt calcium binding and permeability. The T636-N633 interaction may affect calcium binding and contribute to the increased calcium permeability in *Gria1-*A636T AMPARs. These structural predictions align well with our electrophysiological findings, including increased mESPC charge transfer, elevated event frequency, and persistent calcium permeability. Together, these data support a model in which the A636T mutation causes aberrant AMPAR gating and calcium permeability, leading to excitotoxicity and neuronal degeneration.

Our longitudinal analysis revealed a critical developmental window during which pathology emerges in *Gria1-*A636T mice. At P7, electrophysiological alterations were subtle, with no detectable differences in AMPAR-mediated mEPSC amplitude, AMPAR/NMDAR ratio, or inward rectification, and neuronal viability remained largely intact. By P14-21, coinciding with the normal developmental reduction in AMPAR calcium permeability^16–18^, *Gria1*-A636T mice exhibited persistent AMPAR calcium permeability, enhanced excitatory transmission, and the onset of neuronal loss. Since dysregulated CP-AMPAR has been implicated in multiple neurodevelopmental and neurodegenerative disorders^16,27,40^, these findings suggest that persistent calcium permeability driven by altered A636T-containing AMPAR gating is a primary contributor to excitotoxic degeneration in this NDD model.

Therapeutically, general AMPAR antagonists such as perampanel and JNJ-5551118 showed limited efficacy despite broad inhibition of excitatory transmission. In contrast, allele-specific ASO therapy targeting the mutant *Gria1* transcript provided robust and sustained rescue of hippocampal structure, neuronal survival, and animal behaviors following a single neonatal injection. Given the evolutionary conservation of the transmembrane domain 3, ASO-based precision therapies may have broader application to glutamate receptor-related disorders. This includes homologous mutations at the conserved SYTANLAAF motif, such as *GRID2-*A654T^41^ and *GRIK2*-A657T^42^, which correspond to the A636T position in *GRIA1*. Additionally, several functionally analogous mutations in *GRIA2,* including the A639S, T646N and V647L variants, have been shown to alter gating or calcium permeability and are associated with various neurodevelopmental phenotypes^33^. These parallels support the broader translational potential of allele-specific RNA-targeted therapies for glutamate receptor channelopathies.

Our ASO screen was performed in a CRISPR/Cas9 edited mouse model with additional silent mutations introduced to prevent Cas9 from re-cutting and to facilitate screening. Further studies using humanized *Gria1-*A636T mice or patient-derived induced pluripotent stem cells (iPSCs) will be essential to validate this strategy. Ultimately, the development of human-compatible ASOs and improved delivery strategies will be critical next steps towards clinical translation.

Taken together, our study establishes the *Gria1*-A636T variant as a pathogenic gain-of-function mutation that disrupts the normal developmental regulation of AMPAR calcium permeability, leading to hippocampal excitotoxicity, circuit degeneration, and ASD/ID-like behaviors during postnatal maturation. We identify a distinct developmental excitotoxicity, in which sustained CP-AMPAR activity arises from a dominant effect of the A636T mutation on channel permeability, rather than the constitutive channel opening characteristic of canonical models, such as the *Griδ2*-Lurcher mutation. Importantly, allele-specific silencing of the mutant transcript fully prevents neuropathology and normalizes behavior, demonstrating that precise molecular correction can overcome pathogenic AMPAR hyperactivity *in vivo*. Together, these findings define a new framework for glutamate receptor-associated channelopathies and highlight precision RNA-targeted therapies as a promising and generalizable strategy for *GRIA1*-related disorders and other neurodevelopmental and neurodegenerative diseases driven by similar excitotoxic mechanisms.

## Supporting information

Suppl. Table_for statistics

## Acknowledgments

We thank Drs. Hollis Cline, Ingie Hong, Qianwen Zhu, Dylan Hale, and Sin-Jhong Cheng for their expert technical assistance and valuable scientific discussions. We also thank Ting-Wei Lai, Ya-Hui Yang, Shu-Ching Shih, Yebeen Kim and Yi-An Lai for their assistance with imaging analysis, as well as Sarah Richardson, Ashley Irving, and Wei-Yu Hsu for their help with mouse care. Additionally, we are grateful to Lisa Hann for administrative support. We acknowledge the Laboratory Animal Facility and Imaging Core Facility of the Institute of Cellular and Organismic Biology (ICOB), Academia Sinica, for mouse care and imaging support; the DNA Sequencing Core Facility (AS-CFII-113-A12) and Transgenic Core Facility of Academia Sinica for sequencing and consultation on genetic tools; and the Animal Image Facility, BioTReC, Academia Sinica, and the Taiwan Animal Consortium for MRI acquisition and technical support with imaging analysis. The schematic in Fig. 5a was created using BioRender.com.

## Author contributions

Conceptualization: RLH, SLC

Methodology: CMC, YMH, CCC, RCJ

Investigation: CMC, YMH, CCC, YHC, HLT, CYT, FYH

Visualization: CMC, YMH, CCC

Funding acquisition: RLH, SLC

Project administration: CMC, YMH

Supervision: CMC, RLH, SLC

Writing – original draft: CMC, YMH, CCC, SLC

Writing – review & editing: RHL, SLC

## Data and materials availability

All data are available in the main text or the supplementary materials.

Supplementary Table **(Separate Excel file)**

---

**Extended Data Fig. 1.**
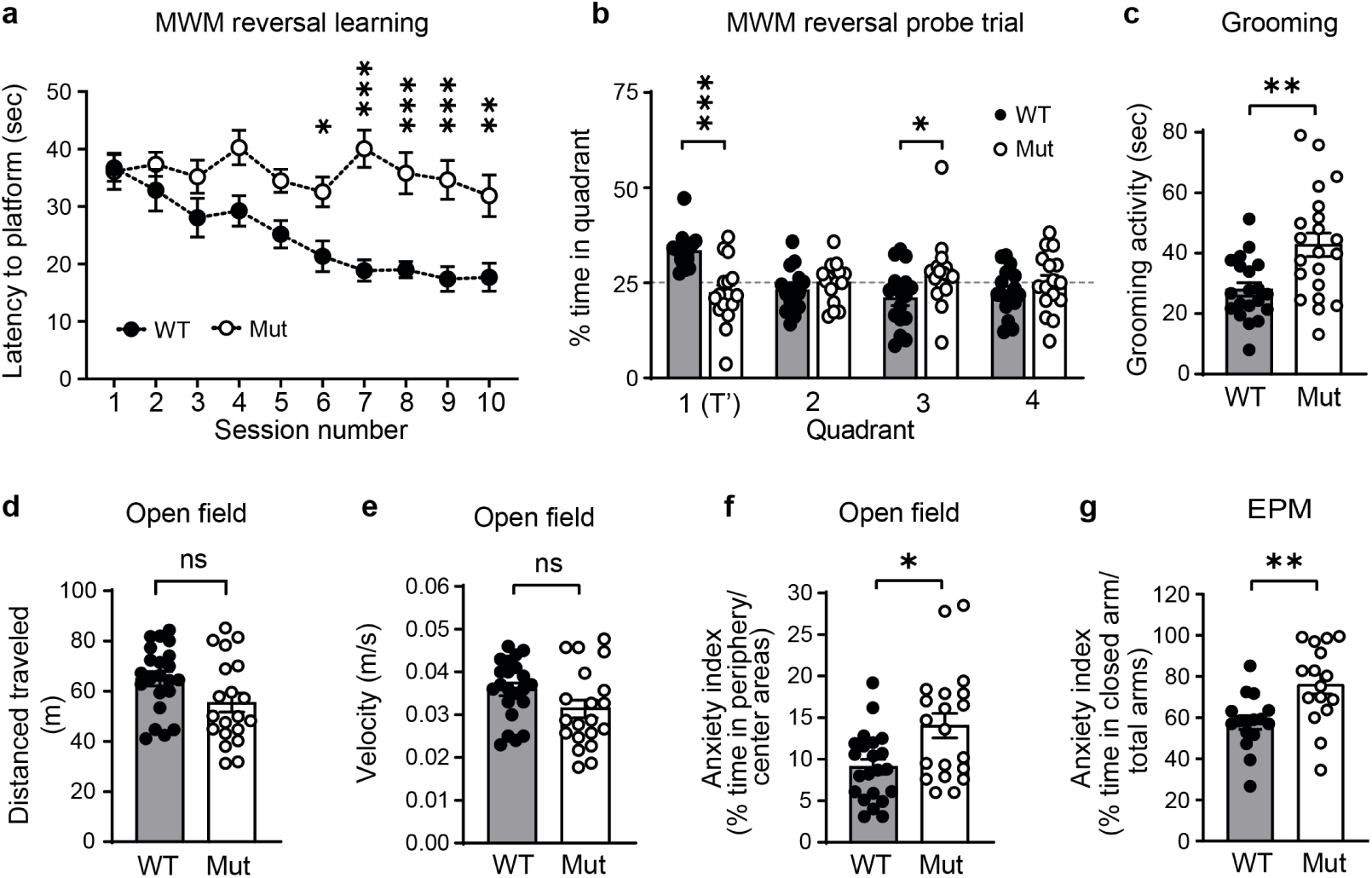
Additional behavioral characterization of *Gria1* Mut mice. **a**, Reversal learning performance in the Morris water maze (MWM), quantified as latency to locate the repositioned target platform (T’) in quadrant 1 across sessions. **b**, Time spent in each quadrant during the reversal probe trial. The dashed line indicates chance level. **c**, Self-grooming time was significantly increased in Mut mice, indicating increased stereotypic behavior (P=0.0039). **d**, **e**, Locomotor activity, including total distance traveled (**d**) and averaged velocity (**e**), test showed no significant genotype differences. **f**, The anxiety index in the open field test, calculated as the percentage of time spent in the periphery versus the center areas, was elevated in Mut mice (P=0.0159). **g**, The anxiety index in the elevated plus maze (EPM), calculated as the percentage of time spent in the closed arms, was increased in the Mut mice (P=0.0022). Data are presented as mean ± s.e.m. *P < 0.05, **P < 0.01, ***P < 0.001. Exact P values, sample sizes, and statistical tests are provided in the Supplementary Table.

**Extended Data Fig. 2.**
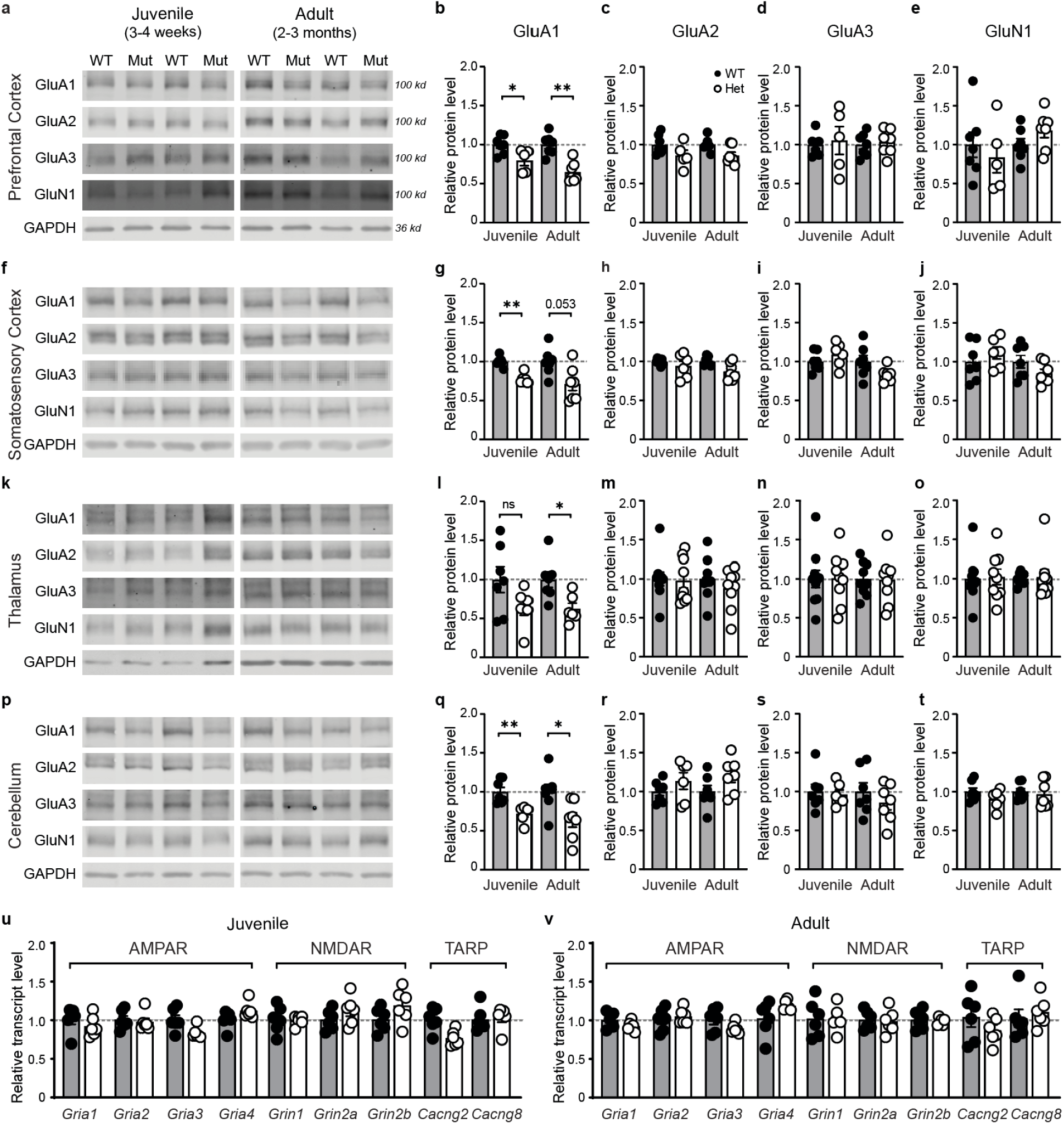
Protein and transcript expression of AMPAR and NMDAR subunits in *Gria1* WT and Mut brains. **a-t**, Representative Western blots and quantification of AMPAR and NMDAR subunit protein levels in the prefrontal cortex (**a-e**), somatosensory cortex (**f-j**), thalamus (**k-o**), and cerebellum (**p-t**). Among the proteins analyzed, GluA1 expression was significantly reduced in Mut mice, whereas no significant differences were observed for the remaining subunits across all examined regions. **u**, **v**, RT-qPCR analysis of AMPAR (*Gria1-4*), NMDAR subunits (*Grin1, 2a, 2b*) and AMPAR axillary proteins (*Cacng2* and *Cacng8*) transcripts in the hippocampus showed no significant genotype differences. Dashed lines indicate normalized WT levels (=1). Data are presented as mean ± s.e.m. *P < 0.05, **P < 0.01, ***P < 0.001. Exact P values, sample sizes, and statistical tests are provided in the Supplementary Table.

**Extended Data Fig. 3.**
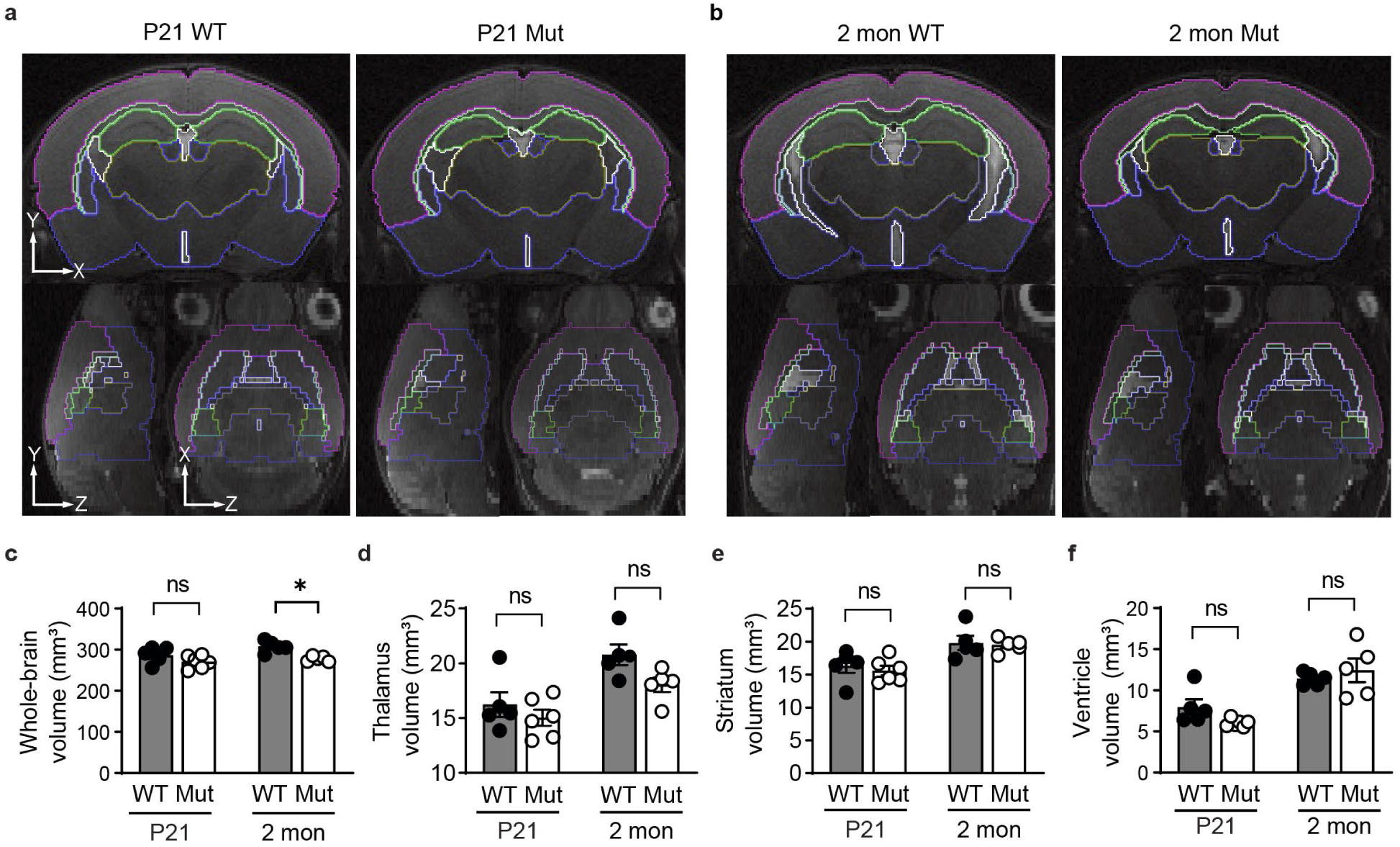
Whole-brain and regional volumetric analysis by MRI. **a**-**b**, Representative coronal, sagittal, and axial MRI images with segmented brain regions outlined in *Gria1* WT and Mut mice at P21 and 2 months. **c-f**, Quantification of whole-brain (**c**), thalamus (**d**), striatum (**e**), and ventricular (**f**) volumes. No significant regional differences were detected, except for a modest but significant reduction in whole-brain volume in Mut mice at 2 months (p=0.035). Data are presented as mean ± s.e.m. *P < 0.05, **P < 0.01, ***P < 0.001. Exact P values, sample sizes, and statistical tests are provided in the Supplementary Table.

**Extended Data Fig. 4.**
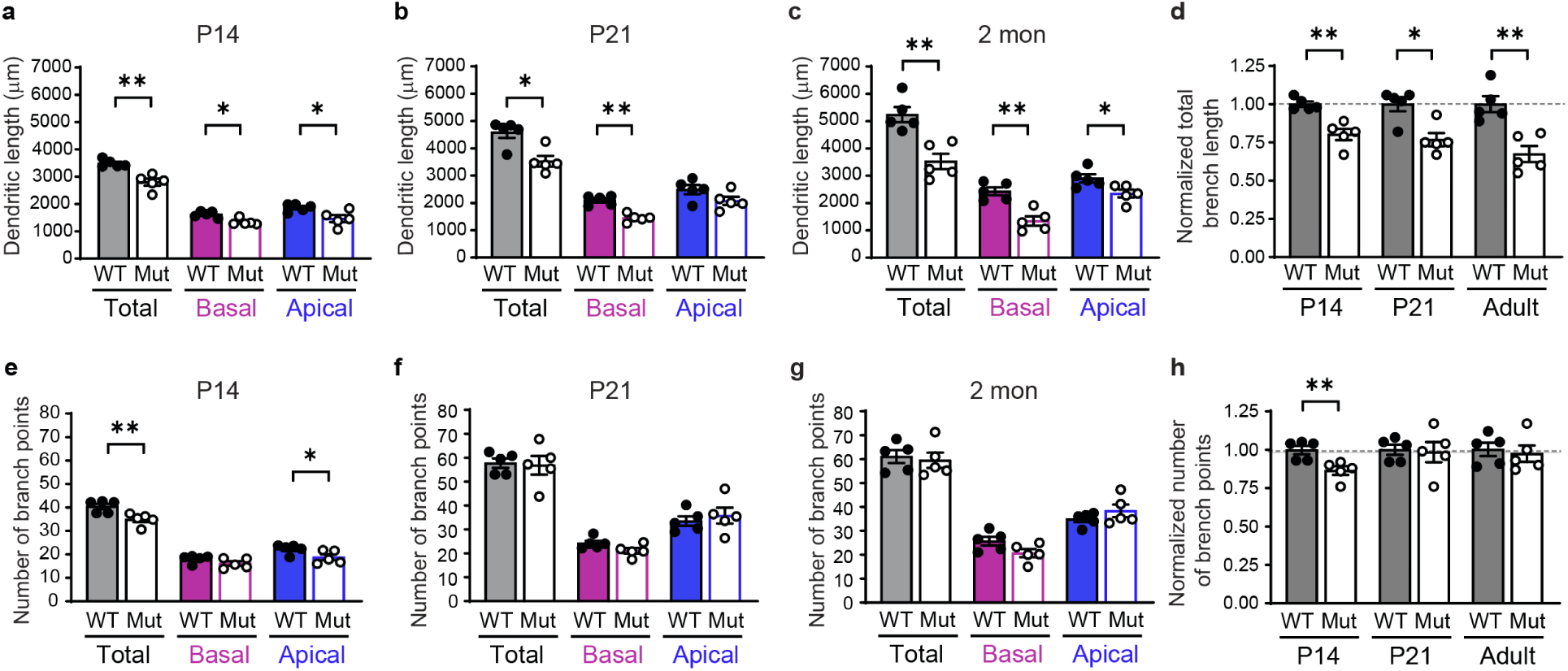
Additional analysis of CA1 dendritic morphology across developmental stages. **a-c**, Total dendritic length of GFP-labeled CA1 pyramidal neurons at P14 (**a**), P21 (**b**), and 2 months (**c**). Mut mice exhibited significantly reduced dendritic length at all ages. **d**, Relative dendritic branch length of CA1 pyramidal neurons compared to WT littermates at P14, P21, and adulthood. Mut mice exhibited a progressive reduction in total dendritic length, with a 20% decrease at P14, 23% at P21, and 33% at adulthood (all P<0.05). **e-g**, Quantification of dendritic branch points in GFP-labeled CA1 pyramidal neurons at P14 (**e**), P21 (**f**), and 2 months (**g**). **h,** Relative number of dendritic branch points compared to WT littermates at P14, P21, and adulthood. No significant differences were observed at P21 or adulthood except for the reduction detected at P14 (P=0.0079). Dashed lines indicate normalized WT levels at each age. Data are presented as mean ± s.e.m. *P < 0.05, **P < 0.01, ***P < 0.001. Exact P values, sample sizes, and statistical tests are provided in the Supplementary Table.

**Extended Data Fig. 5.**
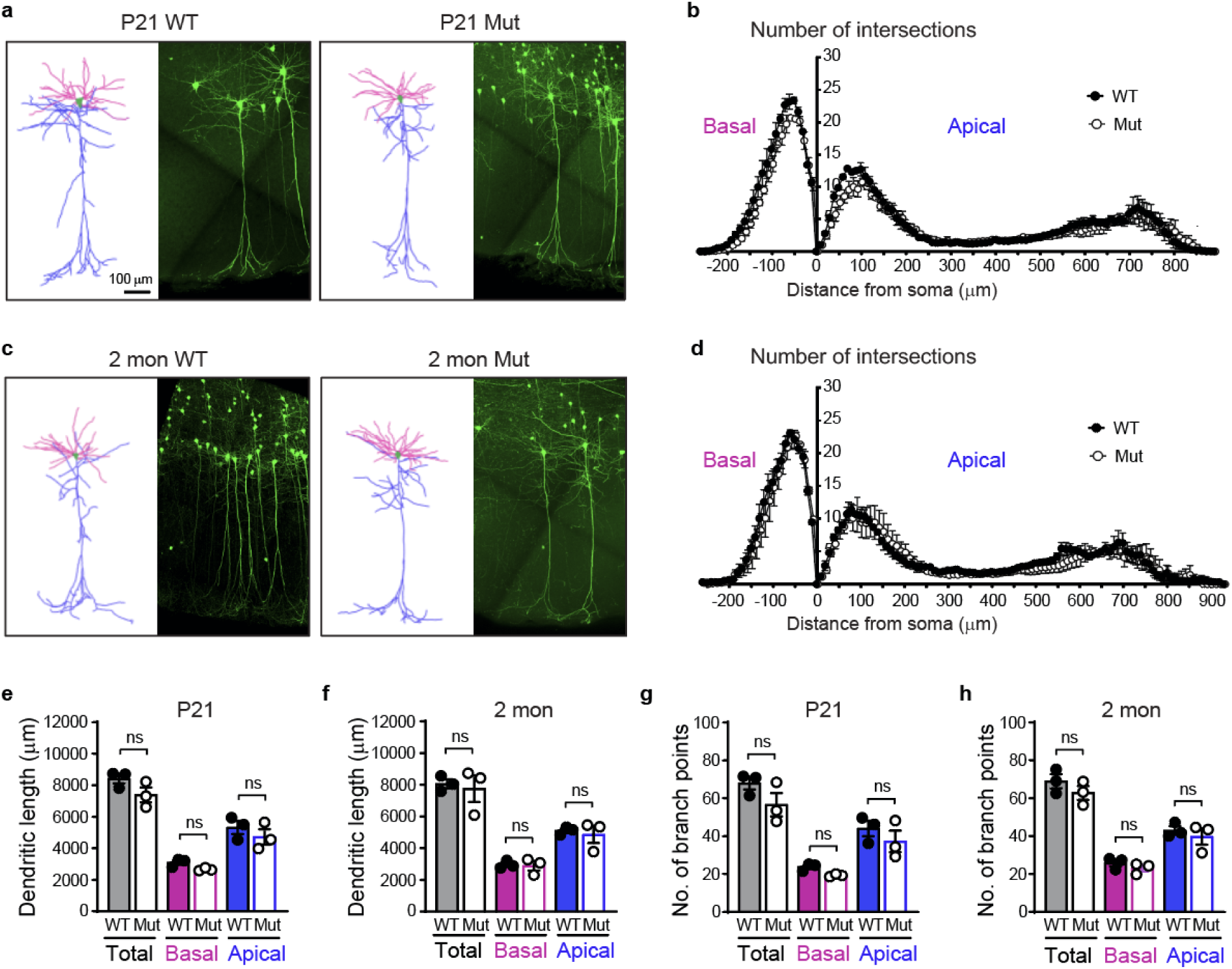
Preserved dendritic architecture in the somatosensory cortex of *Gria1* Mut Mice. **a**, Representative images of GFP-labeled L2/3 pyramidal neurons from juvenile WT and Mut mice. **b**, Sholl analysis of dendritic architecture of cortical neurons from juvenile mice showed no significant genotype differences. **c**, Representative images of GFP-labeled Layer 2/3 pyramidal neurons in adult WT and Mut mice. **d**, Sholl analysis of dendritic architecture of cortical neurons from adult mice showed no significant genotype differences. **e**, **f**, Quantification of total dendrite length in apical and basal dendrites of juvenile (**e**) and adult (**f**) mice showed no significant genotype differences. **g**, **h**, Quantification of dendritic branch points in apical and basal dendrites of juvenile (**g**) and adult (**h**) mice showed no significant genotype differences. Data are presented as mean ± s.e.m. *P < 0.05, **P < 0.01, ***P < 0.001. Exact P values, sample sizes, and statistical tests are provided in the Supplementary Table.

**Extended Data Fig. 6.**
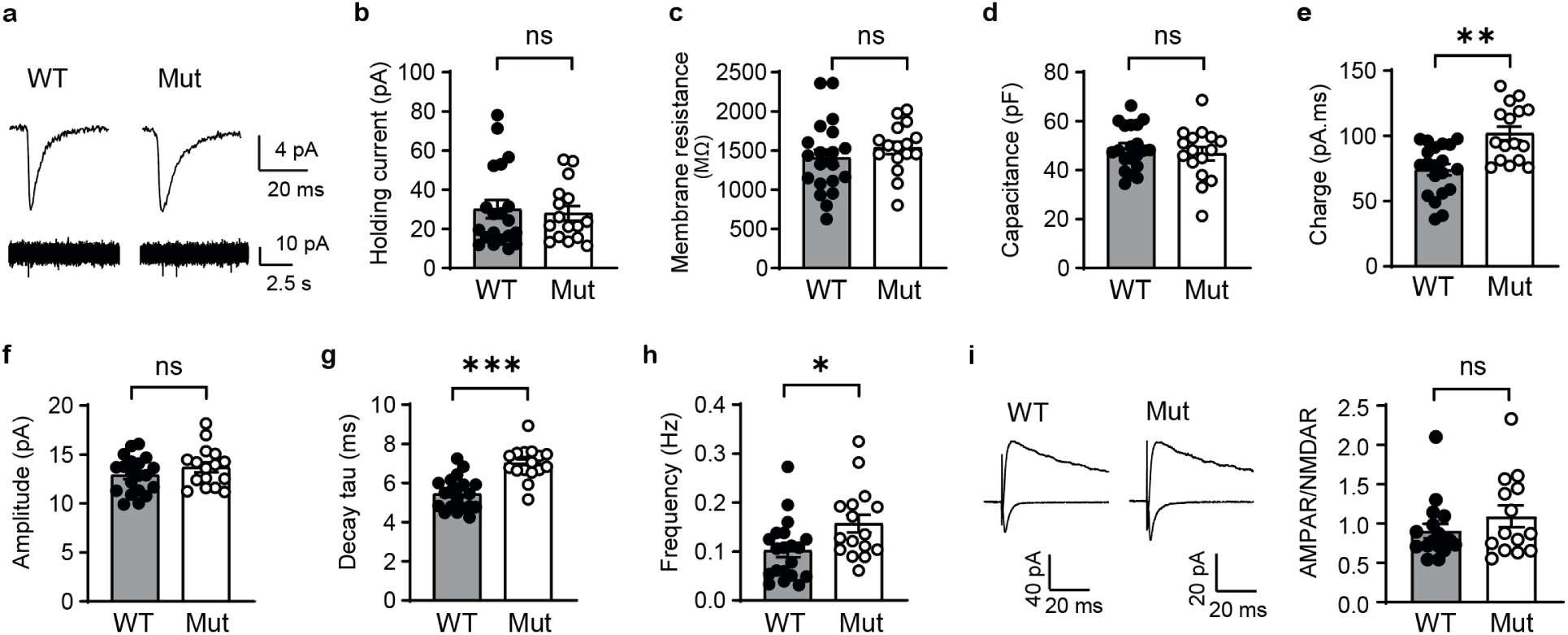
Electrophysiological properties of neonatal CA1 neurons from *Gria1* WT and Mut mice. **a**, Representative AMPAR-mEPSC traces recorded from hippocampal CA1 neurons at P5-9. **b-d**, Basal membrane properties of neonatal Mut CA1 neurons, including holding current (**b**), membrane resistance (**c**), and capacitance (**d**), were comparable between genotypes. **e-h**, AMPAR-mEPSC properties, including charge transfer (**e,** P=0.0015), decay tau (**g,** P<0.0001) and frequency (**h,** p=0.0239) were increased in neonatal Mut neurons, while amplitude (**f,** P=0.3359) showed no genotype difference. **i**, Representative eEPSC traces and quantification of AMPAR/NMDAR ratio showed no genotype difference (p=0.3371). Data are presented as mean ± s.e.m. *P < 0.05, **P < 0.01, ***P < 0.001. Exact P values, sample sizes, and statistical tests are provided in the Supplementary Table.

**Extended Data Fig. 7.**
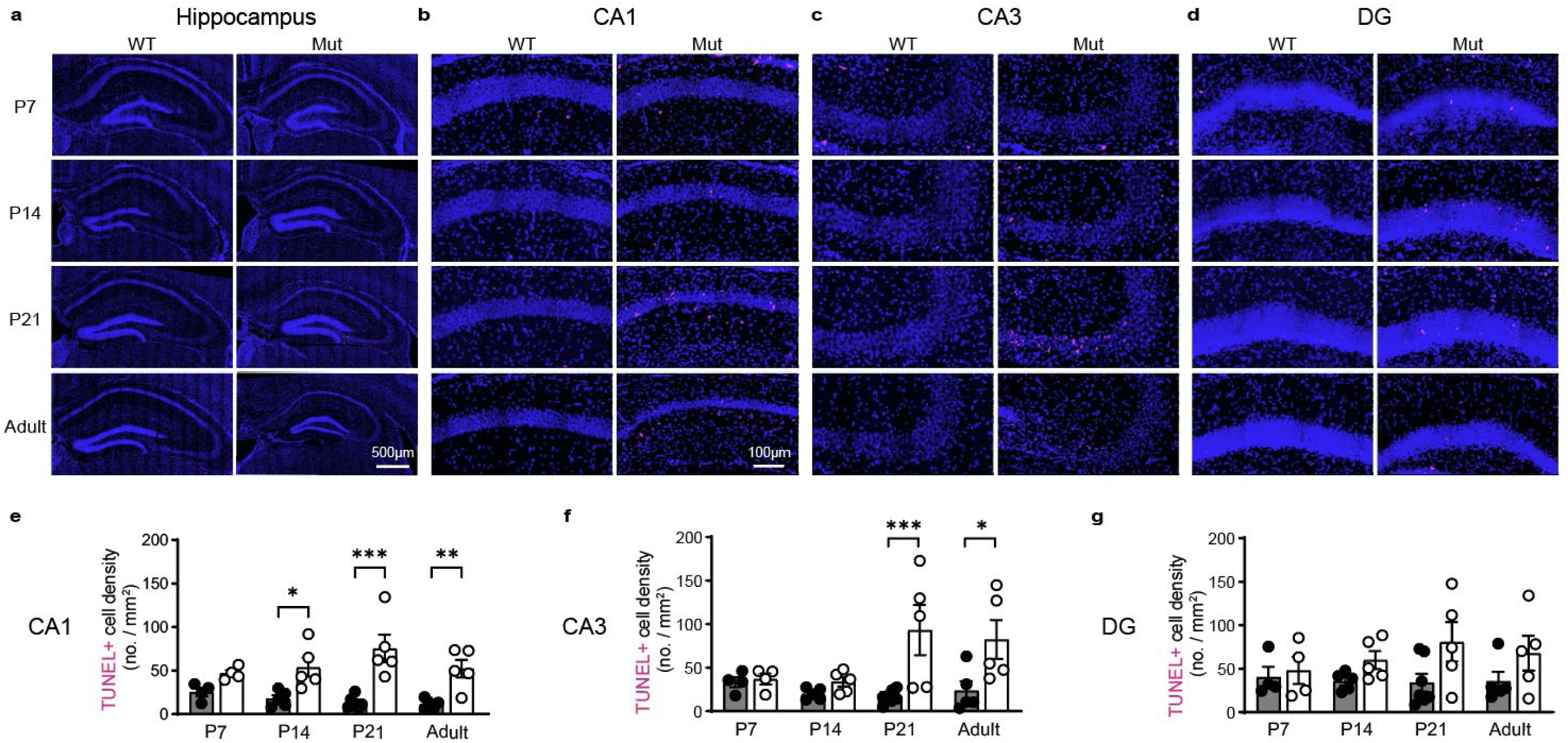
Developmental progression of hippocampal cell death in *Gria1* Mut mice. **a-d**, Representative images of the hippocampus and magnified CA1, CA3 and DG stained for TUNEL (magenta) and DAPI (blue) at P7, P14, P21, and adults (2-3 months)). **e-g,** Quantification of TUNEL+ cell density showing a significant, progressive increase in cell death in CA1 (**e**) and CA3 (**f**) regions, with less pronounced effect in the DG (**g**). Data are presented as mean ± s.e.m. *P < 0.05, **P < 0.01, ***P < 0.001. Exact P values, sample sizes, and statistical tests are provided in the Supplementary Table.

**Extended Data Fig. 8.**
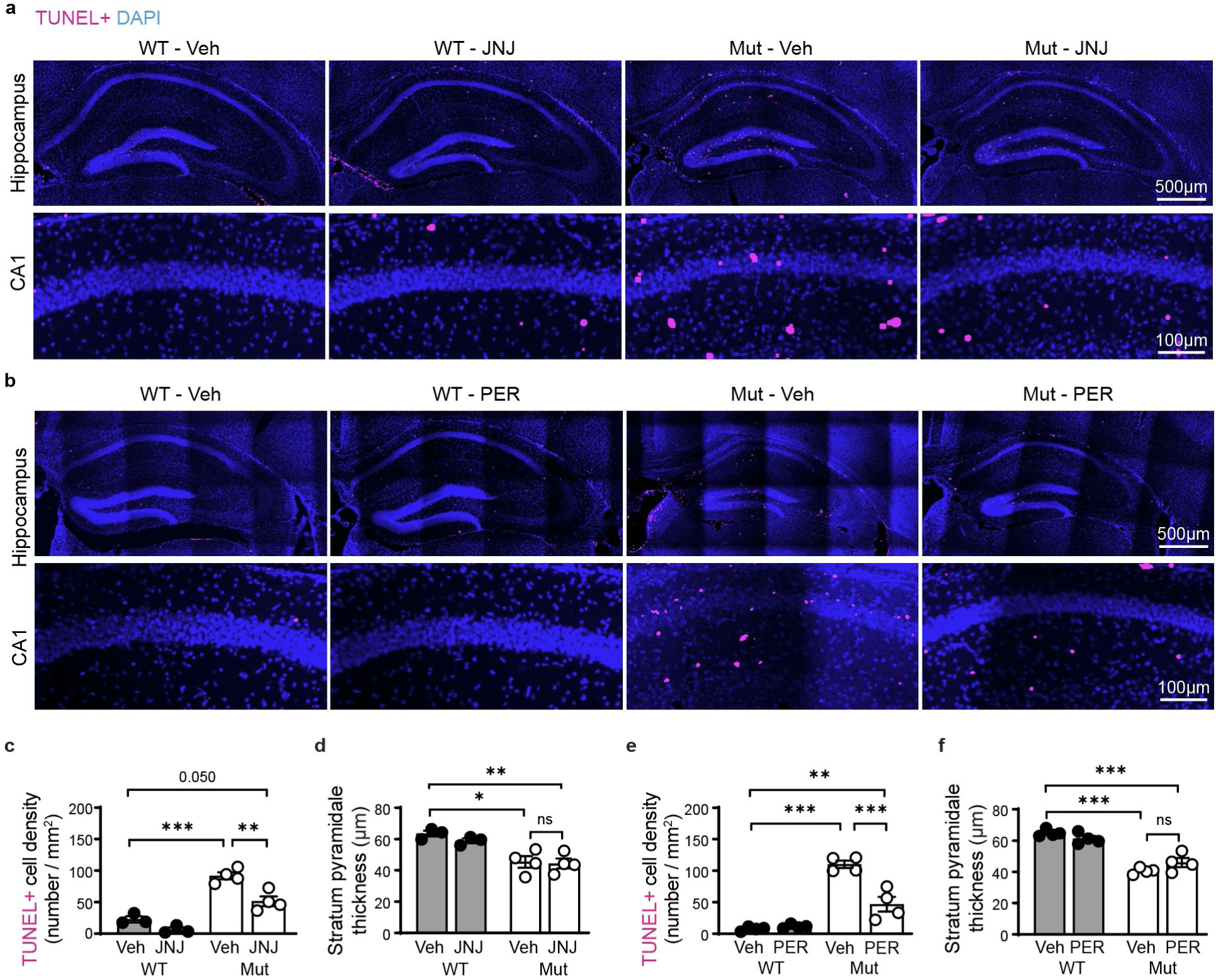
Pharmacological AMPAR blockade partially reduces hippocampal cell death. **a**, Representative TUNEL-stained hippocampus sections and magnified CA1 regions from *Gria1* WT and Mut mice treated with JNJ-5551118 (JNJ) or vehicle (Veh, 0.25% hydroxypropyl methylcellulose) control. **b,** Representative TUNEL-stained hippocampal sections and magnified CA1 regions from WT and Mut mice treated with perampanel (PER) or water Vehicle. **c, e,** Quantification of TUNEL+ cell density in CA1 showing significantly reduced cell death in Mut mice treated with JNJ (**c**, P=0.0033) or PER (**e**, P=0.0001) compared to Veh-treated Mut mice. **d, f,** Quantification of CA1 stratum pyramidale thickness showing a significant reduction in layer thickness in Mut mice compared to WT-Veh controls (**c**, P=0.0016; **e**, p<0.00001), consistent with pyramidal neuron loss. No significant recovery of layer thickness was observed in treated Mut mice compared to Veh-Mut following either JNJ (**d,** P>0.9999) or PER (**f,** P=0.3649) treatment. Data are presented as mean ± s.e.m. *P < 0.05, **P < 0.01, ***P < 0.001. Exact P values, sample sizes, and statistical tests are provided in the Supplementary Table.

**Extended Data Fig. 9.**
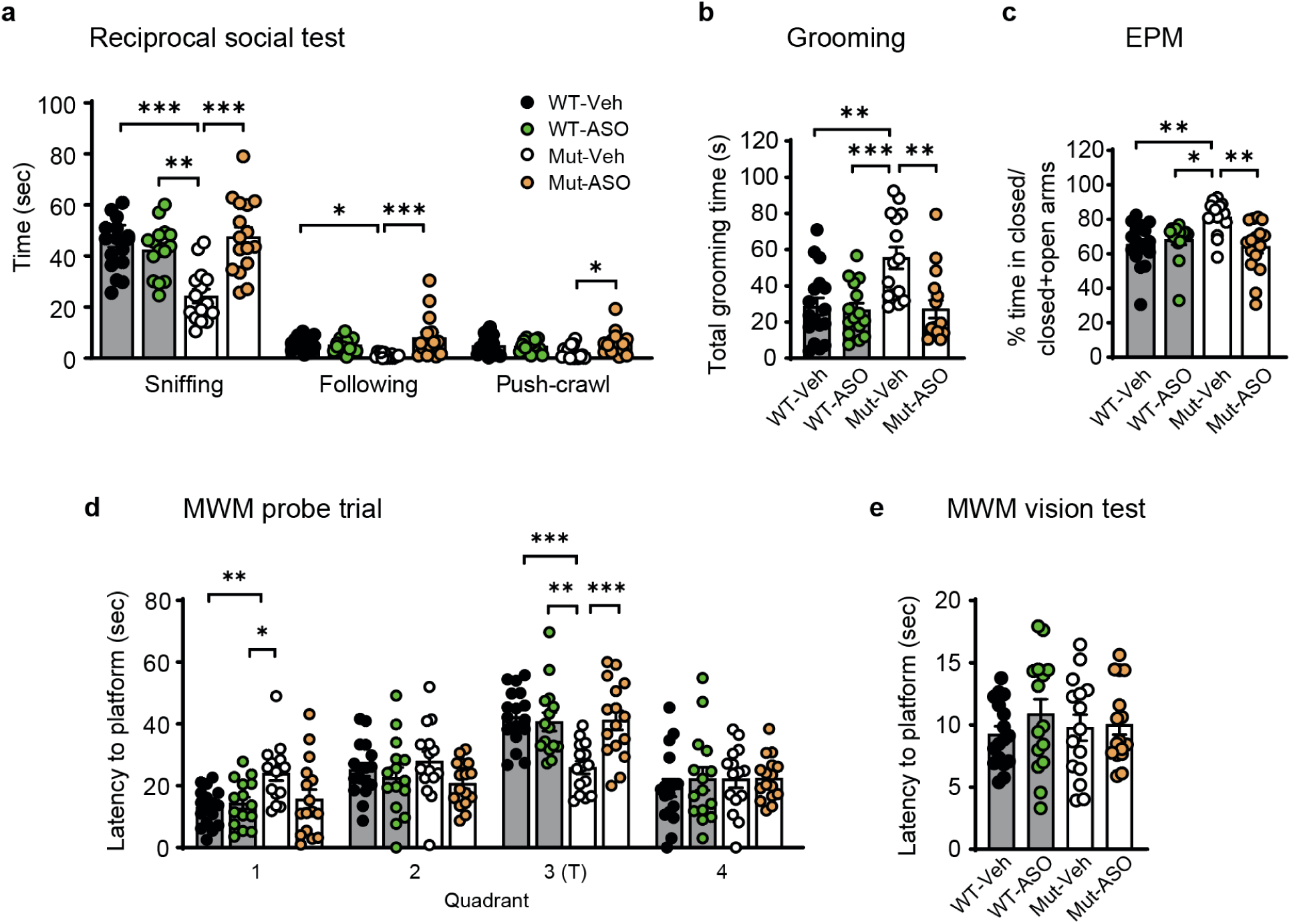
Additional behavioral characterization of ASO-treated *Gria1* mice. **a**, Vehicle-treated Mut mice (Mut-Veh) exhibited reduced social behaviors compared to WT controls. ASO treatment significantly restored social deficits interactions across all measures (Mut-Veh vs Mut-ASO: sniffing, P<0.0001; following, P=0.0004 and push-crawl, P<0.0262). **b**, Increased repetitive self-grooming behavior in Mut-Veh mice was significantly reduced by ASO treatment (Mut-Veh vs Mut-ASO, P=0.0010). **c**, Elevated anxiety-liked behavior in Mut mice was ameliorated by ASO treatment (Mut-Veh vs Mut-ASO, P=0.0016). **d**, in the MWM probe trial, Mut-Veh mice showed impaired spatial memory retention, indicated by reduced time in the target quadrant (T). ASO treatment significantly improved the performance (Mut-Veh vs Mut-ASO, P=0.0008). The dashed line indicates chance level. **e**, Vision test confirming normal visual ability across all groups (P>0.05 for all comparisons). Data are presented as mean ± s.e.m. *P < 0.05, **P < 0.01, ***P < 0.001. Exact P values, sample sizes, and statistical tests are provided in the Supplementary Table.

**Extended Data Table 1.**
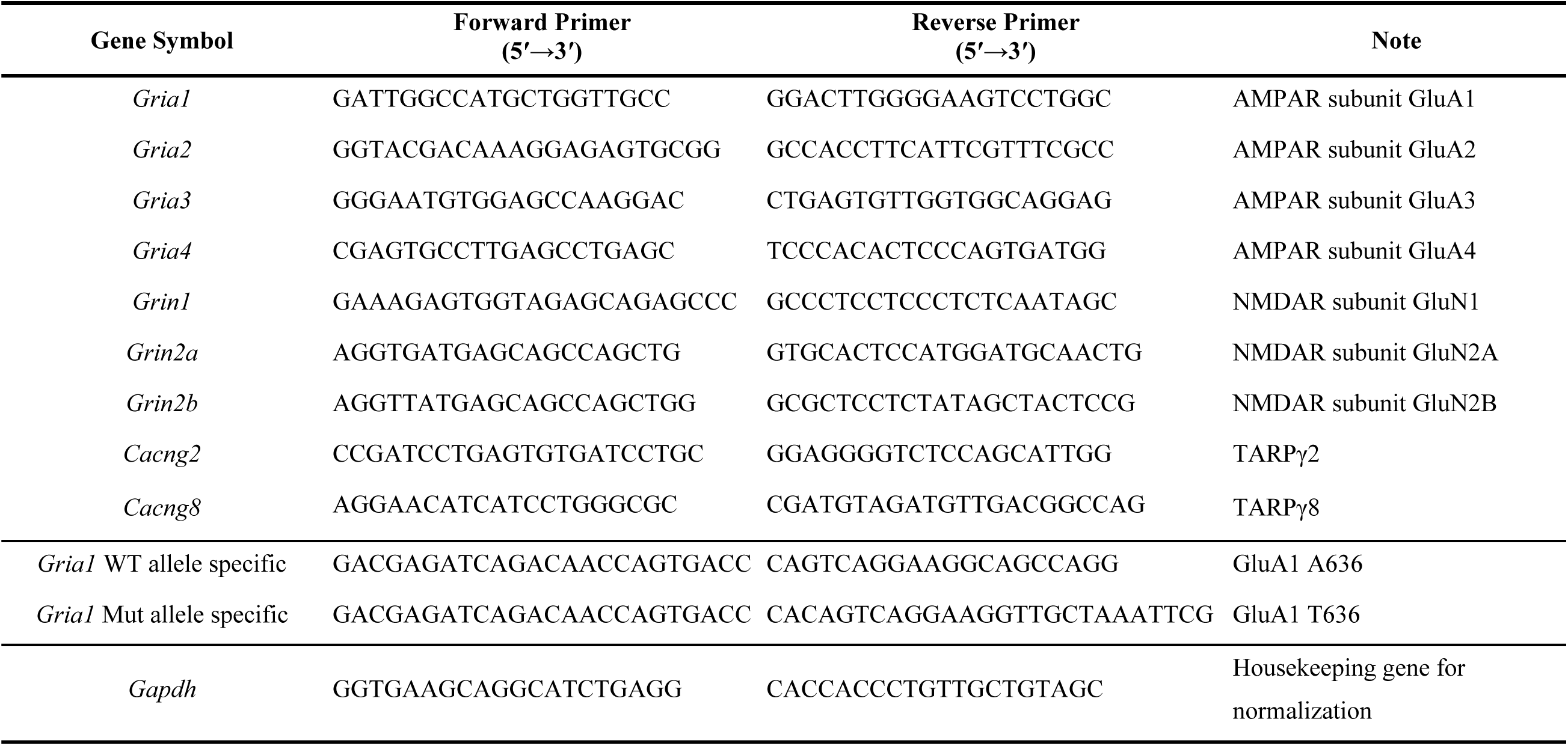
Primer sequences used for RT-qPCR analysis.

## Supplementary Table (Separate Excel file)

Full statistical details for all data plotted in main Figs 1-5, and Extended Data Figs. 1-9.

## References

1 Moretto, E., Murru, L., Martano, G., Sassone, J. & Passafaro, M. Glutamatergic synapses in neurodevelopmental disorders. Progress in neuro-psychopharmacology & biological psychiatry 84, 328–342 (2018). 10.1016/j.pnpbp.2017.09.014

2 Volk, L., Chiu, S. L., Sharma, K. & Huganir, R. L. Glutamate synapses in human cognitive disorders. Annu Rev Neurosci 38, 127–149 (2015). 10.1146/annurev-neuro-071714-033821

3 Nisar, S. et al. Genetics of glutamate and its receptors in autism spectrum disorder. Mol Psychiatry 27, 2380–2392 (2022). 10.1038/s41380-022-01506-w

4 Tvergaard, N. K. et al. Unraveling GRIA1 neurodevelopmental disorders: Lessons learned from the p.(Ala636Thr) variant. Clin Genet 106, 427–436 (2024). 10.1111/cge.14577

5 Schwenk, J. et al. Regional diversity and developmental dynamics of the AMPA-receptor proteome in the mammalian brain. Neuron 84, 41–54 (2014). 10.1016/j.neuron.2014.08.044

6 Geisheker, M. R. et al. Hotspots of missense mutation identify neurodevelopmental disorder genes and functional domains. Nat Neurosci 20, 1043–1051 (2017). 10.1038/nn.4589

7 Ismail, V. et al. Identification and functional evaluation of missense and truncation variants in individuals with ID: An emerging neurodevelopmental syndrome. American Journal of Human Genetics 109, 1217–1241 (2022). 10.1016/j.ajhg.2022.05.009

8 Vogel, M. W., Caston, J., Yuzaki, M. & Mariani, J. The Lurcher mouse: fresh insights from an old mutant. Brain Res 1140, 4–18 (2007). 10.1016/j.brainres.2005.11.086

9 Zuo, J. et al. Neurodegeneration in Lurcher mice caused by mutation in delta2 glutamate receptor gene. Nature 388, 769–773 (1997). 10.1038/42009

10 Taverna, F. et al. The Lurcher mutation of an alpha-amino-3-hydroxy-5-methyl-4-isoxazolepropionic acid receptor subunit enhances potency of glutamate and converts an antagonist to an agonist. J Biol Chem 275, 8475–8479 (2000). 10.1074/jbc.275.12.8475

11 Klein, R. M. & Howe, J. R. Effects of the lurcher mutation on GluR1 desensitization and activation kinetics. J Neurosci 24, 4941–4951 (2004). 10.1523/JNEUROSCI.0660-04.2004

12 Hill, S. F. & Meisler, M. H. Antisense Oligonucleotide Therapy for Neurodevelopmental Disorders. Dev Neurosci 43, 247–252 (2021). 10.1159/000517686

13 Bennett, C. F., Krainer, A. R. & Cleveland, D. W. Antisense Oligonucleotide Therapies for Neurodegenerative Diseases. Annu Rev Neurosci 42, 385–406 (2019). 10.1146/annurev-neuro-070918-050501

14 Crooke, S. T., Baker, B. F., Crooke, R. M. & Liang, X. H. Antisense technology: an overview and prospectus. Nat Rev Drug Discov 20, 427–453 (2021). 10.1038/s41573-021-00162-z

15 Wenthold, R. J., Petralia, R. S., Blahos, J. II, & Niedzielski, A. S. Evidence for multiple AMPA receptor complexes in hippocampal CA1/CA2 neurons. J Neurosci 16, 1982–1989 (1996). 10.1523/JNEUROSCI.16-06-01982.1996

16 Guo, C. & Ma, Y. Y. Calcium Permeable-AMPA Receptors and Excitotoxicity in Neurological Disorders. Front Neural Circuits 15, 711564 (2021). 10.3389/fncir.2021.711564

17 Kumar, S. S., Bacci, A., Kharazia, V. & Huguenard, J. R. A developmental switch of AMPA receptor subunits in neocortical pyramidal neurons. J Neurosci 22, 3005–3015 (2002). 20026285

18 Ho, M. T. W. et al. Developmental expression of Ca-permeable AMPA receptors underlies depolarization-induced long-term depression at mossy fiber-CA3 pyramid Synapses. J Neurosci 27, 11651–11662 (2007). 10.1523/Jneurosci.2671-07.2007

19 Henley, J. M. & Wilkinson, K. A. Synaptic AMPA receptor composition in development, plasticity and disease. Nat Rev Neurosci 17, 337–350 (2016). 10.1038/nrn.2016.37

20 Clem, R. L. & Huganir, R. L. Calcium-permeable AMPA receptor dynamics mediate fear memory erasure. Science 330, 1108–1112 (2010). 10.1126/science.1195298

21 Plant, K. et al. Transient incorporation of native GluR2-lacking AMPA receptors during hippocampal long-term potentiation. Nat Neurosci 9, 602–604 (2006). 10.1038/nn1678

22 Park, P. et al. Calcium-Permeable AMPA Receptors Mediate the Induction of the Protein Kinase A-Dependent Component of Long-Term Potentiation in the Hippocampus. J Neurosci 36, 622–631 (2016). 10.1523/JNEUROSCI.3625-15.2016

23 McCutcheon, J. E. et al. Group I mGluR activation reverses cocaine-induced accumulation of calcium-permeable AMPA receptors in nucleus accumbens synapses via a protein kinase C-dependent mechanism. J Neurosci 31, 14536–14541 (2011). 10.1523/JNEUROSCI.3625-11.2011

24 Liu, S. J. & Zukin, R. S. Ca2+-permeable AMPA receptors in synaptic plasticity and neuronal death. Trends in neurosciences 30, 126–134 (2007). 10.1016/j.tins.2007.01.006

25 Whitehead, G., Regan, P., Whitcomb, D. J. & Cho, K. Ca(2+)-permeable AMPA receptor: A new perspective on amyloid-beta mediated pathophysiology of Alzheimer’s disease. Neuropharmacology 112, 221–227 (2017). 10.1016/j.neuropharm.2016.08.022

26 Bowie, D. & Mayer, M. L. Inward rectification of both AMPA and kainate subtype glutamate receptors generated by polyamine-mediated ion channel block. Neuron 15, 453–462 (1995). 10.1016/0896-6273(95)90049-7

27 Cull-Candy, S. G. & Farrant, M. Ca(2+) - permeable AMPA receptors and their auxiliary subunits in synaptic plasticity and disease. J Physiol 599, 2655–2671 (2021). 10.1113/JP279029

28 Kwon, H. S. & Koh, S. H. Neuroinflammation in neurodegenerative disorders: the roles of microglia and astrocytes. Transl Neurodegener 9, 42 (2020). 10.1186/s40035-020-00221-2

29 Ransohoff, R. M. How neuroinflammation contributes to neurodegeneration. Science 353, 777–783 (2016). 10.1126/science.aag2590

30 Hanada, T. et al. Perampanel: a novel, orally active, noncompetitive AMPA-receptor antagonist that reduces seizure activity in rodent models of epilepsy. Epilepsia 52, 1331–1340 (2011). 10.1111/j.1528-1167.2011.03109.x

31 Maher, M. P. et al. Discovery and Characterization of AMPA Receptor Modulators Selective for TARP-gamma8. J Pharmacol Exp Ther 357, 394–414 (2016). 10.1124/jpet.115.231712

32 Backstrom, E. et al. Tissue pharmacokinetics of antisense oligonucleotides. Mol Ther Nucleic Acids 35, 102133 (2024). 10.1016/j.omtn.2024.102133

33 Salpietro, V. et al. AMPA receptor GluA2 subunit defects are a cause of neurodevelopmental disorders. Nature communications 10, 3094 (2019). 10.1038/s41467-019-10910-w

34 Zamanillo, D. et al. Importance of AMPA receptors for hippocampal synaptic plasticity but not for spatial learning. Science 284, 1805–1811 (1999).

35 Reisel, D. et al. Spatial memory dissociations in mice lacking GluR1. Nat Neurosci 5, 868–873 (2002). 10.1038/nn910

36 Herguedas, B. et al. Mechanisms underlying TARP modulation of the GluA1/2-gamma8 AMPA receptor. Nature communications 13, 734 (2022). 10.1038/s41467-022-28404-7

37 Chen, L., Durr, K. L. & Gouaux, E. X-ray structures of AMPA receptor-cone snail toxin complexes illuminate activation mechanism. Science 345, 1021–1026 (2014). 10.1126/science.1258409

38 Nakagawa, T., Wang, X. T., Miguez-Cabello, F. J. & Bowie, D. The open gate of the AMPA receptor forms a Ca(2+) binding site critical in regulating ion transport. Nat Struct Mol Biol 31, 688–700 (2024). 10.1038/s41594-024-01228-3

39 Miguez-Cabello, F. et al. GluA2-containing AMPA receptors form a continuum of Ca(2+)-permeable channels. Nature 641, 537–544 (2025). 10.1038/s41586-025-08736-2

40 Banke, T. G. & Barria, A. Transient Enhanced GluA2 Expression in Young Hippocampal Neurons of a Fragile X Mouse Model. Frontiers in synaptic neuroscience 12, 588295 (2020). 10.3389/fnsyn.2020.588295

41 Coutelier, M. et al. GRID2 mutations span from congenital to mild adult-onset cerebellar ataxia. Neurology 84, 1751–1759 (2015). 10.1212/WNL.0000000000001524

42 Guzman, Y. F. et al. A gain-of-function mutation in the GRIK2 gene causes neurodevelopmental deficits. Neurol Genet 3, e129 (2017). 10.1212/NXG.0000000000000129

