## Supplementary material for "Rescuing Neurodevelopmental Deficits in AMPA Receptor Gain-of-Function Mutant": Methods

### Animals

All animal procedures were approved by the Animal Care and Use Committees at Academia Sinica and Johns Hopkins University and were performed in accordance with institutional guidelines. *Gria1*-A636T mice (generated by the Johns Hopkins Transgenic Core) and Thy1-GFP (line M) transgenic mice were maintained on a C57BL/6 background. All animals were sex-separated and housed in groups of 2-5 mice per cage under standard conditions (room temperature ~22°C, relative humidity ~50%, 12-hr light/dark cycle) with ad libitum access to food and water.

### Generations of *Gria1*-A636T mice

*Gria1*-A636T mice were generated using CRISPR/Cas9-mediated genome editing. Briefly, one-cell stage fertilized C57BL/6 mouse embryos were injected with Cas9 protein, crRNA, tracrRNA, and homology directed repair (HDR) DNA template. The CRISPR guide RNA target sequence was 5'- CTCCTCATACACAGCCAACC-3'. The HDR template (5'-GATGGGAGACACCATCCTTTCCACAGTCAGGAAGGTTGCTAAATTCGCTGTGTATGA GGAGATGATAATCAAAGTGAAGAA-3', minus strand) introduced the A636T mutation and five silent blocking mutations to prevent Cas9 from re-cutting and to facilitate screening. Both the donor template and crRNAs were synthesized by Integrated DNA Technologies (IDT). Progeny were screened by PCR using primers flanking the introduced mutations, followed by diagnostic digestion with ApoI. The PCR primers included forward (5'-CTTACAGAGTACGGGG TGCAATC-3') and reverse (5'-GTTAGGAAGGAA TGCAGAGCCTC-3'), resulting in a 437 bp PCR product. Mutant products digested with ApoI yielded 267 bp and 170 bp fragments. Sanger sequencing was performed on all mutant mice to confirm accurate integration of the HDR template into the genome. To address potential off-target effects, regions with high sequence homology between *Gria1*, *Gria4*, and *Grin1* were PCR amplified and sequenced. Founders that passed restriction enzyme digestion and sequencing tests were backcrossed to naïve C57BL/6 mice for at least four generations prior to experiments to minimize potential off-target variation.

### Genotyping

Genotyping was performed by PCR-based detection of genomic DNA extracted from tail biopsies. Tail tissue (~2 mm) was incubated in 25 mM NaOH and 0.2 mM EDTA at 100°C for 30 min, then neutralized with an equal volume of 40 mM Tris-EDTA buffer (pH 5.5). The PCR protocol was conducted as follows: 94°C for 2 minutes, 34 cycles at 94°C for 30sec, 64°C for 30sec, 72°C for 30sec. The reaction was completed at 72°C for an additional 3 minutes and held at 4°C. The primers used to identify the A636T sequences are A636T_RJ1275 (5'-CAGAGTAC GGGGTGCAATCATGT-3'), A636T_RJ1276 (5'-CAGTCAGGAAGGCAGCCAGGTTG-3'), A636T_RJ1277-23 (5'-CAGTCA GGAAGGTTGCTAAATTC-3') and A636T_RJ1277-26 (5'-GAGCAGTCAGGAAGGTTGCTAAATTC-3'). Amplification with the A636T_RJ1275/ RJ1276 primer pair yielded a 190 bp product specific to the WT allele, while amplification with A636T_RJ1277-23/RJ1277-26 yielded a 220bp product specific for the A636T Mut allele.

### Behavioral studies

Male or female *Gria1* WT or Mut littermates were group housed after weaning, with both genotypes represented in each cage. All tests were conducted in animal behavioral facilities with the examiners blinded to animal genotype. For mice that underwent multiple tests, the sequence was as follows: open field, grooming, three-chamber social test, Y-maze, reciprocal social interaction, elevated plus maze and Morris water maze. Mice were acclimated to the testing room for at least 30 minutes prior to the start of each behavioral test. All sessions were videotaped, and movement tracking was performed using the ANY-maze behavioral tracking system (Stoelting Co., Wood Dale, IL).

-Three-chamber social test

The three-chamber social test is a widely used test to assess social preference and social novelty in rodents. Test mice were habituated for 10 min in an empty Plexiglas arena (60 × 40 × 25 cm, L × W × H) divided into three interconnected chambers with sliding gates removed. During the sociability phase, each mouse was allowed to explore the apparatus for 10 minutes, with access to two wire cups: one containing an unfamiliar same-sex conspecific (Mouse 1) and the other left empty. The positions of the social and empty cups were counterbalanced across trials. In the social novelty phase, a second unfamiliar mouse (Mouse 2) was placed in the previously empty wire cup, and the test mouse was allowed to explore for another 10 minutes. Time spent within 2 cm of each cup was automatically recorded and used as a measure of social interest and novelty recognition.

Reference mice (Mouse 1 and Mouse 2) were same-sex C57BL/6 mice that were age-matched or 1-2 weeks younger than the test mice. To minimize stress-related artifacts, reference mice were habituated to the wire cup for 20 minutes per day for at least three consecutive days prior to testing. Mice exhibiting overt stress behaviors, such as persistent cup biting, excessive freezing, or excessive urination/defecation, were excluded from use as reference animals.

- Reciprocal social interaction

Test mice and age-matched unfamiliar conspecifics were initially placed in opposite corners of a neutral Plexiglas arena (40×40×38 cm, L×W×H), separated by a diagonal divider. After a 5-min habituation period, the divider was removed to allow free interaction. Social behaviors, including sniffing (nose-to-nose or nose-to-anogenital), following, and push-crawl behaviors, were scored manually by trained observers blinded to genotype.

- Y-maze test

The Y-maze test was used to assess spatial short-term reference memory in mice. Test mice were placed in a transparent Y-shaped maze with three arms (each 35×7×15 cm, L×W×H), each distinguished by unique extra-maze visual cues. During the 5-min familiarization period, one arm (left or right) was blocked, and mice were allowed to freely explore the other two arms. After a 2-min rest period in a holding cage, mice were reintroduced into the maze for a 3-min test phase with all three arms accessible. Time spent in each arm was recorded automatically. A discrimination index was calculated as: % time in novel arm / (% time in novel arm + % time in familiar arm).

- Morris water maze

The classic Morris Water Maze was used to assess long-term spatial learning and memory, following established protocols for mice with minor modifications. Briefly, mice were trained over five days (two sessions/day, four trials/session) to locate a submerged platform using distal visual cues. Each trial lasted a maximum of 60 sec, with a 15-sec resting period on the platform after successful locating. Swim paths and latencies were recorded to generate learning curves. A probe trial on day six, with the platform removed, evaluated memory retention based on time spent in the target quadrant. On day seven, the platform was moved to the opposite quadrant to assess reversal learning, following the same training protocol. A second probe trial on day twelve assessed memory retention of the new platform location.

- Grooming

Test mice were individually placed in a sound-attenuated arena (25×25×25 cm) under infrared illumination during their active phase. Grooming behaviors, including face wiping, head scratching, and full body grooming, were recorded over 15 min and manually scored by trained observers blinded to genotype. This assessment captures self-directed behaviors that are indicative of repetitive behaviors.

- Open field test

Test mice were individually placed in the center of a Plexiglas arena (40×40×32 cm, L×W×H) and allowed to explore freely for 30 min. Locomotor activity, including velocity and total distance traveled, was automatically recorded. Anxiety levels were inferred from the percentage of time spent in the periphery (10 cm zone along the walls) versus the center of the arena.

- Elevated plus maze

Test mice were placed in the center area of an elevated maze, which was raised 38 cm above the ground. The maze consisted of two closed arms and two open arms (each 30×5 cm, L×W), connected by the center area. Each mouse was allowed to explore freely for 5 min, and its position was automatically tracked. Anxiety levels were assessed by measuring the percentage of time spent in the closed arms relative to the total arm exploration.

Western blot analysis

Brain tissues, including hippocampus, and when indicated, prefrontal cortex, somatosensory cortex, thalamus or cerebellum, were dissected from *Gria1* WT and Mut littermates at the juvenile (3-4 weeks) or adult (2-3 months) stages. Tissues were homogenized by 12 passes through a 26-gauge needle in ice-cold RIPA buffer (150mM NaCl, 25mM Tris-HCl (pH 7.4), 1% NP-40, 0.1% sodium deoxycholate, 1mM EDTA, 1mM EGTA) containing protease and phosphatase inhibitors (Complete Mini and PhosSTOP, Roche). Homogenates were centrifuged at 13,000 rpm for 15 min at 4°C to remove debris. Supernatants were collected for protein quantification and Western blot analyses, probing with primary antibodies against GluA1 (homemade, 4.9D), GluA2 (homemade, 32.191), GluA3 (homemade, JH4300), GluA4 (Millipore, AB1508), GluN1 (Synaptic systems, 114011), mGluR5 (Abcam, ab76316), TARPr2 (homemade, JH6520), TARPr8 (homemade, JH6518), and GAPDH (Novus, NB600-502) or Tubulin (Sigma, T6074).

Magnetic resonance imaging (MRI)

Mice were anesthetized with 5% isoflurane for induction and maintained at 1.5-2.0% during scanning, with an airflow of 1 L/min airflow. They were positioned in the prone orientation and secured with a custom-made head holder in the magnet of a 7-T Biospec 70/20 MRI scanner (Bruker, Ettlingen, Germany) at the Animal Image Facility, BioTReC. Image acquisition was performed using an 86-millimeter birdcage transmitter coil and a separate quadratic array coil with a 2x2 coil element topology for signal reception. Multi-slice axial turbo spin echo images were acquired with a field of view of 2.0x2.0 $cm^2$ field of view, 256x256 matrix size, 4100 ms repetition time, 40 ms echo time, 0.5 mm slice thickness, four averages, and 32 slices for chronological views and 25 slices for sagittal views. Anatomical segmentation and volumetric analyses were performed semi-manually using Amira software (Thermo Fisher Scientific), which facilitates 3D reconstruction and measurement of brain structures.

Dendritic morphology and spine density analysis

*Gria1* WT and Mut littermates were perfused with PBS followed by 4% paraformaldehyde in PBS at room temperature. Brains were fixed overnight, coronal sectioned at 250µm thickness, and mounted with PermaFluor mounting medium (Thermo Fisher Scientific). GFP-expressing CA1 neurons in the hippocampus or L2/3 pyramidal neurons in the somatosensory cortex were imaged using a Zeiss LSM 880 confocal microscope with a 40× glycerol objective, acquiring Z-series images at 0.5µm intervals (512x512 resolution) with tile scanning to capture complete dendritic arbors. 3D reconstructions and quantifications of the dendritic length, branch points and complexity were performed using Imaris 9.5 software (Bitplane) with the filament tracer module.

For spine density assessment, high-resolution images (1,024 x 1,024 resolution, 0.5µm z-step, 2X digital zoom) of the first few splits of apical or basal secondary dendrites were imaged. All visible protrusions from dendrites regardless of the shape were counted as spines and normalized to the length of the individual dendrite. Dendrite segments greater than 200µm per neuron were analyzed. Segments exceeding 200µm per neuron were analyzed. Data were collected from four neurons per animal across four WT/Mut littermate pairs.

TUNEL Labeling and immunohistochemistry

Perfused brain samples were cryoprotected in 30% sucrose in phosphate-buffered saline (PBS), embedded in frozen section compound (VWR), and sectioned into 30µm coronal slices using a cryostat (Leica). Sections containing the dorsal hippocampus were mounted onto glass slides and air-dried at room temperature (RT) for an hour. TUNEL labeling was performed using the In Situ Cell Death Detection Kit, TMR Red (Roche, 12156792910) according to the manufacturer's instructions. Briefly, sections were permeabilized with 0.25% Triton X-100, blocked with 3% $H_2O_2$ in methanol (RT, 10 min), and incubated with the TUNEL reaction mixture (1:11 enzyme to label ratio) at 37°C for 1 hour in a humidified chamber. After PBS washes, sections were co-stained with NeuN (Millipore, ABN78) and GFAP (Sigma, G3893/GA5) antibodies, followed by Alex Flour-conjugated secondary antibodies (Invitrogen). Nuclear counterstaining was performed with DAPI (0.1 µg/mL). As Iba1(Abcam, ab5076) is sensitive to menthol treatment, glia staining for IbaI and GFAP was performed on separated sections, omitting the $H_2O_2$/methanol and TUNEL staining. TUNEL-positive or glia cell densities were quantified over the entire CA1, CA2, CA3 and DG regions, normalized to the analyzed area from three sections per brain, with data collected from at least four animals per condition. Only DAPI-costained signals were counted as dying cells or glia to minimize false positives.

Electrophysiology

- Slice preparation

Acute coronal hippocampal slices (300 µm thick) were prepared from *Gria1* WT and Mut littermates at P6-9 and P19-24. One animal was recorded per day in an interleaved, randomized sequence. Mice were anesthetized with the inhalational isoflurane and decapitated. Brains were rapidly extracted and immersed in ice-cold, oxygenated (95% $O_2$, 5% $CO_2$) low-$Ca^{2+}$/high-$Mg^{2+}$ dissection buffer (2.6 mM KCl, 1.25 mM $NaH_2PO_4$, 26 mM $NaHCO_3$, 211 mM sucrose, 11 mM glucose, 0.5 mM $CaCl_2$ and 7 mM $MgCl_2$). Slices were sectioned using a vibratome (Leica VT1200S), and slices containing the dorsal hippocampus were collected and recovered in a static submersion chamber filled with oxygenated artificial cerebrospinal fluid (ACSF, 119 mM NaCl,

2.5 mM KCl, 1 mM $NaH_2PO_4$, 2.5 mM $CaCl_2$, 1.3 mM $MgSO_4$, 26.2 mM $NaHCO_3$ and 11 mM glucose) at 32°C for 32 min before recording.

- Whole-cell recordings and AMPAR mEPSCs

Whole-cell voltage-clamp recordings were performed on CA1 pyramidal neurons at room temperature (~24°C) in ACSF containing 1 μM tetrodotoxin (TTX) and 100 μM picrotoxin, perfused at a flow rate of 2 ml/min. Patch pipettes (4-6 MΩ) were filled with internal solution (115 mM Cs-$MeSO_3$, 0.4 mM EGTA, 5 mM TEA-Cl, 2.8 mM NaCl, 20 mM HEPES, 3 mM Mg-ATP, 0.5 mM $Na_2$-GTP, pH 7.2-7.25, osmolality 290-295 mOsm). Neurons were clamped at -70mV. To ensure stable intracellular conditions, a 3-min equilibration period was allowed after establishing the whole-cell configuration before data analysis for mEPSC analysis.

- Basal AMPAR-mediated currents

Schaffer collateral pathways were stimulated at 0.1 Hz via stratum radiatum stimulation. AMPAR-mediated eEPSCs were recorded at -70mV to establish a stable baseline and determine the baseline holding current. Subsequently, 10uM NBQX was applied to block AMPAR responses to access potential AMPAR-mediated leaky currents.

- AMPA/NMDA ratio

AMPAR-eEPSCs were recorded at -70mV. After complete blockade of AMPAR responses with 10 μM NBQX, NMDAR-eEPSCs were recorded at +50mV. Twelve consecutive responses were averaged for each holding potential. AMPAR current amplitudes were measured at -70mV, while NMDAR currents was quantified 50-60 ms post-stimulation at +50mV.

- Rectification index

AMPAR-eEPSCs were recorded at holding potentials ranging from -70mV to +70mV in 20mV increments. The internal solution contained 100 μM NASPM, a selective CP-AMPAR blocker, and 1 μM QX314 to inhibit voltage-gated sodium channels to ensure stable voltage-clamp conditions. 50μM D-AP5 was included in the bath to block NMDAR-mediated currents. The rectification index was calculated as the ratio of the outward to the inward slope of the I-V curve.

- Calcium-permeable AMPAR

To assess the contribution of CP-AMPARs to synaptic transmission, AMPAR-eEPSCs were recorded at -70 mV. After establishing a 5-min stable baseline, 200μM NASPM was bath-applied, and eEPSCs were monitored for an additional 15 mins. The percentage reduction in EPSC amplitude after NASPM application was used to estimate the synaptic content of CP-AMPARs.

- Data Acquisition & Analysis

Signals were amplified with a MultiClamp 700B amplifier, digitized with a Digidata 1440A (Molecular Devices), and acquired with pClamp 10.5 software at a sampling rate of 10 or 20 kHz. Access resistance (Ra) was monitored throughout, and only recordings with Ra<20 MΩ and <20% change were used for analysis. mEPSCs were detected using a template matching algorithm (Clampfit 10.2), with at least 100 events per cell analyzed.

<u>Primary hippocampal cultures</u>

- Neuronal culture and transfection

Hippocampal neurons were prepared from embryonic day 18 (E18) Sprague-Dawley rat pups and plated onto poly-L-lysine-coated glass coverslips or culture plates in plating medium containing 5% fetal bovine serum, 2% B27, and 2 mM GlutaMAX, and antibiotics (5 U/mL penicillin, and 5 μg/mL streptomycin). The culture medium was replaced with serum-free Neurobasal medium containing the same supplements and antibiotics the next day, and fed weekly until use.

Neurons were transfected at days in 11-15 days in vitro (DIV) with plasmids encoding *Gria1* (Gria1 WT or A636T) ± *Gria2*, along with either GCaMP8s and mCherry for calcium imaging, or Venus for excitotoxicity imaging, using Lipofectamine 2000 (Invitrogen) following manufacture's manual. Imaging experiments were performed 2-3 days after transfection.

- Calcium imaging:

For live calcium imaging, neurons were plated onto 18 mm coverslips in a 12-well culture plate at a density of 140,000 cells per well. Coverslips were transferred to a perfusion chamber and imaged in artificial cerebrospinal fluid (ACSF: 120 mM NaCl, 5 mM KCl, 2 mM $CaCl_2$, 1 mM $MgCl_2$, 10 mM D-glucose, and 10 mM HEPES, pH 7.2-7.4, low osmolarity 260-265 mOsm to match Neurobasal medium) containing 0.5 μM TTX and 50 μM D-APV, and perfused at 2.5 ml/min. After two minutes of baseline imaging, neurons were stimulated with 0.5 μM AMPA for 5 seconds, followed by washout with ACSF. In some experiments, 20 μM NBQX was co-applied with AMPA to confirm AMPAR-dependent responses. All reagents were purchased from Sigma or Abcam.

Images were acquired using an inverted Zeiss LSM880 confocal microscope with a 40x objective at 3Hz (512x512 pixels) from a single focal plane. Regions of interest (ROIs) were defined along secondary dendrites (4-5 segments, total length >200 μm), based on mCherry fluorescence. Background-subtracted GCaMP and mCherry intensities were measured in ImageJ and analyzed in MATLAB. GCaMP signals were normalized to mCherry fluorescence (F = GCaMP/mCherry). The calcium response peak was defined as the maximum $\Delta F/F_0$ within 1.5 minutes after stimulation, where $F_0$ was the mean baseline signal from 1.5 to 0.5 minutes before the rise. Traces were extracted from 1 min before to 2 min after response onset.

-Excitotoxicity imaging:

For longitudinal imaging of AMPA-induced excitotoxicity, hippocampal neurons were plated at a density of 70,000 cells per well in a 24-well culture plate. Two days after transfection, neurons were maintained in their conditioned culture medium and imaged using an ImageXpress Confocal HT.ai High Content Imaging System (Molecular Devices) with a 10x objective (2048x2048 pixels) at 37 °C and 5% $CO_2$ to capture the entire well surface. Following baseline imaging, neurons were treated with 0.5 μM AMPA, returned to the incubator for 16 hours, and then re-imaged under identical conditions. In some experiments, 20 μM NBQX was co-applied with AMPA to determine the contribution of AMPAR activation to excitotoxicity. Cell viability was quantified by comparing pre- and post-treatment images of all from Venus-labeled neurons. Neurons that retained continuous Venus fluorescence and intact morphology were classified as viable, whereas a complete loss of Venus signal or fragmentation of neurites were considered to be indicative excitotoxic damage or cell death.

Pharmacology treatments

To assess the therapeutic potential of AMPAR modulation in *Gria1* Mut mice, we administered perampanel (FYCOMPA; PER; Eisai) and JNJ-55511118 (JNJ; Sigma, SML174) by oral gavage. Treatment began at P7 and continued daily until P21. PER was prepared in water at a concentration of 0.2 or 0.4 mg/ml and administered at 2 mg/kg body weight from P7 to P13, and 4 mg/kg from P14 to P21. JNJ was prepared at 1 mg/ml in 0.25% hydroxypropyl methylcellulose (HMC) in water and administered at 10 mg/kg once daily from P7 to P13, and twice daily from P14 to P21. Control groups received equivalent volumes of the respective vehicles (water for PER; 0.25% HMC for JNJ). Mice were monitored daily for general health and potential adverse effects. Notably, PER-treated mice exhibited signs of ataxia, consistent with known side effects observed in both clinical and preclinical studies. JNJ-treated mice did not display any overt behavioral side effects during the treatment period. At P21, mice were euthanized, and brains were harvested for histological analyses, including TUNEL staining and assessment of gross hippocampal morphology.

<u>ASO design and administration</u>

ASOs were designed as 20-mer gapmers targeting the murine Gria1 transcript carrying the A636T (c.1906G>A) point mutation, along with silent mutations introduced during CRISPR/Cas9-mediated genome editing. The lead ASO (ASO2: 5'-GGTTGCTAAATTCGCTGTGT-3') was synthesized with a phosphorothioate backbone and 2'-O-methoxyethyl (2'-MOE) modifications. A CpG-methylated version ($ASO2^{mC}$) was generated by methylating cytosine residues within all CpG motifs. All ASOs were synthesized by IDT and reconstituted at 5 or 10 µg/µl in sterile PBS with 1% fast green (Abcam, ab146267), and stored at 4°C. Neonatal mice (P2-3) were cryoanesthetized and subjected to intracerebroventricular injection with 5 µg ASO in a final volume of 0.5 or 1.0 µl per pup using a nanoliter injector (Nanoject II, Drummond Scientific). Injections were performed using a stereotaxic frame targeting the lateral ventricles at 1.6 mm anterior to the lambda, 0.8 mm lateral to the midline, and 1.7 mm ventral from the skull surface. Control animals received an equal volume of fast green containing PBS vehicle.

<u>Reverse transcription-quantitative PCR (RT-qPCR) Analysis</u>

Hippocampal tissues were collected from 3-week or 2-mon old *Gria1* WT and Mut mice. Total RNA was extracted using TRIzol-chloroform (Thermo Fisher Scientific; 10296-010), and then reverse transcribed using a RETROscript Reverse Transcription Kit (Thermo Fisher Scientific; AM1710). qPCR was performed using a Maxima SYBR Green/Rox Q-PCR Master Mix (Thermo Fisher Scientific; K0221) on a LightCycler 480 II (Roche). Gene expression levels were normalized to the housekeeping gene *Gapdh*, and relative expression was calculated using the ΔΔCt method. All primer sequences used to assess allele-specific ASO efficacy and expression levels of key glutamate receptor subunits and associated proteins are listed in the Extended Data Table 1.

<u>Statistical analysis</u>

All data were presented as mean ± <u>s</u>tandard <u>e</u>rror of the <u>m</u>ean (SEM). Statistical significance was determined using the Mann-Whitney test, one-way ANOVA, or two-way ANOVA, as indicated in the figure legends. Post-hoc analyses were performed using the Bonferroni test for ANOVA tests. Statistical significance was indicated as: $p<0.05$ (*); $p<0.01$ (**), $p< 0.001$ (***). Full statistical details are provided in the Supplementary Table.
