## Supplementary material for "Rescuing Neurodevelopmental Deficits in AMPA Receptor Gain-of-Function Mutant": Suppl. Table_for statistics

| Figure 1c | Assay Performed | Parameter | Groups | N (animals) | Descriptive Statistics | Statistical Analysis | |
|---|---|---|---|---|---|---|---|
| | | | | | | Mann Whitney test - Two-tailed | |
| | | | | | Mean ± SEM | P value | Significance |
| | Three-chamber social test | Sociability | WT-E | 19 | 78.37 ± 6.36 | p<0.0001 | **** |
| | | | WT-M1 | | 136.60 ± 7.04 | | |
| | | | Mut-E | 19 | 62.93 ± 10.05 | p=0.3284 | ns |
| | | | Mut-M1 | | 76.26 ± 9.48 | | |

| Figure 1d | Assay Performed | Parameter | Groups | N (animals) | Descriptive Statistics | Statistical Analysis | |
|---|---|---|---|---|---|---|---|
| | | | | | | Mann Whitney test - Two-tailed | |
| | | | | | Mean ± SEM | P value | Significance |
| | Three-chamber social test | Social novelty | WT-E | 19 | 54.98 ± 4.30 | p=0.0037 | ** |
| | | | WT-M1 | | 91.00 ± 9.74 | | |
| | | | Mut-E | 19 | 66.91 ± 12.56 | p=0.9770 | ns |
| | | | Mut-M1 | | 61.67 ± 7.742 | | |

| Figure 1e | Assay Performed | Parameter | Groups | N (animals) | Descriptive Statistics | Statistical Analysis | |
|---|---|---|---|---|---|---|---|
| | | | | | | Mann Whitney test - Two-tailed | |
| | | | | | Mean ± SEM | P value | Significance |
| | Reciprocal social interaction | Sniffing | WT | 17 | 54.91 ± 6.38 | p=0.0007 | *** |
| | | | Mut | 18 | 27.01 ± 3.48 | | |
| | | Following | WT | 17 | 11.95 ± 3.40 | p=0.0038 | ** |
| | | | Mut | 18 | 2.58 ± 0.59 | | |
| | | Push-craw | WT | 17 | 6.44 ± 0.93 | p=0.0005 | *** |
| | | | Mut | 18 | 2.59 ± 0.44 | | |

| Figure 1f | Assay Performed | Parameter | Groups | N (animals) | Descriptive Statistics | Statistical Analysis | | | |
|---|---|---|---|---|---|---|---|---|---|
| | | | | | | Mann Whitney test - Two-tailed | | | |
| | | | | | Mean ± SEM | P value | Significance | | |
| | Y maze test | Discrimination index | WT | 16 | 0.7143 ± 0.01923 | p<0.0001 | **** | | |
| | | | Mut | 14 | 0.5161 ± 0.03693 | | | | |
| | | | | | | One sample t test | | | |
| | | | | | | Theoretical mean 0.5 | Significance | P value | Significance |
| | | | WT | | | t=11.15, df=15 | | <0.0001 | **** |
| | | | Mut | | | t=0.4347, df=13 | | 0.6709 | ns |

| Figure 1g | Assay Performed | Parameter | Groups | N (animals) | Descriptive Statistics | Statistical Analysis | | Column statistics | |
|---|---|---|---|---|---|---|---|---|---|
| | | | | | | Two-Way ANOVA | | Bonferroni's multiple comparisons | |
| | | | | | Mean ± SEM | Statistical Test | Significance | P value | Significance |
| | MWM learning test | Section 1 | WT | 16 | 37.11 ± 2.87 | Interaction: F (9, 300) = 1.023, P = 0.4215<br>Genotype: F (1, 300) = 60.43, P value < 0.0001<br>Section: F (9, 300) = 7.270, P value < 0.0001 | | >0.9999 | ns |
| | | | Mut | 16 | 39.18 ± 2.81 | | | | |
| | | Section 2 | WT | 16 | 34.36 ± 2.45 | | | >0.9999 | ns |
| | | | Mut | 16 | 38.41 ± 2.64 | | | | |
| | | Section 3 | WT | 16 | 29.99 ± 2.95 | | | 0.6180 | ns |
| | | | Mut | 16 | 36.89 ± 2.33 | | | | |
| | | Section 4 | WT | 16 | 21.99 ± 2.57 | | | 0.0852 | ns |
| | | | Mut | 16 | 31.75 ± 2.35 | | | | |
| | | Section 5 | WT | 16 | 24.66 ± 2.30 | | | 0.0934 | ns |
| | | | Mut | 16 | 34.29 ± 2.43 | | | | |
| | | Section 6 | WT | 16 | 20.33 ± 1.78 | | | 0.0169 | * |
| | | | Mut | 16 | 32.01 ± 2.46 | | | | |
| | | Section 7 | WT | 16 | 20.37 ± 2.26 | | | 0.1268 | ns |
| | | | Mut | 16 | 29.61 ± 2.39 | | | | |
| | | Section 8 | WT | 16 | 21.02 ± 2.30 | | | 0.0058 | ** |
| | | | Mut | 16 | 33.83 ± 3.97 | | | | |
| | | Section 9 | WT | 16 | 19.07 ± 1.67 | | | 0.0414 | * |
| | | | Mut | 16 | 29.71 ± 3.67 | | | | |
| | | Section 10 | WT | 16 | 18.61 ± 1.75 | | | 0.0022 | ** |
| | | | Mut | 16 | 32.37 ± 2.96 | | | | |

| Figure 1h | Assay Performed | Parameter | Groups | N (animals) | Descriptive Statistics | Statistical Analysis | |
|---|---|---|---|---|---|---|---|
| | | | | | | Mann Whitney test - Two-tailed | |
| | | | | | Mean ± SEM | P value | Significance |
| | MWM probe trial | Quadrant 1 | WT | 16 | 18.78 ± 1.48 | p=0.2349 | ns |
| | | | Mut | 16 | 24.24 ± 2.98 | | |
| | | Quadrant 2 | WT | 16 | 24.00 ± 1.85 | p=0.9778 | ns |
| | | | Mut | 16 | 24.88 ± 2.35 | | |
| | | Quadrant 3 | WT | 16 | 35.87 ± 2.65 | p=0.0266 | * |
| | | | Mut | 16 | 27.74 ± 2.15 | | |
| | | Quadrant 4 | WT | 16 | 21.33 ± 1.62 | p=0.9186 | ns |
| | | | Mut | 16 | 23.16 ± 2.39 | | |

| Figure 1j | Assay Performed | Parameter | Groups | N (animals) | Descriptive Statistics | Statistical Analysis | |
|---|---|---|---|---|---|---|---|
| | | | | | | Mann Whitney test - Two-tailed | |
| | | | | | Mean ± SEM | P value | Significance |
| | Western-GluA1 | Juvenile | WT | 6 | 1.00 ± 0.79 | 0.0022 | ** |
| | | | Mut | 6 | 0.55 ± 0.30 | | |
| | | Adult | WT | 6 | 1.00 ± 0.54 | 0.0022 | ** |
| | | | Mut | 6 | 0.50 ± 0.25 | | |

| Figure 1k | Assay Performed | Parameter | Groups | N (animals) | Descriptive Statistics | Statistical Analysis | |
|---|---|---|---|---|---|---|---|
| | | | | | | Mann Whitney test - Two-tailed | |
| | | | | | Mean ± SEM | P value | Significance |
| | Western-GluA2 | Juvenile | WT | 6 | 1.00 ± 0.04 | 0.0022 | ** |
| | | | Mut | 6 | 0.63 ± 0.02 | | |
| | | Adult | WT | 6 | 1.00 ± 0.05 | 0.0022 | ** |
| | | | Mut | 6 | 0.68 ± 0.03 | | |

| Figure 1l | Assay Performed | Parameter | Groups | N (animals) | Descriptive Statistics | Statistical Analysis | |
|---|---|---|---|---|---|---|---|
| | | | | | | Mann Whitney test - Two-tailed | |
| | | | | | Mean ± SEM | P value | Significance |
| | Western-GluA3 | Juvenile | WT | 6 | 1.00 ± 0.05 | 0.0022 | ** |
| | | | Mut | 6 | 0.73 ± 0.03 | | |
| | | Adult | WT | 6 | 1.00 ± 0.06 | 0.0043 | ** |
| | | | Mut | 6 | 0.68 ± 0.04 | | |

| Figure 1m | Assay Performed | Parameter | Groups | N (animals) | Descriptive Statistics | Statistical Analysis | |
|---|---|---|---|---|---|---|---|
| | | | | | | Mann Whitney test - Two-tailed | |
| | | | | | Mean ± SEM | P value | Significance |
| | Western-GluA4 | Juvenile | WT | 6 | 1.00 ± 0.11 | 0.6991 | ns |
| | | | Mut | 6 | 0.93 ± 0.07 | | |
| | | Adult | WT | 6 | 1.00 ± 0.05 | 0.9372 | ns |
| | | | Mut | 6 | 0.98 ± 0.03 | | |

| Figure 1n | Assay Performed | Parameter | Groups | N (animals) | Descriptive Statistics | Statistical Analysis | |
|---|---|---|---|---|---|---|---|
| | | | | | | Mann Whitney test - Two-tailed | |
| | | | | | Mean ± SEM | P value | Significance |
| | Western-GluN1 | Juvenile | WT | 6 | 1.00 ± 0.06 | 0.2403 | ns |
| | | | Mut | 6 | 0.91 ± 0.05 | | |
| | | Adult | WT | 6 | 1.00 ± 0.09 | 0.3095 | ns |
| | | | Mut | 6 | 0.88 ± 0.10 | | |

| Figure 1o | Assay Performed | Parameter | Groups | N (animals) | Descriptive Statistics | Statistical Analysis | |
|---|---|---|---|---|---|---|---|
| | | | | | | Mann Whitney test - Two-tailed | |
| | | | | | Mean ± SEM | P value | Significance |
| | Western-mGluR5 | Juvenile | WT | 6 | 1.00 ± 0.06 | 0.8182 | ns |
| | | | Mut | 6 | 0.97 ± 0.10 | | |
| | | Adult | WT | 6 | 1.00 ± 0.11 | 0.8182 | ns |
| | | | Mut | 6 | 0.91 ± 0.07 | | |

| Figure 1p | Assay Performed | Parameter | Groups | N (animals) | Descriptive Statistics | Statistical Analysis | |
|---|---|---|---|---|---|---|---|
| | | | | | | Mann Whitney test - Two-tailed | |
| | | | | | Mean ± SEM | P value | Significance |
| | Western-TARP-r8 | Juvenile | WT | 6 | 1.00 ± 0.09 | 0.0022 | ** |
| | | | Mut | 6 | 0.53 ± 0.03 | | |
| | | Adult | WT | 6 | 1.00 ± 0.09 | 0.0022 | ** |
| | | | Mut | 6 | 0.55 ± 0.03 | | |

| Figure 1q | Assay Performed | Parameter | Groups | N (animals) | Descriptive Statistics | Statistical Analysis | |
|---|---|---|---|---|---|---|---|
| | | | | | | Mann Whitney test - Two-tailed | |
| | | | | | Mean ± SEM | P value | Significance |
| | Western-TARP-r2 | Juvenile | WT | 6 | 1.00 ± 0.10 | 0.6991 | ns |
| | | | Mut | 6 | 0.94 ± 0.09 | | |
| | | Adult | WT | 6 | 1.00 ± 0.04 | 0.3939 | ns |
| | | | Mut | 6 | 0.92 ± 0.07 | | |

| Figure 2c | Assay Performed | Parameter | Groups | N (animals) | Descriptive Statistics | Statistical Analysis | | Column statistics | | |
|---|---|---|---|---|---|---|---|---|---|---|
| | | | | | | Two-Way ANOVA | | Bonferroni's multiple comparisons | | |
| | | | | | Mean ± SEM | Statistical Test | Significance | Comparison | P value | Significance |
| | MRI-Hippocampus | P21 | WT | 5 | 15.84 ± 0.63 | Interaction: F (1, 17) = 12.64, P=0.0024<br>Genotype: F (1, 17) = 78.66, P<0.0001<br>Age: F (1, 17) = 0.6673, P=0.4253 | | p21-WT vs. p21-Mut | 0.0078 | ** |
| | | | Mut | 6 | 12.48 ± 0.30 | | | 2m-WT vs. 2m-Mut | <0.0001 | **** |
| | | 2 mon | WT | 5 | 17.57 ± 1.02 | | | p21-WT vs. 2m-WT | 0.4511 | ns |
| | | | Mut | 5 | 9.7 ± 0.44 | | | p21-Mut vs. 2m-Mut | 0.0342 | * |

| Figure 2d | Assay Performed | Parameter | Groups | N (animals) | Descriptive Statistics | Statistical Analysis | | Column statistics | | |
|---|---|---|---|---|---|---|---|---|---|---|
| | | | | | | Two-Way ANOVA | | Bonferroni's multiple comparisons | | |
| | | | | | Mean ± SEM | Statistical Test | Significance | Comparison | P value | Significance |
| | MRI-Cortex | P21 | WT | 5 | 101.4 ± 3.74 | Interaction: F (1, 17) = 0.4959, P=0.4909<br>Genotype: F (1, 17) = 5.133, P=0.0368<br>Age: F (1, 17) = 0.0099, P= 0.9219 | | p21-WT vs. p21-Mut | >0.9999 | ns |
| | | | Mut | 6 | 96.29 ± 3.99 | | | 2m-WT vs. 2m-Mut | 0.9127 | ns |
| | | 2 mon | WT | 5 | 104.1 ± 2.98 | | | p21-WT vs. 2m-WT | >0.9999 | ns |
| | | | Mut | 5 | 94.3 ± 1.07 | | | p21-Mut vs. 2m-Mut | >0.9999 | ns |

| Figure 2e | Assay Performed | Parameter | Groups | N (animals) | Descriptive Statistics | Statistical Analysis | | Column statistics | |
|---|---|---|---|---|---|---|---|---|---|
| | | | | | | Two-Way ANOVA | | Bonferroni's multiple comparisons | |
| | | | | | Mean ± SEM | Statistical Test | Significance | P value | Significance |
| | Sholl Analysis-<br>P14<br>Basal dendrites | Distance- 0 | WT | 5 | 0 ± 0.00 | Distance x Genotype: F (20, 160) = 6.832, P<0.0001<br>Distance: F (20, 160) = 275.8, P<0.0001<br>Genotype: F (1, 8) = 26.02, P=0.0009 | | >0.9999 | ns |
| | | | Mut | 5 | 0.00 ± 0.00 | | | | |
| | | Distance- 10 | WT | 5 | 5 ± 0.21 | | | >0.9999 | ns |
| | | | Mut | 5 | 5.12 ± 0.36 | | | | |
| | | Distance- 20 | WT | 5 | 8.6 ± 0.36 | | | >0.9999 | ns |
| | | | Mut | 5 | 9.64 ± 0.85 | | | | |
| | | Distance- 30 | WT | 5 | 13.44 ± 0.61 | | | >0.9999 | ns |
| | | | Mut | 5 | 13.08 ± 1.04 | | | | |
| | | Distance- 40 | WT | 5 | 16.64 ± 0.50 | | | 0.3475 | ns |
| | | | Mut | 5 | 14.76 ± 1.14 | | | | |
| | | Distance- 50 | WT | 5 | 17.96 ± 0.26 | | | 0.0004 | *** |
| | | | Mut | 5 | 14.56 ± 0.70 | | | | |
| | | Distance- 60 | WT | 5 | 17.08 ± 0.59 | | | <0.0001 | **** |
| | | | Mut | 5 | 13.24 ± 0.90 | | | | |
| | | Distance- 70 | WT | 5 | 15.52 ± 0.76 | | | <0.0001 | **** |
| | | | Mut | 5 | 11.64 ± 1.07 | | | | |
| | | Distance- 80 | WT | 5 | 13.84 ± 0.71 | | | <0.0001 | **** |
| | | | Mut | 5 | 8.76 ± 0.99 | | | | |
| | | Distance- 90 | WT | 5 | 11.52 ± 0.70 | | | <0.0001 | **** |
| | | | Mut | 5 | 6.96 ± 0.89 | | | | |
| | | Distance- 100 | WT | 5 | 8.64 ± 0.72 | | | <0.0001 | **** |
| | | | Mut | 5 | 4.72 ± 0.60 | | | | |
| | | Distance- 110 | WT | 5 | 6.04 ± 0.55 | | | 0.0012 | ** |
| | | | Mut | 5 | 2.84 ± 0.62 | | | | |
| | | Distance- 120 | WT | 5 | 2.96 ± 0.51 | | | >0.9999 | ns |
| | | | Mut | 5 | 1.80 ± 0.43 | | | | |
| | | Distance- 130 | WT | 5 | 1.6 ± 0.32 | | | >0.9999 | ns |
| | | | Mut | 5 | 0.96 ± 0.40 | | | | |
| | | Distance- 140 | WT | 5 | 0.68 ± 0.08 | | | >0.9999 | ns |
| | | | Mut | 5 | 0.60 ± 0.25 | | | | |
| | | Distance- 150 | WT | 5 | 0.4 ± 0.11 | | | >0.9999 | ns |
| | | | Mut | 5 | 0.32 ± 0.16 | | | | |
| | | Distance- 160 | WT | 5 | 0.2 ± 0.06 | | | >0.9999 | ns |
| | | | Mut | 5 | 0.20 ± 0.11 | | | | |
| | | Distance- 170 | WT | 5 | 0.08 ± 0.05 | | | >0.9999 | ns |
| | | | Mut | 5 | 0.12 ± 0.08 | | | | |
| | | Distance- 180 | WT | 5 | 0.08 ± 0.05 | | | >0.9999 | ns |
| | | | Mut | 5 | 0.04 ± 0.04 | | | | |
| | | Distance- 190 | WT | 5 | 0.04 ± 0.04 | | | >0.9999 | ns |
| | | | Mut | 5 | 0.04 ± 0.04 | | | | |
| | | Distance- 200 | WT | 5 | 0 ± 0.00 | | | >0.9999 | ns |
| | | | Mut | 5 | 0 ± 0.00 | | | | |
| | Sholl Analysis-<br>P14<br>Apical dendrites | Distance- 0 | WT | 5 | 0.00 ± 0.00 | Distance x Genotype: F (46, 368) = 6.071, P<0.0001<br>Distance: F (46, 368) = 98.20, P <0.0001<br>Genotype:F (1, 8) = 10.40, P=0.0122 | | >0.9999 | ns |
| | | | Mut | 5 | 0.00 ± 0.00 | | | | |
| | | Distance- 10 | WT | 5 | 0.92 ± 0.05 | | | >0.9999 | ns |
| | | | Mut | 5 | 1.04 ± 0.10 | | | | |
| | | Distance- 20 | WT | 5 | 1.12 ± 0.17 | | | >0.9999 | ns |
| | | | Mut | 5 | 1.40 ± 0.23 | | | | |
| | | Distance- 30 | WT | 5 | 1.40 ± 0.31 | | | >0.9999 | ns |
| | | | Mut | 5 | 2.00 ± 0.33 | | | | |
| | | Distance- 40 | WT | 5 | 1.52 ± 0.34 | | | >0.9999 | ns |
| | | | Mut | 5 | 2.68 ± 0.52 | | | | |
| | | Distance- 50 | WT | 5 | 1.72 ± 0.39 | | | 0.4364 | ns |
| | | | Mut | 5 | 3.16 ± 0.47 | | | | |
| | | Distance- 60 | WT | 5 | 2.96 ± 0.76 | | | >0.9999 | ns |
| | | | Mut | 5 | 3.76 ± 0.61 | | | | |
| | | Distance- 70 | WT | 5 | 4.24 ± 0.74 | | | >0.9999 | ns |
| | | | Mut | 5 | 5.32 ± 0.83 | | | | |
| | | Distance- 80 | WT | 5 | 5.76 ± 0.80 | | | >0.9999 | ns |
| | | | Mut | 5 | 5.56 ± 0.63 | | | | |
| | | Distance- 90 | WT | 5 | 6.96 ± 0.70 | | | >0.9999 | ns |
| | | | Mut | 5 | 6.16 ± 0.51 | | | | |
| | | Distance- 100 | WT | 5 | 8.92 ± 0.80 | | | 0.0003 | *** |
| | | | Mut | 5 | 6.40 ± 0.60 | | | | |
| | | Distance- 110 | WT | 5 | 9.00 ± 0.65 | | | <0.0001 | **** |
| | | | Mut | 5 | 6.28 ± 0.59 | | | | |
| | | Distance- 120 | WT | 5 | 9.32 ± 0.47 | | | <0.0001 | **** |
| | | | Mut | 5 | 6.40 ± 0.37 | | | | |
| | | Distance- 130 | WT | 5 | 9.20 ± 0.51 | | | <0.0001 | **** |
| | | | Mut | 5 | 5.76 ± 0.41 | | | | |
| | | Distance- 140 | WT | 5 | 8.92 ± 0.12 | | | <0.0001 | **** |
| | | | Mut | 5 | 5.20 ± 0.36 | | | | |
| | | Distance- 150 | WT | 5 | 8.36 ± 0.20 | | | <0.0001 | **** |
| | | | Mut | 5 | 4.92 ± 0.52 | | | | |
| | | Distance- 160 | WT | 5 | 7.64 ± 0.50 | | | <0.0001 | **** |
| | | | Mut | 5 | 4.56 ± 0.39 | | | | |
| | | Distance- 170 | WT | 5 | 6.96 ± 0.53 | | | 0.0003 | *** |
| | | | Mut | 5 | 4.44 ± 0.35 | | | | |
| | | Distance- 180 | WT | 5 | 6.04 ± 0.32 | | | 0.0027 | ** |
| | | | Mut | 5 | 3.80 ± 0.44 | | | | |
| | | Distance- 190 | WT | 5 | 4.72 ± 0.51 | | | >0.9999 | ns |
| | | | Mut | 5 | 3.56 ± 0.53 | | | | |
| | | Distance- 200 | WT | 5 | 3.84 ± 0.48 | | | >0.9999 | ns |
| | | | Mut | 5 | 3.16 ± 0.40 | | | | |
| | | Distance- 210 | WT | 5 | 3.52 ± 0.36 | | | >0.9999 | ns |
| | | | Mut | 5 | 2.64 ± 0.48 | | | | |
| | | Distance- 220 | WT | 5 | 3.08 ± 0.42 | | | >0.9999 | ns |
| | | | Mut | 5 | 2.16 ± 0.47 | | | | |
| | | Distance- 230 | WT | 5 | 2.44 ± 0.28 | | | >0.9999 | ns |
| | | | Mut | 5 | 2.28 ± 0.44 | | | | |
| | | Distance- 240 | WT | 5 | 2.20 ± 0.38 | | | >0.9999 | ns |
| | | | Mut | 5 | 2.08 ± 0.51 | | | | |
| | | Distance- 250 | WT | 5 | 2.00 ± 0.33 | | | >0.9999 | ns |
| | | | Mut | 5 | 1.80 ± 0.49 | | | | |
| | | Distance- 260 | WT | 5 | 2.04 ± 0.22 | | | >0.9999 | ns |
| | | | Mut | 5 | 1.60 ± 0.57 | | | | |
| | | Distance- 270 | WT | 5 | 2.00 ± 0.14 | | | >0.9999 | ns |
| | | | Mut | 5 | 1.40 ± 0.49 | | | | |
| | | Distance- 280 | WT | 5 | 1.88 ± 0.22 | | | >0.9999 | ns |
| | | | Mut | 5 | 1.24 ± 0.39 | | | | |
| | | Distance- 290 | WT | 5 | 2.08 ± 0.19 | | | >0.9999 | ns |
| | | | Mut | 5 | 1.08 ± 0.32 | | | | |
| | | Distance- 300 | WT | 5 | 2.16 ± 0.27 | | | >0.9999 | ns |
| | | | Mut | 5 | 1.00 ± 0.32 | | | | |
| | | Distance- 310 | WT | 5 | 1.92 ± 0.27 | | | >0.9999 | ns |
| | | | Mut | 5 | 0.92 ± 0.26 | | | | |
| | | Distance- 320 | WT | 5 | 1.60 ± 0.28 | | | >0.9999 | ns |
| | | | Mut | 5 | 0.80 ± 0.23 | | | | |
| | | Distance- 330 | WT | 5 | 1.44 ± 0.27 | | | >0.9999 | ns |
| | | | Mut | 5 | 0.72 ± 0.21 | | | | |
| | | Distance- 340 | WT | 5 | 1.44 ± 0.28 | | | >0.9999 | ns |

| | | | | | | | | | |
|---|---|---|---|---|---|---|---|---|---|
| | | [illegible] | Mut | 5 | 0.64 ± 0.17 | | | [illegible] | [illegible] |
| | | Distance- 350 | WT | 5 | 1.08 ± 0.31 | | | >0.9999 | ns |
| | | | Mut | 5 | 0.44 ± 0.12 | | | | |
| | | Distance- 360 | WT | 5 | 1.00 ± 0.35 | | | >0.9999 | ns |
| | | | Mut | 5 | 0.44 ± 0.13 | | | | |
| | | Distance- 370 | WT | 5 | 0.80 ± 0.28 | | | >0.9999 | ns |
| | | | Mut | 5 | 0.28 ± 0.12 | | | | |
| | | Distance- 380 | WT | 5 | 0.72 ± 0.36 | | | >0.9999 | ns |
| | | | Mut | 5 | 0.28 ± 0.12 | | | | |
| | | Distance- 390 | WT | 5 | 0.36 ± 0.18 | | | >0.9999 | ns |
| | | | Mut | 5 | 0.28 ± 0.14 | | | | |
| | | Distance- 400 | WT | 5 | 0.32 ± 0.16 | | | >0.9999 | ns |
| | | | Mut | 5 | 0.28 ± 0.14 | | | | |
| | | Distance- 410 | WT | 5 | 0.28 ± 0.14 | | | >0.9999 | ns |
| | | | Mut | 5 | 0.24 ± 0.12 | | | | |
| | | Distance-420 | WT | 5 | 0.20 ± 0.11 | | | >0.9999 | ns |
| | | | Mut | 5 | 0.24 ± 0.12 | | | | |
| | | Distance- 430 | WT | 5 | 0.16 ± 0.12 | | | >0.9999 | ns |
| | | | Mut | 5 | 0.16 ± 0.12 | | | | |
| | | Distance- 440 | WT | 5 | 0.16 ± 0.12 | | | >0.9999 | ns |
| | | | Mut | 5 | 0.16 ± 0.12 | | | | |
| | | Distance- 450 | WT | 5 | 0.16 ± 0.12 | | | >0.9999 | ns |
| | | | Mut | 5 | 0.16 ± 0.12 | | | | |
| | | Distance- 460 | WT | 5 | 0.16 ± 0.12 | | | >0.9999 | ns |
| | | | Mut | 5 | 0.16 ± 0.12 | | | | |

| Figure 2f | Assay Performed | Parameter | Groups | N (animals) | Descriptive Statistics | Statistical Analysis | | Column statistics | |
|---|---|---|---|---|---|---|---|---|---|
| | | | | | | Two-Way ANOVA | | Bonferroni's multiple comparisons | |
| | | | | | Mean ± SEM | Statistical Test | Significance | P value | Significance |
| | Sholl Analysis- P21 Basal dendrites | Distance- 0 | WT | 5 | 0.00 ± 0.00 | | | >0.9999 | ns |
| | | | Mut | 5 | 0.00 ± 0.00 | | | | |
| | | Distance- 10 | WT | 5 | 5.88 ± 0.32 | | | >0.9999 | ns |
| | | | Mut | 5 | 6.68 ± 0.43 | | | | |
| | | Distance- 20 | WT | 5 | 10.2 ± 0.37 | | | >0.9999 | ns |
| | | | Mut | 5 | 11.88 ± 0.68 | | | | |
| | | Distance- 30 | WT | 5 | 15.04 ± 0.50 | | | >0.9999 | ns |
| | | | Mut | 5 | 16.44 ± 0.76 | | | | |
| | | Distance- 40 | WT | 5 | 18.72 ± 0.80 | | | >0.9999 | ns |
| | | | Mut | 5 | 18.64 ± 0.85 | | | | |
| | | Distance- 50 | WT | 5 | 20.36 ± 0.93 | | | >0.9999 | ns |
| | | | Mut | 5 | 18.52 ± 1.01 | | | | |
| | | Distance- 60 | WT | 5 | 20.24 ± 1.05 | | | 0.0018 | ** |
| | | | Mut | 5 | 15.92 ± 1.67 | | | | |
| | | Distance- 70 | WT | 5 | 19.48 ± 1.02 | | | <0.0001 | **** |
| | | | Mut | 5 | 12.00 ± 1.98 | | | | |
| | | Distance- 80 | WT | 5 | 18.36 ± 0.99 | | | <0.0001 | **** |
| | | | Mut | 5 | 8.64 ± 1.42 | | | | |
| | | Distance- 90 | WT | 5 | 16.28 ± 1.11 | | | <0.0001 | **** |
| | | | Mut | 5 | 5.48 ± 1.28 | | | | |
| | | Distance- 100 | WT | 5 | 13.76 ± 0.85 | Distance x Genotype: F (20, 160) = 17.95, P<0.0001<br>Distance: F (20, 160) = 211.5, P <0.0001<br>Genotype: F (1, 8) = 37.48, P=0.0003 | | <0.0001 | **** |
| | | | Mut | 5 | 3.40 ± 0.71 | | | | |
| | | Distance- 110 | WT | 5 | 10.60 ± 0.86 | | | <0.0001 | **** |
| | | | Mut | 5 | 2.16 ± 0.54 | | | | |
| | | Distance- 120 | WT | 5 | 6.96 ± 0.87 | | | <0.0001 | **** |
| | | | Mut | 5 | 1.24 ± 0.42 | | | | |
| | | Distance- 130 | WT | 5 | 4.60 ± 0.86 | | | 0.0072 | ** |
| | | | Mut | 5 | 0.68 ± 0.31 | | | | |
| | | Distance- 140 | WT | 5 | 2.00 ± 0.50 | | | >0.9999 | ns |
| | | | Mut | 5 | 0.40 ± 0.18 | | | | |
| | | Distance- 150 | WT | 5 | 0.80 ± 0.32 | | | >0.9999 | ns |
| | | | Mut | 5 | 0.28 ± 0.12 | | | | |
| | | Distance- 160 | WT | 5 | 0.32 ± 0.14 | | | >0.9999 | ns |
| | | | Mut | 5 | 0.24 ± 0.12 | | | | |
| | | Distance- 170 | WT | 5 | 0.12 ± 0.05 | | | >0.9999 | ns |
| | | | Mut | 5 | 0.20 ±0.09 | | | | |
| | | Distance- 180 | WT | 5 | 0.00 ± 0.00 | | | >0.9999 | ns |
| | | | Mut | 5 | 0.16 ± 0.07 | | | | |
| | | Distance- 190 | WT | 5 | 0.00 ± 0.00 | | | >0.9999 | ns |
| | | | Mut | 5 | 0.12 ± 0.05 | | | | |
| | | Distance- 200 | WT | 5 | 0.00 ± 0.00 | | | >0.9999 | ns |
| | | | Mut | 5 | 0.12 ± 0.05 | | | | |
| | Sholl Analysis- p21 Apical dendrites | Distance- 0 | WT | 5 | 0.00 ± 0.00 | | | >0.9999 | ns |
| | | | Mut | 5 | 0.00 ± 0.00 | | | | |
| | | Distance- 10 | WT | 5 | 0.92 ± 0.05 | | | >0.9999 | ns |
| | | | Mut | 5 | 1.00 ± 0.00 | | | | |
| | | Distance- 20 | WT | 5 | 1.00 ± 0.06 | | | >0.9999 | ns |
| | | | Mut | 5 | 1.48 ± 0.20 | | | | |
| | | Distance- 30 | WT | 5 | 1.40 ± 0.30 | | | 0.3895 | ns |
| | | | Mut | 5 | 3.52 ± 0.62 | | | | |
| | | Distance- 40 | WT | 5 | 2.20 ± 0.33 | | | <0.0001 | **** |
| | | | Mut | 5 | 6.88 ± 0.85 | | | | |
| | | Distance- 50 | WT | 5 | 3.56 ± 0.83 | | | <0.0001 | **** |
| | | | Mut | 5 | 9.56 ± 0.29 | | | | |
| | | Distance- 60 | WT | 5 | 5.48 ± 1.01 | | | <0.0001 | **** |
| | | | Mut | 5 | 10.72 ± 0.46 | | | | |
| | | Distance- 70 | WT | 5 | 6.76 ± 1.06 | | | <0.0001 | **** |
| | | | Mut | 5 | 11.48 ± 0.80 | | | | |
| | | Distance- 80 | WT | 5 | 8.68 ± 0.95 | | | 0.0043 | ** |
| | | | Mut | 5 | 11.84 ± 0.93 | | | | |
| | | Distance- 90 | WT | 5 | 9.48 ± 0.88 | | | >0.9999 | ns |
| | | | Mut | 5 | 10.72 ± 0.95 | | | | |
| | | Distance- 100 | WT | 5 | 9.56 ± 0.87 | | | >0.9999 | ns |
| | | | Mut | 5 | 9.88 ± 1.17 | | | | |
| | | Distance- 110 | WT | 5 | 9.68 ± 0.72 | | | >0.9999 | ns |
| | | | Mut | 5 | 9.44 ± 0.82 | | | | |
| | | Distance- 120 | WT | 5 | 9.20 ± 0.67 | | | >0.9999 | ns |
| | | | Mut | 5 | 7.52 ± 0.95 | | | | |
| | | Distance- 130 | WT | 5 | 8.20 ± 0.54 | | | >0.9999 | ns |
| | | | Mut | 5 | 6.72 ± 0.86 | | | | |
| | | Distance- 140 | WT | 5 | 8.40 ± 0.67 | | | 0.0344 | * |
| | | | Mut | 5 | 5.68 ± 0.91 | | | | |
| | | Distance- 150 | WT | 5 | 7.80 ± 0.43 | | | 0.024 | * |
| | | | Mut | 5 | 5.00 ± 0.75 | | | | |
| | | Distance- 160 | WT | 5 | 7.28 ± 0.69 | | | 0.0689 | ns |
| | | | Mut | 5 | 4.72 ± 0.62 | | | | |
| | | Distance- 170 | WT | 5 | 6.60 ± 0.32 | | | 0.2898 | ns |
| | | | Mut | 5 | 4.40 ± 0.62 | | | | |
| | | Distance- 180 | WT | 5 | 6.08 ± 0.48 | | | 0.5189 | ns |
| | | | Mut | 5 | 4.04 ± 0.55 | | | | |
| | | Distance- 190 | WT | 5 | 5.76 ± 0.48 | | | >0.9999 | ns |
| | | | Mut | 5 | 4.36 ± 0.82 | | | | |
| | | Distance- 200 | WT | 5 | 5.72 ± 0.45 | | | >0.9999 | ns |
| | | | Mut | 5 | 4.04 ± 0.77 | | | | |
| | | Distance- 210 | WT | 5 | 5.28 ± 0.41 | | | >0.9999 | ns |
| | | | Mut | 5 | 3.72 ± 0.52 | | | | |
| | | Distance- 220 | WT | 5 | 5.00 ± 0.47 | | | >0.9999 | ns |
| | | | Mut | 5 | 3.40 ± 0.31 | | | | |
| | | Distance- 230 | WT | 5 | 4.36 ± 0.26 | Distance x Genotype: F (46, 368) = 10.24, P<0.0001<br>Distance: F (46, 368) = 76.13, P <0.0001<br>Genotype: F (1, 8) = 5.492, P=0.0472 | | 0.1563 | ns |
| | | | Mut | 5 | 2.00 ± 0.09 | | | | |
| | | Distance- 240 | WT | 5 | 3.72 ± 0.52 | | | 0.3363 | ns |
| | | | Mut | 5 | 1.56 ± 0.15 | | | | |
| | | Distance- 250 | WT | 5 | 2.80 ± 0.32 | | | >0.9999 | ns |
| | | | Mut | 5 | 1.16 ± 0.16 | | | | |
| | | Distance- 260 | WT | 5 | 2.72 ± 0.30 | | | >0.9999 | ns |
| | | | Mut | 5 | 0.92 ± 0.12 | | | | |
| | | Distance- 270 | WT | 5 | 2.60 ± 0.19 | | | >0.9999 | ns |
| | | | Mut | 5 | 0.80 ± 0.20 | | | | |
| | | Distance- 280 | WT | 5 | 2.52 ± 0.15 | | | 0.5189 | ns |
| | | | Mut | 5 | 0.48 ± 0.21 | | | | |
| | | Distance- 290 | WT | 5 | 2.24 ± 0.19 | | | >0.9999 | ns |
| | | | Mut | 5 | 0.52 ± 0.24 | | | | |
| | | Distance- 300 | WT | 5 | 2.20 ± 0.37 | | | >0.9999 | ns |

| | | | | | | | | | |
|---|---|---|---|---|---|---|---|---|---|
| | | [illegible] | Mut | 5 | 0.40 ±0.21 | | | [illegible] | [illegible] |
| | | Distance- 310 | WT | 5 | 2.56 ± 0.54 | | | 0.2898 | ns |
| | | | Mut | 5 | 0.36 ± 0.13 | | | | |
| | | Distance- 320 | WT | 5 | 2.52 ± 0.45 | | | 0.2492 | ns |
| | | | Mut | 5 | 0.28 ± 0.08 | | | | |
| | | Distance- 330 | WT | 5 | 2.52 ± 0.46 | | | 0.2138 | ns |
| | | | Mut | 5 | 0.24 ± 0.07 | | | | |
| | | Distance- 340 | WT | 5 | 2.68 ± 0.42 | | | 0.1134 | ns |
| | | | Mut | 5 | 0.24 ±0.07 | | | | |
| | | Distance- 350 | WT | 5 | 2.96 ± 0.52 | | | 0.0288 | * |
| | | | Mut | 5 | 0.20 ± 0.09 | | | | |
| | | Distance- 360 | WT | 5 | 2.96 ± 0.62 | | | 0.0288 | * |
| | | | Mut | 5 | 0.20 ± 0.09 | | | | |
| | | Distance- 370 | WT | 5 | 2.80 ± 0.80 | | | 0.0489 | * |
| | | | Mut | 5 | 0.16 ± 0.07 | | | | |
| | | Distance- 380 | WT | 5 | 2.88 ± 0.99 | | | 0.0288 | * |
| | | | Mut | 5 | 0.12 ± 0.08 | | | | |
| | | Distance- 390 | WT | 5 | 2.88 ± 1.20 | | | 0.0288 | * |
| | | | Mut | 5 | 0.12 ± 0.08 | | | | |
| | | Distance- 400 | WT | 5 | 2.52 ± 1.04 | | | 0.1333 | ns |
| | | | Mut | 5 | 0.12 ± 0.08 | | | | |
| | | Distance- 410 | WT | 5 | 2.36 ± 1.00 | | | 0.2898 | ns |
| | | | Mut | 5 | 0.16 ± 0.10 | | | | |
| | | Distance-420 | WT | 5 | 2.04 ± 0.95 | | | 0.7852 | ns |
| | | | Mut | 5 | 0.12 ± 0.08 | | | | |
| | | Distance- 430 | WT | 5 | 1.20 ± 0.58 | | | >0.9999 | ns |
| | | | Mut | 5 | 0.08 ± 0.08 | | | | |
| | | Distance- 440 | WT | 5 | 1.04 ± 0.47 | | | >0.9999 | ns |
| | | | Mut | 5 | 0.08 ± 0.08 | | | | |
| | | Distance- 450 | WT | 5 | 0.64 ± 0.26 | | | >0.9999 | ns |
| | | | Mut | 5 | 0.08 ± 0.08 | | | | |
| | | Distance- 460 | WT | 5 | 0.64 ± 0.26 | | | >0.9999 | ns |
| | | | Mut | 5 | 0.08 ± 0.08 | | | | |

| Figure 2g | Assay Performed | Parameter | Groups | N (animals) | Descriptive Statistics | Statistical Analysis Two-Way ANOVA | | Column statistics Bonferroni's multiple comparisons | |
|---|---|---|---|---|---|---|---|---|---|
| | | | | | Mean ± SEM | Statistical Test | Significance | P value | Significance |
| | | Distance- 0 | WT | 5 | 0.00 ± 0.00 | | | >0.9999 | ns |
| | | | Mut | 5 | 0.00 ± 0.00 | | | | |
| | | Distance- 10 | WT | 5 | 5.38 ± 0.06 | | | >0.9999 | ns |
| | | | Mut | 5 | 6.89 ± 0.80 | | | | |
| | | Distance- 20 | WT | 5 | 9.91 ± 0.39 | | | 0.9784 | ns |
| | | | Mut | 5 | 12.30 ± 0.74 | | | | |
| | | Distance- 30 | WT | 5 | 14.46 ± 0.40 | | | >0.9999 | ns |
| | | | Mut | 5 | 16.22 ± 0.82 | | | | |
| | | Distance- 40 | WT | 5 | 18.02 ± 0.59 | | | >0.9999 | ns |
| | | | Mut | 5 | 18.12 ± 1.51 | | | | |
| | | Distance- 50 | WT | 5 | 19.78 ± 0.73 | | | 0.2705 | ns |
| | | | Mut | 5 | 16.84 ± 2.29 | | | | |
| | | Distance- 60 | WT | 5 | 20.56 ± 0.92 | | | <0.0001 | **** |
| | | | Mut | 5 | 13.92 ± 2.35 | | | | |
| | | Distance- 70 | WT | 5 | 20.03 ± 0.76 | | | <0.0001 | **** |
| | | | Mut | 5 | 10.44 ± 2.05 | | | | |
| | | Distance- 80 | WT | 5 | 18.58 ± 0.74 | | | <0.0001 | **** |
| | | | Mut | 5 | 7.60 ± 1.75 | | | | |
| | | Distance- 90 | WT | 5 | 17.62 ± 0.85 | | | <0.0001 | **** |
| | | | Mut | 5 | 4.57 ± 1.27 | | | | |
| | Sholl Analysis- 2 months Basal dendrites | Distance- 100 | WT | 5 | 15.41 ± 0.89 | Distance x Genotype: F (26, 208) = 24.61, P<0.0001<br>Distance: F (26, 208) = 191.0, P <0.0001<br>Genotype: F (1, 8) = 25.63, P=0.0010 | | <0.0001 | **** |
| | | | Mut | 5 | 2.81 ± 0.94 | | | | |
| | | Distance- 110 | WT | 5 | 12.21 ± 1.11 | | | <0.0001 | **** |
| | | | Mut | 5 | 1.24 ± 0.61 | | | | |
| | | Distance- 120 | WT | 5 | 9.56 ± 1.19 | | | <0.0001 | **** |
| | | | Mut | 5 | 0.47 ± 0.18 | | | | |
| | | Distance- 130 | WT | 5 | 6.21 ± 1.08 | | | <0.0001 | **** |
| | | | Mut | 5 | 0.12 ± 0.10 | | | | |
| | | Distance- 140 | WT | 5 | 3.96 ± 0.97 | | | 0.0225 | * |
| | | | Mut | 5 | 0.12 ± 0.10 | | | | |
| | | Distance- 150 | WT | 5 | 1.68 ± 0.35 | | | >0.9999 | ns |
| | | | Mut | 5 | 0.06 ± 0.05 | | | | |
| | | Distance- 160 | WT | 5 | 0.60 ± 0.10 | | | >0.9999 | ns |
| | | | Mut | 5 | 0.00 ± 0.00 | | | | |
| | | Distance- 170 | WT | 5 | 0.25 ± 0.10 | | | >0.9999 | ns |
| | | | Mut | 5 | 0.00 ± 0.00 | | | | |
| | | Distance- 180 | WT | 5 | 0.05 ± 0.04 | | | >0.9999 | ns |
| | | | Mut | 5 | 0.00 ± 0.00 | | | | |
| | | Distance- 190 | WT | 5 | 0.05 ± 0.04 | | | >0.9999 | ns |
| | | | Mut | 5 | 0.00 ± 0.00 | | | | |
| | | Distance- 200 | WT | 5 | 0.05 ± 0.04 | | | >0.9999 | ns |
| | | | Mut | 5 | 0.00 ± 0.00 | | | | |
| | | Distance- 0 | WT | 5 | 0.00 ± 0.00 | | | >0.9999 | ns |
| | | | Mut | 5 | 0.00 ± 0.00 | | | | |
| | | Distance- 10 | WT | 5 | 1.06 ± 0.05 | | | >0.9999 | ns |
| | | | Mut | 5 | 1.52 ± 0.21 | | | | |
| | | Distance- 20 | WT | 5 | 1.23 ± 0.14 | | | 0.0009 | *** |
| | | | Mut | 5 | 4.54 ± 1.15 | | | | |
| | | Distance- 30 | WT | 5 | 1.84 ± 0.28 | | | <0.0001 | **** |
| | | | Mut | 5 | 9.50 ± 1.65 | | | | |
| | | Distance- 40 | WT | 5 | 3.35 ± 0.34 | | | <0.0001 | **** |
| | | | Mut | 5 | 12.45 ± 2.09 | | | | |
| | | Distance- 50 | WT | 5 | 5.90 ± 0.43 | | | <0.0001 | **** |
| | | | Mut | 5 | 15.41 ± 1.50 | | | | |
| | | Distance- 60 | WT | 5 | 7.77 ± 0.39 | | | <0.0001 | **** |
| | | | Mut | 5 | 14.72 ± 0.84 | | | | |
| | | Distance- 70 | WT | 5 | 9.30 ± 0.55 | | | <0.0001 | **** |
| | | | Mut | 5 | 13.33 ± 0.45 | | | | |
| | | Distance- 80 | WT | 5 | 10.14 ± 0.51 | | | >0.9999 | ns |
| | | | Mut | 5 | 11.37 ± 0.37 | | | | |
| | | Distance- 90 | WT | 5 | 10.95 ± 0.70 | | | 0.0945 | ns |
| | | | Mut | 5 | 8.57 ± 0.90 | | | | |
| | | Distance- 100 | WT | 5 | 10.70 ± 0.89 | | | <0.0001 | **** |
| | | | Mut | 5 | 6.70 ± 0.86 | | | | |
| | | Distance- 110 | WT | 5 | 10.08 ± 0.72 | | | <0.0001 | **** |
| | | | Mut | 5 | 5.04 ± 0.45 | | | | |
| | | Distance- 120 | WT | 5 | 9.79 ± 0.41 | | | <0.0001 | **** |
| | | | Mut | 5 | 3.91 ± 0.64 | | | | |
| | | Distance- 130 | WT | 5 | 9.59 ± 0.28 | | | <0.0001 | **** |
| | | | Mut | 5 | 4.15 ± 0.79 | | | | |
| | | Distance- 140 | WT | 5 | 9.01 ± 0.34 | | | <0.0001 | **** |
| | | | Mut | 5 | 4.22 ± 0.69 | | | | |
| | | Distance- 150 | WT | 5 | 8.74 ± 0.27 | | | <0.0001 | **** |
| | | | Mut | 5 | 4.48 ± 0.89 | | | | |
| | | Distance- 160 | WT | 5 | 8.58 ± 0.36 | | | <0.0001 | **** |
| | | | Mut | 5 | 4.11 ± 0.95 | | | | |
| | | Distance- 170 | WT | 5 | 8.50 ± 0.34 | | | <0.0001 | **** |
| | | | Mut | 5 | 3.37 ± 0.85 | | | | |
| | | Distance- 180 | WT | 5 | 7.98 ± 0.49 | | | <0.0001 | **** |
| | | | Mut | 5 | 2.66 ± 0.76 | | | | |
| | | Distance- 190 | WT | 5 | 7.12 ± 0.43 | | | <0.0001 | **** |
| | | | Mut | 5 | 2.53 ± 0.76 | | | | |
| | | Distance- 200 | WT | 5 | 6.70 ± 0.30 | | | <0.0001 | **** |
| | | | Mut | 5 | 2.42 ± 0.73 | | | | |
| | | Distance- 210 | WT | 5 | 6.12 ± 0.28 | | | <0.0001 | **** |
| | | | Mut | 5 | 2.07 ± 0.63 | | | | |
| | | Distance- 220 | WT | 5 | 5.62 ± 0.30 | | | <0.0001 | **** |
| | Sholl Analysis- 2 months Apical dendrites | | Mut | 5 | 1.92 ± 0.65 | Distance x Genotype: F (50, 400) = 24.58, P<0.0001<br>Distance: F (50, 400) = 93.02, P <0.0001<br>Genotype: F (1, 8) = 27.00, P=0.0008 | | | |
| | | Distance- 230 | WT | 5 | 5.27 ± 0.30 | | | <0.0001 | **** |
| | | | Mut | 5 | 1.22 ± 0.47 | | | | |
| | | Distance- 240 | WT | 5 | 4.61 ± 0.55 | | | 0.0001 | *** |
| | | | Mut | 5 | 0.98 ± 0.34 | | | | |
| | | Distance- 250 | WT | 5 | 4.11 ± 0.75 | | | 0.0009 | *** |
| | | | Mut | 5 | 0.80 ± 0.25 | | | | |
| | | Distance- 260 | WT | 5 | 3.22 ± 0.48 | | | 0.0348 | * |

| | | | | | | |
|---|---|---|---|---|---|---|
| | [illegible] | Mut | 5 | 0.62 ± 0.18 | [illegible] | |
| | Distance- 270 | WT | 5 | 2.73 ± 0.28 | 0.1225 | ns |
| | | Mut | 5 | 0.41 ± 0.17 | | |
| | Distance- 280 | WT | 5 | 2.47 ± 0.41 | 0.1335 | ns |
| | | Mut | 5 | 0.17 ± 0.09 | | |
| | Distance- 290 | WT | 5 | 2.16 ± 0.43 | 0.3085 | ns |
| | | Mut | 5 | 0.06 ± 0.05 | | |
| | Distance- 300 | WT | 5 | 2.42 ± 0.48 | 0.0806 | ns |
| | | Mut | 5 | 0.00 ± 0.00 | | |
| | Distance- 310 | WT | 5 | 2.71 ± 0.41 | 0.0207 | * |
| | | Mut | 5 | 0.00 ± 0.00 | | |
| | Distance- 320 | WT | 5 | 2.55 ± 0.41 | 0.0435 | * |
| | | Mut | 5 | 0.00 ± 0.00 | | |
| | Distance- 330 | WT | 5 | 2.38 ± 0.49 | 0.0928 | ns |
| | | Mut | 5 | 0.00 ± 0.00 | | |
| | Distance- 340 | WT | 5 | 2.65 ± 0.42 | 0.0272 | * |
| | | Mut | 5 | 0.00 ± 0.00 | | |
| | Distance- 350 | WT | 5 | 2.54 ± 0.40 | 0.0469 | * |
| | | Mut | 5 | 0.00 ± 0.00 | | |
| | Distance- 360 | WT | 5 | 2.54 ± 0.42 | 0.0469 | * |
| | | Mut | 5 | 0.00 ± 0.00 | | |
| | Distance- 370 | WT | 5 | 2.32 ± 0.50 | 0.1225 | ns |
| | | Mut | 5 | 0.00 ± 0.00 | | |
| | Distance- 380 | WT | 5 | 2.38 ± 0.59 | 0.0928 | ns |
| | | Mut | 5 | 0.00 ± 0.00 | | |
| | Distance- 390 | WT | 5 | 2.03 ± 0.55 | 0.3961 | ns |
| | | Mut | 5 | 0.00 ± 0.00 | | |
| | Distance- 400 | WT | 5 | 1.43 ± 0.42 | >0.9999 | ns |
| | | Mut | 5 | 0.00 ± 0.00 | | |
| | Distance- 410 | WT | 5 | 0.88 ± 0.29 | >0.9999 | ns |
| | | Mut | 5 | 0.00 ± 0.00 | | |
| | Distance-420 | WT | 5 | 0.64 ± 0.29 | >0.9999 | ns |
| | | Mut | 5 | 0.00 ± 0.00 | | |
| | Distance- 430 | WT | 5 | 0.46 ± 0.28 | >0.9999 | ns |
| | | Mut | 5 | 0.00 ± 0.00 | | |
| | Distance- 440 | WT | 5 | 0.36 ± 0.29 | >0.9999 | ns |
| | | Mut | 5 | 0.00 ± 0.00 | | |
| | Distance- 450 | WT | 5 | 0.36 ± 0.29 | >0.9999 | ns |
| | | Mut | 5 | 0.00 ± 0.00 | | |
| | Distance- 460 | WT | 5 | 0.30 ± 0.24 | >0.9999 | ns |
| | | Mut | 5 | 0.00 ± 0.00 | | |

| **Figure 2h** | **Assay Performed** | **Parameter** | **Groups** | **N (animals)** | **Descriptive Statistics** | **Statistical Analysis** | |
|---|---|---|---|---|---|---|---|
| | | | | | | **Mann Whitney test- Two-tailed** | |
| | | | | | **Mean ± SEM** | **P value** | **Significance** |
| | **Spine density- p21** | Total | WT | 5 | 1.50 ± 0.07 | 0.0079 | ** |
| | | | Mut | 5 | 1.07 ± 0.04 | | |
| | | Basal | WT | 5 | 1.45 ± 0.09 | 0.0079 | ** |
| | | | Mut | 5 | 1.04 ± 0.06 | | |
| | | Apical | WT | 5 | 1.56 ± 0.05 | 0.0079 | ** |
| | | | Mut | 5 | 1.09 ± 0.05 | | |

| **Figure 2i** | **Assay Performed** | **Parameter** | **Groups** | **N (animals)** | **Descriptive Statistics** | **Statistical Analysis** | |
|---|---|---|---|---|---|---|---|
| | | | | | | **Mann Whitney test- Two-tailed** | |
| | | | | | **Mean ± SEM** | **P value** | **Significance** |
| | **Spine density- 2 months** | Total | WT | 5 | 1.60 ± 0.03 | 0.0079 | ** |
| | | | Mut | 5 | 1.12 ± 0.37 | | |
| | | Basal | WT | 5 | 1.54 ± 0.05 | 0.0079 | ** |
| | | | Mut | 5 | 1.17 ± 0.04 | | |
| | | Apical | WT | 5 | 1.66 ± 0.04 | 0.0079 | ** |
| | | | Mut | 5 | 1.08 ± 0.05 | | |

| Figure 3b | Assay Performed | Parameter | Groups | N (slices) | Descriptive Statistics | Statistical Analysis | |
|---|---|---|---|---|---|---|---|
| | | | | | | Mann Whitney test- Two-tailed | |
| | | | | | Mean ± SEM | P value | Significance |
| | AMPAR-mEPSC | Holding current | WT | 21 | 54.81 ± 4.80 | p=0.9003 | ns |
| | | | Mut | 18 | 53.61 ± 4.25 | | |

| Figure 3c | Assay Performed | Parameter | Groups | N (slices) | Descriptive Statistics | Statistical Analysis | |
|---|---|---|---|---|---|---|---|
| | | | | | | Mann Whitney test- Two-tailed | |
| | | | | | Mean ± SEM | P value | Significance |
| | AMPAR-mEPSC | Membrane Resistance | WT | 21 | 298.91 ± 19.45 | p=0.4769 | ns |
| | | | Mut | 18 | 321.57 ± 22.64 | | |

| Figure 3d | Assay Performed | Parameter | Groups | N (slices) | Descriptive Statistics | Statistical Analysis | |
|---|---|---|---|---|---|---|---|
| | | | | | | Mann Whitney test- Two-tailed | |
| | | | | | Mean ± SEM | P value | Significance |
| | AMPAR-mEPSC | Capacitance | WT | 21 | 114.17 ± 6.08 | p=0.0462 | * |
| | | | Mut | 18 | 102.74 ± 4.48 | | |

| Figure 3e | Assay Performed | Parameter | Groups | N (slices) | Descriptive Statistics | Statistical Analysis | | Column statistics | |
|---|---|---|---|---|---|---|---|---|---|
| | | | | | | Two-Way ANOVA | | Bonferroni's multiple comparisons | |
| | | | | | Mean ± SEM | Statistical Test | Significance | P value | Significance |
| | NBQX treatment | Holding current | WT-Before | 8 | 54.81 ± 4.80 | Interaction: F (1, 26) = 8.641e-005, P=0.9927<br>Treatment: F (1, 26) = 0.02611, P=0.8729<br>Genotype: F (1, 26) = 1.962, P=0.1731 | | >0.9999 | ns |
| | | | WT-After | 8 | 53.61 ± 4.25 | | | | |
| | | | WT-Before | 7 | 54.81 ± 4.80 | | | >0.9999 | ns |
| | | | WT-After | 7 | 53.61 ± 4.25 | | | | |

| Figure 3g | Assay Performed | Parameter | Groups | N (slices) | Descriptive Statistics | Statistical Analysis | |
|---|---|---|---|---|---|---|---|
| | | | | | | Mann Whitney test- Two-tailed | |
| | | | | | Mean ± SEM | P value | Significance |
| | AMPAR-mEPSC | Charge | WT | 21 | 90.65 ± 3.27 | p=0.0004 | **** |
| | | | Mut | 18 | 124.03 ± 4.66 | | |

| Figure 3h | Assay Performed | Parameter | Groups | N (slices) | Descriptive Statistics | Statistical Analysis | |
|---|---|---|---|---|---|---|---|
| | | | | | | Mann Whitney test- Two-tailed | |
| | | | | | Mean ± SEM | P value | Significance |
| | AMPAR-mEPSC | Amplitude | WT | 21 | 12.16 ± 0.31 | p=0.0052 | ** |
| | | | Mut | 18 | 13.67 ± 0.55 | | |

| Figure 3i | Assay Performed | Parameter | Groups | N (slices) | Descriptive Statistics | Statistical Analysis | |
|---|---|---|---|---|---|---|---|
| | | | | | | Mann Whitney test- Two-tailed | |
| | | | | | Mean ± SEM | P value | Significance |
| | AMPAR-mEPSC | Decay tau | WT | 21 | 8.64 ± 0.20 | p=0.0038 | ** |
| | | | Mut | 18 | 9.84 ± 0.31 | | |

| Figure 3j | Assay Performed | Parameter | Groups | N (slices) | Descriptive Statistics | Statistical Analysis | |
|---|---|---|---|---|---|---|---|
| | | | | | | Mann Whitney test- Two-tailed | |
| | | | | | Mean ± SEM | P value | Significance |
| | AMPAR-mEPSC | Frequency | WT | 21 | 0.18 ± 0.01 | p<0.0001 | **** |
| | | | Mut | 18 | 0.38 ± 0.04 | | |

| Figure 3k | Assay Performed | Parameter | Groups | N (slices) | Descriptive Statistics | Statistical Analysis | |
|---|---|---|---|---|---|---|---|
| | | | | | | Mann Whitney test- Two-tailed | |
| | | | | | Mean ± SEM | P value | Significance |
| | AMPAR/NMDAR ratio | Ratio | WT | 21 | 1.22 ± 0.12 | p=0.0145 | * |
| | | | Mut | 18 | 1.76 ± 0.16 | | |

| Figure 3l | Assay Performed | Parameter | Groups | N (animals) | Descriptive Statistics | Statistical Analysis | | Column statistics | |
|---|---|---|---|---|---|---|---|---|---|
| | | | | | | Two-Way ANOVA | | Bonferroni's multiple comparisons | |
| | | | | | Mean ± SEM | Statistical Test | Significance | P value | Significance |
| | IV plot-<br>Juvenile | Holding -70 | WT | 17 | -1.00 ± 0.00 | Voltage x Genotype: F (7, 210) = 8.903, P<0.0001<br>Voltage: F (7, 210) = 1639, P<0.0001<br>Genotype: F (1, 30) = 2.252, P=0.1439 | | >0.9999 | ns |
| | | | Mut | 15 | -1.00 ± 0.00 | | | | |
| | | Holding -50 | WT | 17 | -0.82 ± 0.02 | | | >0.9999 | ns |
| | | | Mut | 15 | -0.78 ± 0.02 | | | | |
| | | Holding -30 | WT | 17 | -0.58 ± 0.02 | | | >0.10000 | ns |
| | | | Mut | 15 | -0.53 ± 0.02 | | | | |
| | | Holding -10 | WT | 17 | -0.23 ± 0.02 | | | >0.10001 | ns |
| | | | Mut | 15 | -0.24 ± 0.02 | | | | |
| | | Holding +10 | WT | 17 | 0.08 ± 0.02 | | | >0.10002 | ns |
| | | | Mut | 15 | 0.08 ± 0.02 | | | | |
| | | Holding +30 | WT | 17 | 0.35 ± 0.03 | | | >0.10003 | ns |
| | | | Mut | 15 | 0.30 ± 0.03 | | | | |
| | | Holding +50 | WT | 17 | 0.63 ± 0.04 | | | 0.0123 | * |
| | | | Mut | 15 | 0.50 ± 0.04 | | | | |
| | | Holding +70 | WT | 17 | 0.90 ± 0.04 | | | <0.0001 | **** |
| | | | Mut | 15 | 0.66 ± 0.05 | | | | |

| Figure 3m | Assay Performed | Parameter | Groups | N (animals) | Descriptive Statistics | Statistical Analysis | | Column statistics | |
|---|---|---|---|---|---|---|---|---|---|
| | | | | | | Two-Way ANOVA | | Bonferroni's multiple comparisons | |
| | | | | | Mean ± SEM | Statistical Test | Significance | P value | Significance |
| | IV plot-<br>Neonate | Holding -70 | WT | 12 | -1.00 ± 0.00 | Voltage x Genotype: F (7, 161) = 2.270, P=0.0313<br>Voltage: F (2.364, 54.37) = 771.5, P<0.0001<br>Genotype: F (1, 23) = 8.025e-031, P>0.9999 | | na | na |
| | | | Mut | 13 | -1.00 ± 0.00 | | | | |
| | | Holding -50 | WT | 12 | -0.72 ± 0.02 | | | 0.3976 | ns |
| | | | Mut | 13 | -0.78 ± 0.02 | | | | |
| | | Holding -30 | WT | 12 | -0.48 ± 0.03 | | | >0.9999 | ns |
| | | | Mut | 13 | -0.53 ± 0.03 | | | | |
| | | Holding -10 | WT | 12 | -0.23 ± 0.02 | | | 0.7682 | ns |
| | | | Mut | 13 | -0.28 ± 0.02 | | | | |
| | | Holding +10 | WT | 12 | -0.03 ± 0.03 | | | >0.9999 | ns |
| | | | Mut | 13 | -0.04 ± 0.03 | | | | |
| | | Holding +30 | WT | 12 | 0.17 ± 0.03 | | | >0.9999 | ns |
| | | | Mut | 13 | 0.20 ± 0.02 | | | | |
| | | Holding +50 | WT | 12 | 0.32 ± 0.03 | | | 0.7049 | ns |

| | | | | | | | |
|---|---|---|---|---|---|---|---|
| | [illegible] | Mut | 13 | 0.40 ± 0.03 | | [illegible] | [illegible] |
| | Holding +70 | WT | 12 | 0.47 ± 0.05 | | >0.9999 | ns |
| | | Mut | 13 | 0.54 ± 0.05 | | | |

| Figure 3n | Assay Performed | Parameter | Groups | N (animals) | Descriptive Statistics | Statistical Analysis | | Column statistics | | |
|---|---|---|---|---|---|---|---|---|---|---|
| | | | | | | Two-Way ANOVA | | Bonferroni's multiple comparisons | | |
| | | | | | Mean ± SEM | Statistical Test | Significance | Comparison | P value | Significance |
| | **Rectification index** | Neonate | WT | 12 | 0.62 ± 0.07 | Interaction: F (1, 52) = 8.728, P=0.0032<br>Genotype: F (1, 52) = 1.265, P=0.2658<br>Age: F (1, 52) = 13.86, P=0.0005 | | Neonate WT vs Juvenile WT | 0.0001 | *** |
| | | | Mut | 13 | 0.73 ± 0.07 | | | Neonate WT vs Neonate Mut | >0.9999 | ns |
| | | Juvenile | WT | 17 | 1.07 ± 0.06 | | | Juvenile WT vs Juvenile Mut | 0.0214 | * |
| | | | Mut | 15 | 0.78 ± 0.06 | | | Juvenile Mut vs Neonate Mut | >0.9999 | ns |

| Figure 3o | Assay Performed | Parameter | Groups | N (animals) | Descriptive Statistics | Statistical Analysis | | Column statistics | |
|---|---|---|---|---|---|---|---|---|---|
| | | | | | | Two-Way ANOVA | | Bonferroni's multiple comparisons | |
| | | | | | Mean ± SEM | Statistical Test | Significance | P value | Significance |
| | **NASPM-<br>time course** | Baseline -4 min | WT | 7 | 1.00 ± 0.05 | Time x Genotype: F (19, 209) = 12.16, P <0.0001<br>Time: F (4.819, 53.01) = 20.29, P<0.0001<br>Genotype: 3.264F (1, 11) = 31.28, P=0.0002 | | >0.9999 | ns |
| | | | Mut | 6 | 1.00 ± 0.06 | | | | |
| | | Baseline -3 min | WT | 7 | 0.97 ± 0.04 | | | >0.9999 | ns |
| | | | Mut | 6 | 0.97 ± 0.03 | | | | |
| | | Baseline -2 min | WT | 7 | 1.01 ± 0.05 | | | >0.9999 | ns |
| | | | Mut | 6 | 1.02 ± 0.01 | | | | |
| | | Baseline -1min | WT | 7 | 0.96 ± 0.04 | | | >0.9999 | ns |
| | | | Mut | 6 | 1.01 ± 0.01 | | | | |
| | | Baseline 0min | WT | 7 | 1.03 ± 0.04 | | | >0.9999 | ns |
| | | | Mut | 6 | 0.97 ± 0.01 | | | | |
| | | NASPM 1min | WT | 7 | 0.95 ± 0.07 | | | >0.9999 | ns |
| | | | Mut | 6 | 1.00 ± 0.04 | | | | |
| | | NASPM 2min | WT | 7 | 0.93 ± 0.06 | | | >0.9999 | ns |
| | | | Mut | 6 | 0.95 ± 0.05 | | | | |
| | | NASPM 3min | WT | 7 | 0.92 ± 0.06 | | | >0.9999 | ns |
| | | | Mut | 6 | 0.89 ± 0.04 | | | | |
| | | NASPM 4min | WT | 7 | 0.94 ± 0.06 | | | 0.8642 | ns |
| | | | Mut | 6 | 0.78 ± 0.04 | | | | |
| | | NASPM 5min | WT | 7 | 0.96 ± 0.08 | | | 0.1751 | ns |
| | | | Mut | 6 | 0.65 ± 0.05 | | | | |
| | | NASPM 6min | WT | 7 | 1.02 ± 0.06 | | | 0.011 | * |
| | | | Mut | 6 | 0.64 ± 0.05 | | | | |
| | | NASPM 7min | WT | 7 | 0.94 ± 0.06 | | | 0.0564 | ns |
| | | | Mut | 6 | 0.63 ± 0.05 | | | | |
| | | NASPM 8min | WT | 7 | 0.82 ± 0.06 | | | 0.3178 | ns |
| | | | Mut | 6 | 0.61 ± 0.04 | | | | |
| | | NASPM 9min | WT | 7 | 0.91 ± 0.07 | | | 0.0224 | * |
| | | | Mut | 6 | 0.53 ± 0.03 | | | | |
| | | NASPM 10min | WT | 7 | 0.90 ± 0.07 | | | 0.0172 | * |
| | | | Mut | 6 | 0.50 ± 0.04 | | | | |
| | | NASPM 11min | WT | 7 | 0.86 ± 0.03 | | | <0.0001 | *** |
| | | | Mut | 6 | 0.47 ± 0.03 | | | | |
| | | NASPM 12min | WT | 7 | 0.91 ± 0.04 | | | 0.0001 | *** |
| | | | Mut | 6 | 0.43 ± 0.04 | | | | |
| | | NASPM 13min | WT | 7 | 0.91 ± 0.04 | | | 0.0002 | *** |
| | | | Mut | 6 | 0.39 ± 0.05 | | | | |
| | | NASPM 14min | WT | 7 | 0.90 ± 0.06 | | | 0.0013 | ** |
| | | | Mut | 6 | 0.42 ± 0.04 | | | | |
| | | NASPM 15min | WT | 7 | 0.92 ± 0.04 | | | 0.0012 | ** |
| | | | Mut | 6 | 0.40 ± 0.06 | | | | |

| Figure 3p | Assay Performed | Parameter | Groups | N (slices) | Descriptive Statistics | Statistical Analysis | |
|---|---|---|---|---|---|---|---|
| | | | | | | Mann Whitney test- Two-tailed | |
| | | | | | Mean ± SEM | P value | Significance |
| | **NASPM sensitivity** | Baseline normalized AMPAR-eEPSC | WT | 7 | 0.91 ± 0.04 | p=0.0012 | ** |
| | | | Mut | 6 | 0.44 ± 0.07 | | |

| Fig. 4e | Assay Performed | Parameter | Groups | N (neurons) | Descriptive Statistics | Statistical Analysis | | Column statistics | | |
|---|---|---|---|---|---|---|---|---|---|---|
| | | | | | | One-Way ANOVA | | Bonferroni's multiple comparisons | | |
| | | | | | Mean ± SEM | Statistical Test | Significance | Comparison | P value | Significance |
| | Calcium imaging | GluA1 constructs | A1-WT | 12 | 0.10 ± 0.02 | Treatment: F (5, 52) = 35.82, P<0.0001 | | A1-WT vs. A1-A363T | <0.0001 | *** |
| | | | A1-A636T | 13 | 4.17 ± 0.44 | | | A1-A636T vs. A1-A636T+NBQX | <0.0001 | *** |
| | | | A1-A636T+NBQX | 5 | 0.21 ± 0.04 | | | A2/A1-WT vs A2/A1-A636T | <0.0001 | *** |
| | | GluA2/GluA1 Constructs | A2/A1-WT | 10 | 0.10 ± 0.03 | | | A2/A1-A636T vs. A2/A1-A636T+NBQX | <0.0001 | *** |
| | | | A2/A1-A636T | 13 | 4.31 ± 0.51 | | | A1-WT vs. A2/A1-WT | >0.9999 | ns |
| | | | A2/A1-A636T+NBQX | 5 | 0.14 ± 0.02 | | | A1-A636T vs. A2/A1-A636T | >0.9999 | ns |

| Fig. 4g | Assay Performed | Parameter | Groups | N (wells) | Descriptive Statistics | Statistical Analysis | | Column statistics | | |
|---|---|---|---|---|---|---|---|---|---|---|
| | | | | | | One-Way ANOVA | | Bonferroni's multiple comparisons | | |
| | | | | | Mean ± SEM | Statistical Test | Significance | Comparison | P value | Significance |
| | Excitotoxicity | Venus only | Venus only | 4 | 6.25 ± 0.49 | Treatment: F (11, 41) = 29.64, P<0.0001 | | Venus vs. A1-WT | >0.9999 | ns |
| | | GluA1 constructs | A1-WT | 6 | 11.59 ± 1.34 | | | A1-WT vs. A1-A363T | <0.0001 | *** |
| | | | A1-A636T | 7 | 64.52 ± 5.34 | | | A1-A636T vs. A1-A636T+NBQX | <0.0001 | *** |
| | | | A1-A636T+NBQX | 4 | 18.53 ± 4.15 | | | A2/A1-WT vs A2/A1-A636T | <0.0001 | *** |
| | | GluA2/GluA1 Constructs | A2/A1-WT | 6 | 8.08 ± 2.21 | | | A2/A1-A636T vs. A2/A1-A636T+NBQX | <0.0001 | *** |
| | | | A2/A1-A636T | 7 | 7.32 ± 1.26 | | | A1-WT vs. A2/A1-WT | >0.9999 | ns |
| | | | A2/A1-A636T+NBQX | 4 | 13.57 ± 2.22 | | | A1-A636T vs. A2/A1-A636T | >0.9999 | ns |

| Figure 4k | Assay Performed | Parameter | Groups | N (animals) | Descriptive Statistics | Statistical Analysis | | Column statistics | |
|---|---|---|---|---|---|---|---|---|---|
| | | | | | | Two-Way ANOVA | | Bonferroni's multiple comparisons | |
| | | | | | Mean ± SEM | Statistical Test | Significance | P value | Significance |
| | TUNEL staining | CA1 | WT | 7 | 12.87 ± 2.52 | Interaction: F (2, 30) = 0.5295, P=0.5943<br>Genotype: F (1, 30) = 27.29, P<0.0001<br>Subregion: F (2, 30) = 0.4782, P=0.6245 | | 0.0144 | * |
| | | | Mut | 5 | 75.44 ± 15.76 | | | | |
| | | CA3 | WT | 7 | 16.74 ± 2.83 | | | 0.0024 | ** |
| | | | Mut | 5 | 93.28 ± 28.87 | | | | |
| | | DG | WT | 7 | 34.17 ± 10.14 | | | 0.091 | ns |
| | | | Mut | 5 | 80.85 ± 22.75 | | | | |

| Figure 4l | Assay Performed | Parameter | Groups | N (animals) | Descriptive Statistics | Statistical Analysis | | Column statistics | |
|---|---|---|---|---|---|---|---|---|---|
| | | | | | | Two-Way ANOVA | | Bonferroni's multiple comparisons | |
| | | | | | Mean ± SEM | Statistical Test | Significance | P value | Significance |
| | GFAP staining | CA1 | WT | 4 | 58.05 ± 5.18 | Interaction: F (2, 18) = 8.889, P=0.0021<br>Genotype: F (1, 18) = 95.68, P<0.0001<br>Subregion: F (2, 18) = 11.16, P=0.0007 | | <0.0001 | **** |
| | | | Mut | 4 | 980.54 ± 77.74 | | | | |
| | | CA3 | WT | 4 | 75.73 ± 21.06 | | | <0.0001 | **** |
| | | | Mut | 4 | 929.94 ± 185.68 | | | | |
| | | DG | WT | 4 | 28.23 ± 3.32 | | | 0.1184 | ns |
| | | | Mut | 4 | 296.19 ± 51.89 | | | | |

| Figure 4m | Assay Performed | Parameter | Groups | N (animals) | Descriptive Statistics | Statistical Analysis | | Column statistics | |
|---|---|---|---|---|---|---|---|---|---|
| | | | | | | Two-Way ANOVA | | Bonferroni's multiple comparisons | |
| | | | | | Mean ± SEM | Statistical Test | Significance | P value | Significance |
| | Iba1 staining | CA1 | WT | 4 | 59.47 ± 9.86 | Interaction: F (2, 18) = 37.10, P<0.0001<br>Genotype: F (1, 18) = 215.1, P<0.0001<br>Subregion: F (2, 18) = 37.27, P<0.0001 | | <0.0001 | **** |
| | | | Mut | 4 | 402.29 ± 36.28 | | | | |
| | | CA3 | WT | 4 | 50.40 ± 4.61 | | | <0.0001 | **** |
| | | | Mut | 4 | 693.37 ± 43.55 | | | | |
| | | DG | WT | 4 | 49.12 ± 5.97 | | | 0.0459 | * |
| | | | Mut | 4 | 165.35 ± 47.73 | | | | |

| Figure 5b | Assay Performed | Parameter | Groups | N (animals) | Descriptive Statistics | Statistical Analysis | |
|---|---|---|---|---|---|---|---|
| | | | | | | Mann Whitney test - Two-tailed | |
| | | | | | Mean ± SEM | P value | Significance |
| | qRT-PCR | Mut mRNA | WT | 6 | 0.0006 ± 0.0000 | p=0.0022 | ** |
| | | | Mut | 6 | 1.0030 ± 0.0314 | | |

| Figure 5c | Assay Performed | Parameter | Groups | N (animals) | Descriptive Statistics | Statistical Analysis | |
|---|---|---|---|---|---|---|---|
| | | | | | | Mann Whitney test - Two-tailed | |
| | | | | | Mean ± SEM | P value | Significance |
| | qRT-PCR | WT mRNA | WT | 6 | 1.007 ± 0.0502 | p=0.0022 | ** |
| | | | Mut | 6 | 0.4314 ± 0.0148 | | |

| Figure 5d | Assay Performed | Parameter | Groups | N (animals) | Descriptive Statistics | Statistical Analysis | | Column statistics | | |
|---|---|---|---|---|---|---|---|---|---|---|
| | | | | | | One-Way ANOVA | | Bonferroni's multiple comparisons | | |
| | | | | | Mean ± SEM | Statistical Test | Significance | Comparison | P value | Significance |
| | qRT-PCR | Mut mRNA | VEH | 6 | 1.00 ± 0.03 | Treatment: F (2, 15) = 203.7, P<0.0001 | | VEH vs. ASO2 | <0.0001 | **** |
| | | | ASO2 | 6 | 0.39 ± 0.04 | | | VEH vs. ASO2F | <0.0001 | **** |
| | | | $ASO2^{mC}$ | 6 | 0.26 ± 0.02 | | | ASO2 vs. ASO2F | 0.0162 | * |

| Figure 5e | Assay Performed | Parameter | Groups | N (animals) | Descriptive Statistics | Statistical Analysis | | Column statistics | | |
|---|---|---|---|---|---|---|---|---|---|---|
| | | | | | | One-Way ANOVA | | Bonferroni's multiple comparisons | | |
| | | | | | Mean ± SEM | Statistical Test | Significance | Comparison | P value | Significance |
| | qRT-PCR | WT mRNA | Veh | 6 | 1.01 ± 0.05 | Treatment: F (2, 15) = 1.365, P=0.2854 | | Veh vs. ASO2 | >0.9999 | ns |
| | | | ASO2 | 6 | 1.01 ± 0.06 | | | Veh vs. ASO2F | 0.5179 | ns |
| | | | $ASO2^{mC}$ | 6 | 1.11 ± 0.04 | | | ASO2 vs. ASO2F | 0.5199 | ns |

| Figure 5g | Assay Performed | Parameter | Groups | N (animals) | Descriptive Statistics | Statistical Analysis | | Column statistics | | |
|---|---|---|---|---|---|---|---|---|---|---|
| | | | | | | Two-Way ANOVA | | Bonferroni's multiple comparisons | | |
| | | | | | Mean ± SEM | Statistical Test | Significance | Comparison | P value | Significance |
| | TUNEL staining | WT | Veh | 4 | 10.71 ± 5.44 | Interaction: F (1, 12) = 15.03, P=0.0022 | | WT Veh vs. Het Veh | 0.0005 | *** |
| | | | ASO | 4 | 11.36 ± 1.55 | Treatment: F (1, 12) = 14.60, P= 0.0024 | | WT Veh vs. WT ASO | >0.9999 | ns |
| | | Mut | Veh | 4 | 107.2 ± 22.67 | Genotype: F (1, 12) = 18.43, P=0.0010 | | WT Veh vs. Het ASO | >0.9999 | ns |
| | | | ASO | 4 | 16.29 ± 3.45 | | | Het Veh vs. Het ASO | 0.0009 | *** |

| Figure 5i | Assay Performed | Parameter | Groups | N (animals) | Descriptive Statistics | Statistical Analysis | | Column statistics | | |
|---|---|---|---|---|---|---|---|---|---|---|
| | | | | | | Two-Way ANOVA | | Bonferroni's multiple comparisons | | |
| | | | | | Mean ± SEM | Statistical Test | Significance | Comparison | P value | Significance |
| | GFAP staining | WT | Veh | 4 | 53.13 ± 9.12 | Interaction: F (1, 12) = 27.81, P=0.0002 | | WT Veh vs. Het Veh | <0.0001 | **** |
| | | | ASO | 4 | 41.97 ± 8.71 | Treatment: F (1, 12) = 30.63, P= 0.0001 | | WT Veh vs. WT ASO | >0.9999 | ns |
| | | Mut | Veh | 4 | 530.98 ± 82.78 | Genotype: F (1, 12) = 35.18, P<0.0001 | | WT Veh vs. Het ASO | >0.9999 | ns |
| | | | ASO | 4 | 70.02 ± 16.29 | | | Het Veh vs. Het ASO | <0.0001 | **** |

| Figure 5k | Assay Performed | Parameter | Groups | N (animals) | Descriptive Statistics | Statistical Analysis | | Column statistics | | |
|---|---|---|---|---|---|---|---|---|---|---|
| | | | | | | Two-Way ANOVA | | Bonferroni's multiple comparisons | | |
| | | | | | Mean ± SEM | Statistical Test | Significance | Comparison | P value | Significance |
| | Iba1 staining | WT | Veh | 4 | 60.55 ± 15.25 | Interaction: F (1, 12) = 21.20, P=0.0006 | | WT Veh vs. Het Veh | <0.0001 | **** |
| | | | ASO | 4 | 44.32 ± 21.47 | Treatment: F (1, 12) = 24.90, P= 0.0003 | | WT Veh vs. WT ASO | >0.9999 | ns |
| | | Mut | Veh | 4 | 477.96 ± 75.21 | Genotype: F (1, 12) = 28.28, P<0.0002 | | WT Veh vs. Het ASO | >0.9999 | ns |
| | | | ASO | 4 | 74.33 ± 26.99 | | | Het Veh vs. Het ASO | 0.0001 | *** |

| Figure 5m | Assay Performed | Parameter | Groups | N (animals) | Descriptive Statistics | Statistical Analysis | | Column statistics | | |
|---|---|---|---|---|---|---|---|---|---|---|
| | | | | | | Two-Way ANOVA | | Bonferroni's multiple comparisons | | |
| | | | | | Mean ± SEM | Statistical Test | Significance | Comparison | P value | Significance |
| | Sholl Analysis- Basal dendrites (3 wks) | Distance- 0 | WT Veh | 5 | 0.00 ± 0.00 | | | WT Veh vs. Het Veh | na | na |
| | | | WT ASO | 5 | 0.00 ± 0.00 | | | WT Veh vs. WT ASO | na | na |
| | | | Mut Veh | 5 | 0.00 ± 0.00 | | | WT Veh vs. Het ASO | na | na |
| | | | Mut ASO | 5 | 0.00 ± 0.00 | | | Het Veh vs. Het ASO | na | na |
| | | Distance- 10 | WT Veh | 5 | 5.20 ± 0.37 | | | WT Veh vs. Het Veh | >0.9999 | ns |
| | | | WT ASO | 5 | 5.12 ± 0.29 | | | WT Veh vs. WT ASO | >0.9999 | ns |
| | | | Mut Veh | 5 | 5.60 ± 0.84 | | | WT Veh vs. Het ASO | >0.9999 | ns |
| | | | Mut ASO | 5 | 5.68 ± 0.29 | | | Het Veh vs. Het ASO | >0.9999 | ns |
| | | Distance- 20 | WT Veh | 5 | 10.04 ± 0.87 | | | WT Veh vs. Het Veh | >0.9999 | ns |
| | | | WT ASO | 5 | 9.44 ± 0.55 | | | WT Veh vs. WT ASO | >0.9999 | ns |
| | | | Mut Veh | 5 | 11.36 ± 1.06 | | | WT Veh vs. Het ASO | >0.9999 | ns |
| | | | Mut ASO | 5 | 9.96 ± 0.87 | | | Het Veh vs. Het ASO | >0.9999 | ns |
| | | Distance- 30 | WT Veh | 5 | 15.52 ± 0.95 | | | WT Veh vs. Het Veh | >0.9999 | ns |
| | | | WT ASO | 5 | 14.64 ± 0.66 | | | WT Veh vs. WT ASO | >0.9999 | ns |
| | | | Mut Veh | 5 | 16.60 ± 1.62 | | | WT Veh vs. Het ASO | >0.9999 | ns |
| | | | Mut ASO | 5 | 14.36 ± 0.63 | | | Het Veh vs. Het ASO | >0.9999 | ns |
| | | Distance- 40 | WT Veh | 5 | 18.76 ± 1.04 | | | WT Veh vs. Het Veh | >0.9999 | ns |
| | | | WT ASO | 5 | 17.60 ± 0.51 | | | WT Veh vs. WT ASO | >0.9999 | ns |
| | | | Mut Veh | 5 | 17.60 ± 1.31 | | | WT Veh vs. Het ASO | >0.9999 | ns |
| | | | Mut ASO | 5 | 18.24 ± 0.95 | | | Het Veh vs. Het ASO | >0.9999 | ns |
| | | Distance- 50 | WT Veh | 5 | 20.52 ± 0.96 | | | WT Veh vs. Het Veh | 0.2253 | ns |
| | | | WT ASO | 5 | 19.84 ± 0.48 | | | WT Veh vs. WT ASO | >0.9999 | ns |
| | | | Mut Veh | 5 | 16.64 ± 1.21 | | | WT Veh vs. Het ASO | 0.8826 | ns |
| | | | Mut ASO | 5 | 18.40 ± 0.90 | | | Het Veh vs. Het ASO | >0.9999 | ns |
| | | Distance- 60 | WT Veh | 5 | 21.20 ± 1.02 | | | WT Veh vs. Het Veh | 0.0148 | * |
| | | | WT ASO | 5 | 19.64 ± 0.43 | | | WT Veh vs. WT ASO | >0.9999 | ns |
| | | | Mut Veh | 5 | 12.56 ± 1.56 | | | WT Veh vs. Het ASO | >0.9999 | 0.5398 |
| | | | Mut ASO | 5 | 18.04 ± 1.27 | | | Het Veh vs. Het ASO | >0.9999 | 0.1612 |
| | | Distance- 70 | WT Veh | 5 | 20.00 ± 0.75 | | | WT Veh vs. Het Veh | 0.0037 | ** |
| | | | WT ASO | 5 | 19.28 ± 0.42 | | | WT Veh vs. WT ASO | >0.9999 | ns |
| | | | Mut Veh | 5 | 8.72 ± 1.52 | | | WT Veh vs. Het ASO | 0.3816 | ns |
| | | | Mut ASO | 5 | 16.40 ± 1.40 | | | Het Veh vs. Het ASO | 0.0358 | * |
| | | Distance- 80 | WT Veh | 5 | 18.48 ± 0.42 | | | WT Veh vs. Het Veh | 0.0045 | ** |
| | | | WT ASO | 5 | 17.64 ± 0.73 | | | WT Veh vs. WT ASO | >0.9999 | ns |
| | | | Mut Veh | 5 | 5.28 ± 1.59 | | | WT Veh vs. Het ASO | 0.2495 | ns |
| | | | Mut ASO | 5 | 14.84 ± 1.26 | | | Het Veh vs. Het ASO | 0.0106 | * |
| | | Distance- 90 | WT Veh | 5 | 15.64 ± 0.51 | | | WT Veh vs. Het Veh | 0.0023 | ** |
| | | | WT ASO | 5 | 16.24 ± 0.69 | | | WT Veh vs. WT ASO | >0.9999 | ns |
| | | | Mut Veh | 5 | 2.84 ± 1.43 | | | WT Veh vs. Het ASO | >0.9999 | ns |
| | | | Mut ASO | 5 | 11.64 ± 1.36 | | | Het Veh vs. Het ASO | 0.0127 | * |
| | | Distance- 100 | WT Veh | 5 | 12.04 ± 0.71 | Interaction: F (60, 320) = 11.93, P<0.0001 | | WT Veh vs. Het Veh | 0.0020 | ** |
| | | | WT ASO | 5 | 13.12 ± 0.62 | Distance: F (3.020, 48.32) = 435.3, P<0.0001 | | WT Veh vs. WT ASO | >0.9999 | ns |
| | | | Mut Veh | 5 | 1.80 ± 1.26 | Genotype+treatment:F (3, 16) = 22.90, P<0.0001 | | WT Veh vs. Het ASO | >0.9999 | ns |
| | | | Mut ASO | 5 | 9.80 ± 1.53 | | | Het Veh vs. Het ASO | 0.0242 | * |
| | | Distance- 110 | WT Veh | 5 | 9.20 ± 0.66 | | | WT Veh vs. Het Veh | 0.0003 | *** |
| | | | WT ASO | 5 | 9.28 ± 0.75 | | | WT Veh vs. WT ASO | >0.9999 | ns |
| | | | Mut Veh | 5 | 1.08 ± 0.79 | | | WT Veh vs. Het ASO | 0.4298 | ns |
| | | | Mut ASO | 5 | 6.56 ± 1.05 | | | Het Veh vs. Het ASO | 0.0217 | * |
| | | Distance- 120 | WT Veh | 5 | 5.64 ± 0.73 | | | WT Veh vs. Het Veh | >0.9999 | ns |
| | | | WT ASO | 5 | 5.80 ± 0.75 | | | WT Veh vs. WT ASO | 0.0045 | ** |
| | | | Mut Veh | 5 | 0.64 ± 0.55 | | | WT Veh vs. Het ASO | >0.9999 | ns |
| | | | Mut ASO | 5 | 4.12 ± 0.83 | | | Het Veh vs. Het ASO | 0.0612 | ns |
| | | Distance- 130 | WT Veh | 5 | 2.84 ± 0.45 | | | WT Veh vs. Het Veh | 0.0154 | * |
| | | | WT ASO | 5 | 3.52 ± 0.83 | | | WT Veh vs. WT ASO | >0.9999 | ns |
| | | | Mut Veh | 5 | 0.24 ± 0.19 | | | WT Veh vs. Het ASO | >0.9999 | ns |

| | | | | | | | | | |
|---|---|---|---|---|---|---|---|---|---|
| | | Mut ASO | 5 | 2.44 ± 0.50 | | | Het Veh vs. Het ASO | 0.05 | ns |
| | Distance- 140 | WT Veh | 5 | 1.40 ± 0.18 | | | WT Veh vs. Het Veh | 0.0051 | ** |
| | | WT ASO | 5 | 2.20 ± 0.86 | | | WT Veh vs. WT ASO | >0.9999 | ns |
| | | Mut Veh | 5 | 0.12 ± 0.08 | | | WT Veh vs. Het ASO | >0.9999 | ns |
| | | Mut ASO | 5 | 1.32 ± 0.36 | | | Het Veh vs. Het ASO | 0.1653 | ns |
| | Distance- 150 | WT Veh | 5 | 0.28 ± 0.08 | | | WT Veh vs. Het Veh | 0.2225 | ns |
| | | WT ASO | 5 | 0.92 ± 0.47 | | | WT Veh vs. WT ASO | >0.9999 | ns |
| | | Mut Veh | 5 | 0.04 ± 0.04 | | | WT Veh vs. Het ASO | >0.9999 | ns |
| | | Mut ASO | 5 | 0.72 ± 0.29 | | | Het Veh vs. Het ASO | 0.4877 | ns |
| | Distance- 160 | WT Veh | 5 | 0.08 ± 0.05 | | | WT Veh vs. Het Veh | >0.9999 | ns |
| | | WT ASO | 5 | 0.32 ± 0.22 | | | WT Veh vs. WT ASO | >0.9999 | ns |
| | | Mut Veh | 5 | 0.00 ± 0.00 | | | WT Veh vs. Het ASO | >0.9999 | ns |
| | | Mut ASO | 5 | 0.44 ± 0.25 | | | Het Veh vs. Het ASO | 0.9057 | ns |
| | Distance- 170 | WT Veh | 5 | 0.04 ± 0.04 | | | WT Veh vs. Het Veh | >0.9999 | ns |
| | | WT ASO | 5 | 0.12 ± 0.12 | | | WT Veh vs. WT ASO | >0.9999 | ns |
| | | Mut Veh | 5 | 0.00 ± 0.00 | | | WT Veh vs. Het ASO | 0.9992 | ns |
| | | Mut ASO | 5 | 0.24 ± 0.12 | | | Het Veh vs. Het ASO | 0.6522 | ns |
| | Distance- 180 | WT Veh | 5 | 0.00 ± 0.00 | | | WT Veh vs. Het Veh | na | na |
| | | WT ASO | 5 | 0.04 ± 0.04 | | | WT Veh vs. WT ASO | >0.9999 | ns |
| | | Mut Veh | 5 | 0.00 ± 0.00 | | | WT Veh vs. Het ASO | 0.5958 | ns |
| | | Mut ASO | 5 | 0.16 ± 0.07 | | | Het Veh vs. Het ASO | 0.5958 | ns |
| | Distance- 190 | WT Veh | 5 | 0.00 ± 0.00 | | | WT Veh vs. Het Veh | na | na |
| | | WT ASO | 5 | 0.00 ± 0.00 | | | WT Veh vs. WT ASO | >0.9999 | ns |
| | | Mut Veh | 5 | 0.00 ± 0.00 | | | WT Veh vs. Het ASO | >0.9999 | ns |
| | | Mut ASO | 5 | 0.12 ± 0.08 | | | Het Veh vs. Het ASO | >0.9999 | ns |
| | Distance- 200 | WT Veh | 5 | 0.00 ± 0.00 | | | WT Veh vs. Het Veh | na | na |
| | | WT ASO | 5 | 0.00 ± 0.00 | | | WT Veh vs. WT ASO | na | na |
| | | Mut Veh | 5 | 0.00 ± 0.00 | | | WT Veh vs. Het ASO | >0.9999 | ns |
| | | Mut ASO | 5 | 0.08 ± 0.05 | | | Het Veh vs. Het ASO | >0.9999 | ns |

| Assay Performed | Parameter | Groups | N (animals) | Descriptive Statistics | Statistical Analysis | | Column statistics | | |
|---|---|---|---|---|---|---|---|---|---|
| | | | | | Two-Way ANOVA | | Bonferroni's multiple comparisons | | |
| | | | | Mean ± SEM | Statistical Test | Significance | Comparison | P value | Significance |
| | Distance- 0 | WT Veh | 5 | 0.00 ± 0.00 | | | WT Veh vs. Het Veh | na | na |
| | | WT ASO | 5 | 0.00 ± 0.00 | | | WT Veh vs. WT ASO | na | na |
| | | Mut Veh | 5 | 0.00 ± 0.00 | | | WT Veh vs. Het ASO | na | na |
| | | Mut ASO | 5 | 0.00 ± 0.00 | | | Het Veh vs. Het ASO | na | na |
| | Distance- 10 | WT Veh | 5 | 1.04 ± 0.07 | | | WT Veh vs. Het Veh | >0.9999 | ns |
| | | WT ASO | 5 | 0.92 ± 0.08 | | | WT Veh vs. WT ASO | >0.9999 | ns |
| | | Mut Veh | 5 | 1.04 ± 0.04 | | | WT Veh vs. Het ASO | >0.9999 | ns |
| | | Mut ASO | 5 | 1.28 ± 0.19 | | | Het Veh vs. Het ASO | >0.9999 | ns |
| | Distance- 20 | WT Veh | 5 | 1.52 ± 0.19 | | | WT Veh vs. Het Veh | >0.9999 | ns |
| | | WT ASO | 5 | 1.20 ± 0.18 | | | WT Veh vs. WT ASO | >0.9999 | ns |
| | | Mut Veh | 5 | 1.60 ± 0.30 | | | WT Veh vs. Het ASO | >0.9999 | ns |
| | | Mut ASO | 5 | 1.80 ± 0.43 | | | Het Veh vs. Het ASO | >0.9999 | ns |
| | Distance- 30 | WT Veh | 5 | 1.84 ± 0.19 | | | WT Veh vs. Het Veh | 0.7466 | na |
| | | WT ASO | 5 | 1.56 ± 0.30 | | | WT Veh vs. WT ASO | >0.9999 | ns |
| | | Mut Veh | 5 | 3.96 ± 1.09 | | | WT Veh vs. Het ASO | >0.9999 | ns |
| | | Mut ASO | 5 | 2.16 ± 0.54 | | | Het Veh vs. Het ASO | >0.9999 | ns |
| | Distance- 40 | WT Veh | 5 | 2.92 ± 0.27 | | | WT Veh vs. Het Veh | 0.0445 | * |
| | | WT ASO | 5 | 2.72 ± 0.49 | | | WT Veh vs. WT ASO | >0.9999 | ns |
| | | Mut Veh | 5 | 7.60 ± 0.99 | | | WT Veh vs. Het ASO | >0.9999 | ns |
| | | Mut ASO | 5 | 2.64 ± 0.55 | | | Het Veh vs. Het ASO | 0.0254 | * |
| | Distance- 50 | WT Veh | 5 | 4.16 ± 0.79 | | | WT Veh vs. Het Veh | 0.0516 | ns |
| | | WT ASO | 5 | 3.80 ± 0.55 | | | WT Veh vs. WT ASO | >0.9999 | ns |
| | | Mut Veh | 5 | 9.72 ± 1.29 | | | WT Veh vs. Het ASO | >0.9999 | ns |
| | | Mut ASO | 5 | 4.00 ± 0.72 | | | Het Veh vs. Het ASO | 0.0448 | * |
| | Distance- 60 | WT Veh | 5 | 6.44 ± 0.99 | | | WT Veh vs. Het Veh | 0.2299 | ns |
| | | WT ASO | 5 | 5.56 ± 1.07 | | | WT Veh vs. WT ASO | >0.9999 | ns |
| | | Mut Veh | 5 | 10.76 ± 1.39 | | | WT Veh vs. Het ASO | >0.9999 | ns |
| | | Mut ASO | 5 | 5.92 ± 1.06 | | | Het Veh vs. Het ASO | 0.1565 | ns |
| | Distance- 70 | WT Veh | 5 | 8.44 ± 1.11 | | | WT Veh vs. Het Veh | 0.9788 | ns |
| | | WT ASO | 5 | 6.76 ± 1.04 | | | WT Veh vs. WT ASO | >0.9999 | ns |
| | | Mut Veh | 5 | 11.24 ± 1.43 | | | WT Veh vs. Het ASO | >0.9999 | ns |
| | | Mut ASO | 5 | 7.80 ± 1.36 | | | Het Veh vs. Het ASO | 0.7201 | ns |
| | Distance- 80 | WT Veh | 5 | 10.60 ± 1.43 | | | WT Veh vs. Het Veh | >0.9999 | ns |
| | | WT ASO | 5 | 8.04 ± 1.34 | | | WT Veh vs. WT ASO | >0.9999 | ns |
| | | Mut Veh | 5 | 10.60 ± 1.16 | | | WT Veh vs. Het ASO | >0.9999 | ns |
| | | Mut ASO | 5 | 8.80 ± 1.42 | | | Het Veh vs. Het ASO | >0.9999 | ns |
| | Distance- 90 | WT Veh | 5 | 11.32 ± 0.93 | | | WT Veh vs. Het Veh | >0.9999 | ns |
| | | WT ASO | 5 | 8.72 ± 0.96 | | | WT Veh vs. WT ASO | >0.9999 | ns |
| | | Mut Veh | 5 | 10.12 ± 1.30 | | | WT Veh vs. Het ASO | >0.9999 | ns |
| | | Mut ASO | 5 | 10.48 ± 1.45 | | | Het Veh vs. Het ASO | >0.9999 | ns |
| | Distance- 100 | WT Veh | 5 | 11.88 ± 0.80 | | | WT Veh vs. Het Veh | 0.5943 | ns |
| | | WT ASO | 5 | 9.00 ± 0.50 | | | WT Veh vs. WT ASO | 0.119 | ns |
| | | Mut Veh | 5 | 9.36 ± 1.07 | | | WT Veh vs. Het ASO | >0.9999 | ns |
| | | Mut ASO | 5 | 11.12 ± 1.02 | | | Het Veh vs. Het ASO | >0.9999 | ns |
| | Distance- 110 | WT Veh | 5 | 11.08 ± 0.70 | | | WT Veh vs. Het Veh | 0.0724 | ns |
| | | WT ASO | 5 | 9.48 ± 0.46 | | | WT Veh vs. WT ASO | 0.5891 | ns |
| | | Mut Veh | 5 | 7.12 ± 0.97 | | | WT Veh vs. Het ASO | >0.9999 | ns |
| | | Mut ASO | 5 | 10.60 ± 0.72 | | | Het Veh vs. Het ASO | 0.1316 | ns |
| | Distance- 120 | WT Veh | 5 | 10.64 ± 0.63 | | | WT Veh vs. Het Veh | 0.0264 | * |
| | | WT ASO | 5 | 10.08 ± 0.33 | | | WT Veh vs. WT ASO | >0.9999 | ns |
| | | Mut Veh | 5 | 6.32 ± 0.86 | | | WT Veh vs. Het ASO | >0.9999 | ns |
| | | Mut ASO | 5 | 9.44 ± 0.97 | | | Het Veh vs. Het ASO | 0.2576 | ns |
| | Distance- 130 | WT Veh | 5 | 9.76 ± 0.93 | | | WT Veh vs. Het Veh | 0.0321 | * |
| | | WT ASO | 5 | 10.24 ± 0.39 | | | WT Veh vs. WT ASO | >0.9999 | ns |
| | | Mut Veh | 5 | 4.96 ± 0.85 | | | WT Veh vs. Het ASO | >0.9999 | ns |
| | | Mut ASO | 5 | 8.80 ± 0.64 | | | Het Veh vs. Het ASO | 0.0479 | * |
| | Distance- 140 | WT Veh | 5 | 8.72 ± 0.56 | | | WT Veh vs. Het Veh | 0.0095 | ** |
| | | WT ASO | 5 | 10.44 ± 0.56 | | | WT Veh vs. WT ASO | 0.3627 | ns |
| | | Mut Veh | 5 | 3.88 ± 0.80 | | | WT Veh vs. Het ASO | >0.9999 | ns |
| | | Mut ASO | 5 | 8.84 ± 0.60 | | | Het Veh vs. Het ASO | 0.0086 | ** |
| | Distance- 150 | WT Veh | 5 | 8.76 ± 0.17 | | | WT Veh vs. Het Veh | 0.0003 | *** |
| | | WT ASO | 5 | 9.56 ± 0.70 | | | WT Veh vs. WT ASO | >0.9999 | ns |
| | | Mut Veh | 5 | 3.52 ± 0.41 | | | WT Veh vs. Het ASO | >0.9999 | ns |
| | | Mut ASO | 5 | 8.28 ± 0.57 | | | Het Veh vs. Het ASO | 0.0014 | ** |
| | Distance- 160 | WT Veh | 5 | 8.68 ± 0.21 | | | WT Veh vs. Het Veh | <0.0001 | **** |
| | | WT ASO | 5 | 9.24 ± 0.87 | | | WT Veh vs. WT ASO | >0.9999 | ns |
| | | Mut Veh | 5 | 2.80 ± 0.36 | | | WT Veh vs. Het ASO | >0.9999 | ns |
| | | Mut ASO | 5 | 8.64 ± 0.44 | | | Het Veh vs. Het ASO | <0.0001 | **** |
| | Distance- 170 | WT Veh | 5 | 7.92 ± 0.15 | | | WT Veh vs. Het Veh | <0.0001 | **** |
| | | WT ASO | 5 | 8.96 ± 0.53 | | | WT Veh vs. WT ASO | >0.9999 | ns |
| | | Mut Veh | 5 | 2.64 ± 0.17 | | | WT Veh vs. Het ASO | >0.9999 | ns |
| | | Mut ASO | 5 | 8.16 ± 0.45 | | | Het Veh vs. Het ASO | 0.0004 | *** |
| | Distance- 180 | WT Veh | 5 | 7.24 ± 0.28 | | | WT Veh vs. Het Veh | <0.0001 | **** |
| | | WT ASO | 5 | 8.24 ± 0.69 | | | WT Veh vs. WT ASO | 0.7231 | ns |
| | | Mut Veh | 5 | 2.60 ± 0.14 | | | WT Veh vs. Het ASO | >0.9999 | ns |
| | | Mut ASO | 5 | 7.40 ± 0.59 | | | Het Veh vs. Het ASO | 0.0054 | ** |
| | Distance- 190 | WT Veh | 5 | 6.80 ± 0.28 | | | WT Veh vs. Het Veh | <0.0001 | **** |
| | | WT ASO | 5 | 7.84 ± 0.45 | | | WT Veh vs. WT ASO | 0.56 | ns |
| | | Mut Veh | 5 | 2.20 ± 0.13 | | | WT Veh vs. Het ASO | >0.9999 | ns |
| | | Mut ASO | 5 | 7.52 ± 0.65 | | | Het Veh vs. Het ASO | 0.0057 | ** |
| | Distance- 200 | WT Veh | 5 | 6.60 ± 0.26 | | | WT Veh vs. Het Veh | <0.0001 | **** |
| | | WT ASO | 5 | 6.76 ± 0.58 | | | WT Veh vs. WT ASO | >0.9999 | ns |

| Parameter | Groups | N (animals) | Mean ± SEM | Statistical Analysis | Comparison | P value | Significance |
|---|---|---|---|---|---|---|---|
| Distance- 200 | Mut Veh | 5 | 1.88 ± 0.19 | | WT Veh vs. Het ASO | >0.9999 | ns |
| | Mut ASO | 5 | 7.04 ± 0.59 | | Het Veh vs. Het ASO | 0.0031 | ** |
| Distance- 210 | WT Veh | 5 | 6.40 ± 0.53 | | WT Veh vs. Het Veh | 0.0017 | ** |
| | WT ASO | 5 | 5.92 ± 0.77 | | WT Veh vs. WT ASO | >0.9999 | ns |
| | Mut Veh | 5 | 1.88 ± 0.26 | | WT Veh vs. Het ASO | >0.9999 | ns |
| | Mut ASO | 5 | 6.60 ± 0.40 | | Het Veh vs. Het ASO | 0.0002 | *** |
| Distance- 220 | WT Veh | 5 | 5.28 ± 0.61 | | WT Veh vs. Het Veh | 0.0154 | * |
| | WT ASO | 5 | 5.08 ± 0.73 | | WT Veh vs. WT ASO | >0.9999 | ns |
| | Mut Veh | 5 | 1.92 ± 0.43 | | WT Veh vs. Het ASO | >0.9999 | ns |
| | Mut ASO | 5 | 5.52 ± 0.71 | Interaction: F (135, 720) = 6.498, P<0.0001<br>Distance: F (3.911, 62.58) = 126.3, P<0.0001<br>Genotype+treatment: F (3, 16) = 23.03, P<0.0001 | Het Veh vs. Het ASO | 0.0234 | * |
| Distance- 230 | WT Veh | 5 | 4.36 ± 0.35 | | WT Veh vs. Het Veh | 0.0314 | * |
| | WT ASO | 5 | 4.68 ± 0.59 | | WT Veh vs. WT ASO | >0.9999 | ns |
| | Mut Veh | 5 | 1.76 ± 0.54 | | WT Veh vs. Het ASO | >0.9999 | ns |
| | Mut ASO | 5 | 4.72 ± 0.65 | | Het Veh vs. Het ASO | 0.0511 | ns |
| Distance- 240 | WT Veh | 5 | 3.80 ± 0.32 | | WT Veh vs. Het Veh | 0.0653 | ns |
| | WT ASO | 5 | 3.76 ± 0.38 | | WT Veh vs. WT ASO | 0.0653 | ns |
| | Mut Veh | 5 | 1.40 ± 0.59 | | WT Veh vs. Het ASO | >0.9999 | ns |
| | Mut ASO | 5 | 3.64 ± 0.55 | | Het Veh vs. Het ASO | 0.1424 | ns |
| Distance- 250 | WT Veh | 5 | 3.40 ± 0.32 | | WT Veh vs. Het Veh | 0.0332 | * |
| | WT ASO | 5 | 3.40 ± 0.47 | | WT Veh vs. WT ASO | >0.9999 | ns |
| | Mut Veh | 5 | 0.96 ± 0.52 | | WT Veh vs. Het ASO | >0.9999 | ns |
| | Mut ASO | 5 | 3.32 ± 0.44 | | Het Veh vs. Het ASO | 0.0509 | ns |
| Distance- 260 | WT Veh | 5 | 3.60 ± 0.40 | | WT Veh vs. Het Veh | 0.0389 | * |
| | WT ASO | 5 | 3.40 ± 0.24 | | WT Veh vs. WT ASO | >0.9999 | ns |
| | Mut Veh | 5 | 0.88 ± 0.59 | | WT Veh vs. Het ASO | >0.9999 | ns |
| | Mut ASO | 5 | 3.00 ± 0.40 | | Het Veh vs. Het ASO | 0.1234 | ns |
| Distance- 270 | WT Veh | 5 | 3.84 ± 0.50 | | WT Veh vs. Het Veh | 0.0078 | ** |
| | WT ASO | 5 | 3.12 ± 0.23 | | WT Veh vs. WT ASO | >0.9999 | ns |
| | Mut Veh | 5 | 0.56 ± 0.46 | | WT Veh vs. Het ASO | >0.9999 | ns |
| | Mut ASO | 5 | 2.96 ± 0.47 | | Het Veh vs. Het ASO | 0.0387 | * |
| Distance- 280 | WT Veh | 5 | 3.72 ± 0.66 | | WT Veh vs. Het Veh | 0.0268 | * |
| | WT ASO | 5 | 2.68 ± 0.15 | | WT Veh vs. WT ASO | >0.9999 | ns |
| | Mut Veh | 5 | 0.44 ± 0.34 | | WT Veh vs. Het ASO | >0.9999 | ns |
| | Mut ASO | 5 | 2.56 ± 0.38 | | Het Veh vs. Het ASO | 0.0202 | * |
| Distance- 290 | WT Veh | 5 | 3.72 ± 0.80 | | WT Veh vs. Het Veh | 0.0653 | ns |
| | WT ASO | 5 | 2.44 ± 0.15 | | WT Veh vs. WT ASO | >0.9999 | ns |
| | Mut Veh | 5 | 0.40 ± 0.30 | | WT Veh vs. Het ASO | >0.9999 | ns |
| | Mut ASO | 5 | 2.40 ± 0.31 | | Het Veh vs. Het ASO | 0.0104 | * |
| Distance- 300 | WT Veh | 5 | 3.48 ± 0.38 | | WT Veh vs. Het Veh | 0.0022 | ** |
| | WT ASO | 5 | 2.24 ± 0.34 | | WT Veh vs. WT ASO | 0.2493 | ns |
| | Mut Veh | 5 | 0.40 ± 0.35 | | WT Veh vs. Het ASO | 0.5539 | ns |
| | Mut ASO | 5 | 2.44 ± 0.39 | | Het Veh vs. Het ASO | 0.0277 | * |
| Distance- 310 | WT Veh | 5 | 3.72 ± 0.29 | | WT Veh vs. Het Veh | 0.0001 | *** |
| | WT ASO | 5 | 2.24 ± 0.29 | | WT Veh vs. WT ASO | 0.439 | * |
| | Mut Veh | 5 | 0.24 ± 0.24 | | WT Veh vs. Het ASO | 0.5328 | ns |
| | Mut ASO | 5 | 2.56 ± 0.50 | | Het Veh vs. Het ASO | 0.038 | * |
| Distance- 320 | WT Veh | 5 | 3.40 ± 0.19 | | WT Veh vs. Het Veh | 0.0002 | *** |
| | WT ASO | 5 | 2.48 ± 0.47 | | WT Veh vs. WT ASO | 0.7631 | ns |
| | Mut Veh | 5 | 0.28 ± 0.28 | | WT Veh vs. Het ASO | 0.4003 | ns |
| | Mut ASO | 5 | 2.36 ± 0.42 | | Het Veh vs. Het ASO | 0.026 | * |
| Distance- 330 | WT Veh | 5 | 3.56 ± 0.56 | | WT Veh vs. Het Veh | 0.0188 | * |
| | WT ASO | 5 | 2.68 ± 0.53 | | WT Veh vs. WT ASO | >0.9999 | ns |
| | Mut Veh | 5 | 0.00 ± 0.00 | | WT Veh vs. Het ASO | 0.6536 | ns |
| | Mut ASO | 5 | 2.24 ± 0.47 | | Het Veh vs. Het ASO | 0.0518 | ns |
| Distance- 340 | WT Veh | 5 | 3.40 ± 0.51 | | WT Veh vs. Het Veh | 0.0158 | * |
| | WT ASO | 5 | 2.56 ± 0.53 | | WT Veh vs. WT ASO | >0.9999 | ns |
| | Mut Veh | 5 | 0.00 ± 0.00 | | WT Veh vs. Het ASO | >0.9999 | ns |
| | Mut ASO | 5 | 2.32 ± 0.60 | | Het Veh vs. Het ASO | 0.1055 | ns |
| Distance- 350 | WT Veh | 5 | 2.60 ± 0.69 | | WT Veh vs. Het Veh | 0.1178 | ns |
| | WT ASO | 5 | 2.40 ± 0.52 | | WT Veh vs. WT ASO | >0.9999 | ns |
| | Mut Veh | 5 | 0.00 ± 0.00 | | WT Veh vs. Het ASO | >0.9999 | ns |
| | Mut ASO | 5 | 1.96 ± 0.54 | | Het Veh vs. Het ASO | 0.1345 | ns |
| Distance- 360 | WT Veh | 5 | 2.12 ± 0.62 | | WT Veh vs. Het Veh | 0.1595 | ns |
| | WT ASO | 5 | 2.16 ± 0.53 | | WT Veh vs. WT ASO | >0.9999 | ns |
| | Mut Veh | 5 | 0.00 ± 0.00 | | WT Veh vs. Het ASO | >0.9999 | ns |
| | Mut ASO | 5 | 1.88 ± 0.45 | | Het Veh vs. Het ASO | 0.0864 | ns |
| Distance- 370 | WT Veh | 5 | 1.28 ± 0.38 | | WT Veh vs. Het Veh | 0.1722 | ns |
| | WT ASO | 5 | 1.40 ± 0.52 | | WT Veh vs. WT ASO | >0.9999 | ns |
| | Mut Veh | 5 | 0.00 ± 0.00 | | WT Veh vs. Het ASO | >0.9999 | ns |
| | Mut ASO | 5 | 1.52 ± 0.43 | | Het Veh vs. Het ASO | 0.1416 | ns |
| Distance- 380 | WT Veh | 5 | 0.88 ± 0.36 | | WT Veh vs. Het Veh | 0.4114 | ns |
| | WT ASO | 5 | 1.16 ± 0.50 | | WT Veh vs. WT ASO | >0.9999 | ns |
| | Mut Veh | 5 | 0.00 ± 0.00 | | WT Veh vs. Het ASO | >0.9999 | ns |
| | Mut ASO | 5 | 1.32 ± 0.43 | | Het Veh vs. Het ASO | 0.2265 | ns |
| Distance- 390 | WT Veh | 5 | 0.68 ± 0.42 | | WT Veh vs. Het Veh | >0.9999 | ns |
| | WT ASO | 5 | 0.88 ± 0.53 | | WT Veh vs. WT ASO | >0.9999 | ns |
| | Mut Veh | 5 | 0.00 ± 0.00 | | WT Veh vs. Het ASO | >0.9999 | ns |
| | Mut ASO | 5 | 0.92 ± 0.37 | | Het Veh vs. Het ASO | 0.3966 | ns |
| Distance- 400 | WT Veh | 5 | 0.52 ± 0.33 | | WT Veh vs. Het Veh | >0.9999 | ns |
| | WT ASO | 5 | 0.96 ± 0.64 | | WT Veh vs. WT ASO | >0.9999 | ns |
| | Mut Veh | 5 | 0.00 ± 0.00 | | WT Veh vs. Het ASO | >0.9999 | ns |
| | Mut ASO | 5 | 0.64 ± 0.31 | | Het Veh vs. Het ASO | 0.6276 | ns |
| Distance- 410 | WT Veh | 5 | 0.28 ± 0.17 | | WT Veh vs. Het Veh | >0.9999 | ns |
| | WT ASO | 5 | 0.88 ± 0.69 | | WT Veh vs. WT ASO | >0.9999 | ns |
| | Mut Veh | 5 | 0.00 ± 0.00 | | WT Veh vs. Het ASO | >0.9999 | ns |
| | Mut ASO | 5 | 0.48 ± 0.23 | | Het Veh vs. Het ASO | 0.6522 | ns |
| Distance- 420 | WT Veh | 5 | 0.20 ± 0.13 | | WT Veh vs. Het Veh | >0.9999 | ns |
| | WT ASO | 5 | 0.84 ± 0.60 | | WT Veh vs. WT ASO | >0.9999 | ns |
| | Mut Veh | 5 | 0.00 ± 0.00 | | WT Veh vs. Het ASO | >0.9999 | ns |
| | Mut ASO | 5 | 0.44 ± 0.22 | | Het Veh vs. Het ASO | 0.7164 | ns |
| Distance- 430 | WT Veh | 5 | 0.08 ± 0.08 | | WT Veh vs. Het Veh | >0.9999 | ns |
| | WT ASO | 5 | 0.68 ± 0.54 | | WT Veh vs. WT ASO | >0.9999 | ns |
| | Mut Veh | 5 | 0.00 ± 0.00 | | WT Veh vs. Het ASO | >0.9999 | ns |
| | Mut ASO | 5 | 0.44 ± 0.22 | | Het Veh vs. Het ASO | 0.7164 | ns |
| Distance- 440 | WT Veh | 5 | 0.04 ± 0.04 | | WT Veh vs. Het Veh | >0.9999 | ns |
| | WT ASO | 5 | 0.32 ± 0.22 | | WT Veh vs. WT ASO | >0.9999 | ns |
| | Mut Veh | 5 | 0.00 ± 0.00 | | WT Veh vs. Het ASO | >0.9999 | ns |
| | Mut ASO | 5 | 0.36 ± 0.22 | | Het Veh vs. Het ASO | >0.9999 | ns |
| Distance- 450 | WT Veh | 5 | 0.04 ± 0.04 | | WT Veh vs. Het Veh | >0.9999 | ns |
| | WT ASO | 5 | 0.04 ± 0.04 | | WT Veh vs. WT ASO | >0.9999 | ns |
| | Mut Veh | 5 | 0.00 ± 0.00 | | WT Veh vs. Het ASO | >0.9999 | ns |
| | Mut ASO | 5 | 0.04 ± 0.04 | | Het Veh vs. Het ASO | >0.9999 | ns |

| Figure 5m | Assay Performed | Parameter | Groups | N (animals) | Descriptive Statistics | Statistical Analysis | |
|---|---|---|---|---|---|---|---|
| | | | | | | Mann Whitney test - Two-tailed | |
| | | | | | Mean ± SEM | P value | Significance |
| | Three-chamber social test<br><br>Sociability | WT Veh | WT-E | 16 | 36.07 ± 5.54 | p<0.0001 | **** |
| | | | WT-M1 | | 108.70 ± 10.74 | | |
| | | WT ASO | Mut-E | 14 | 44.13 ± 6.53 | p=0.0002 | *** |
| | | | Mut-M1 | | 86.07 ± 7.68 | | |
| | | Mut Veh | WT-E | 16 | 50.34 ± 7.58 | p=0.8381 | ns |
| | | | WT-M1 | | 60.84 ± 11.79 | | |
| | | Mut ASO | Mut-E | 16 | 35.72 ± 5.04 | p<0.0001 | **** |
| | | | Mut-M1 | | 110.50 ± 9.56 | | |

| Figure 5o | Assay Performed | Parameter | Groups | N (animals) | Descriptive Statistics<br>Mean ± SEM | Statistical Analysis<br>Mann Whitney test - Two-tailed<br>P value | <br><br>Significance |
|---|---|---|---|---|---|---|---|
| | Three-chamber social test<br><br>Social Novelty | WT Veh | WT-E | 16 | 40.44 ± 4.64 | p=0.0034 | ** |
| | | | WT-M1 | | 77.59 ± 10.82 | | |
| | | WT ASO | Mut-E | 14 | 32.47 ± 5.08 | p<0.0001 | **** |
| | | | Mut-M1 | | 93.64 ± 9.05 | | |
| | | Mut Veh | WT-E | 16 | 48.38 ± 8.64 | p=0.9556 | ns |
| | | | WT-M1 | | 50.60 ± 9.81 | | |
| | | Mut ASO | Mut-E | 16 | 33.47 ± 4.31 | p<0.0001 | **** |
| | | | Mut-M1 | | 84.83 ± 10.05 | | |

| Figure 5p | Assay Performed | Parameter | Groups | N (animals) | Descriptive Statistics<br>Mean ± SEM | Statistical Analysis<br>One-Way ANOVA<br>Statistical Test | <br><br>Significance | Column statistics<br>Bonferroni's multiple comparisons<br>Comparison | <br><br>P value | <br><br>Significance |
|---|---|---|---|---|---|---|---|---|---|---|
| | Y maze test | Discrimination index | WT Veh | 15 | 0.7054 ± 0.03077 | Group (Genotype+treatment): F (3, 56) = 7.884, P=0.0002 | | WT Veh vs. Mut Veh | 0.0004 | *** |
| | | | WT ASO | 15 | 0.6921 ± 0.03702 | | | WT Veh vs. WT ASO | >0.9999 | ns |
| | | | Mut Veh | 14 | 0.4937 ± 0.04132 | | | WT ASO vs. Mut Veh | 0.001 | *** |
| | | | Mut ASO | 16 | 0.6639 ± 0.02745 | | | Mut Veh vs. Mut ASO | 0.0052 | ** |
| | | | | | | One sample t test<br>Theoretical mean 0.5 | <br>Significance | <br>P value | <br>Significance | |
| | | | WT Veh | 15 | 0.7054 ± 0.03077 | t=6.67, df=14 | | <0.0001 | **** | |
| | | | WT ASO | 15 | 0.6921 ± 0.03702 | t=5.19, df=14 | | 0.0001 | *** | |
| | | | Mut Veh | 14 | 0.4937 ± 0.04132 | t=0.15, df=13 | | 0.882 | ns | |
| | | | Mut ASO | 16 | 0.6639 ± 0.02745 | t=5.97, df=15 | | <0.0001 | **** | |

| Figure 5q | Assay Performed | Parameter | Groups | N (animals) | Descriptive Statistics<br>Mean ± SEM | Statistical Analysis<br>Two-way RM ANOVA<br>Statistical Test | <br><br>Significance | Column statistics<br>Bonferroni's multiple comparisons<br>Comparison | <br><br>P value | <br><br>Significance |
|---|---|---|---|---|---|---|---|---|---|---|
| | MWM learning test | Section 1 | WT Veh | 17 | 38.03 ± 3.51 | Interaction: F (19.39, 381.3) = 2.875, P<0.0001<br>Group (Genotype+treatment): F (3, 59) = 91.70, P<0.0001<br>Section: F (6.464, 381.3) = 35.91, P<0.0001 | | WT Veh vs. Mut Veh | >0.9999 | ns |
| | | | WT ASO | 15 | 41.35 ± 2.07 | | | WT Veh vs. WT ASO | >0.9999 | ns |
| | | | Mut Veh | 15 | 43.28 ± 2.61 | | | WT Veh vs. Mut ASO | >0.9999 | ns |
| | | | Mut ASO | 16 | 42.75 ± 2.34 | | | Mut Veh vs. Mut ASO | >0.9999 | ns |
| | | Section 2 | WT Veh | 17 | 28.77 ± 3.07 | | | WT Veh vs. Mut Veh | 0.0196 | * |
| | | | WT ASO | 15 | 32.83 ± 3.08 | | | WT Veh vs. WT ASO | >0.9999 | ns |
| | | | Mut Veh | 15 | 41.36 ± 2.45 | | | WT Veh vs. Mut ASO | >0.9999 | ns |
| | | | Mut ASO | 16 | 33.81 ± 2.77 | | | Mut Veh vs. Mut ASO | 0.3008 | ns |
| | | Section 3 | WT Veh | 17 | 24.05 ± 1.82 | | | WT Veh vs. Mut Veh | <0.0001 | **** |
| | | | WT ASO | 15 | 28.17 ± 2.61 | | | WT Veh vs. WT ASO | >0.9999 | ns |
| | | | Mut Veh | 15 | 40.53 ± 2.49 | | | WT Veh vs. Mut ASO | >0.9999 | ns |
| | | | Mut ASO | 16 | 26.98 ± 2.17 | | | Mut Veh vs. Mut ASO | 0.0019 | ** |
| | | Section 4 | WT Veh | 17 | 17.86 ± 1.84 | | | WT Veh vs. Mut Veh | <0.0001 | **** |
| | | | WT ASO | 15 | 21.81 ± 2.21 | | | WT Veh vs. WT ASO | >0.9999 | ns |
| | | | Mut Veh | 15 | 37.25 ± 2.44 | | | WT Veh vs. Mut ASO | >0.9999 | ns |
| | | | Mut ASO | 16 | 21.32 ± 2.10 | | | Mut Veh vs. Mut ASO | 0.0002 | *** |
| | | Section 5 | WT Veh | 17 | 17.51 ± 1.40 | | | WT Veh vs. Mut Veh | <0.0001 | **** |
| | | | WT ASO | 15 | 20.75 ± 2.11 | | | WT Veh vs. WT ASO | >0.9999 | ns |
| | | | Mut Veh | 15 | 40.26 ± 1.55 | | | WT Veh vs. Mut ASO | >0.9999 | ns |
| | | | Mut ASO | 16 | 18.37 ± 2.42 | | | WT Veh vs. Mut ASO | <0.0001 | **** |
| | | Section 6 | WT Veh | 17 | 14.30 ± 1.20 | | | WT Veh vs. Mut Veh | <0.0001 | **** |
| | | | WT ASO | 15 | 22.35 ± 2.98 | | | WT Veh vs. WT ASO | 0.1303 | ns |
| | | | Mut Veh | 15 | 40.70 ± 2.30 | | | WT Veh vs. Mut ASO | 0.1197 | ns |
| | | | Mut ASO | 16 | 18.47 ± 1.20 | | | Mut Veh vs. Mut ASO | <0.0001 | **** |
| | | Section 7 | WT Veh | 17 | 17.45 ± 1.82 | | | WT Veh vs. Mut Veh | <0.0001 | **** |
| | | | WT ASO | 15 | 20.31 ± 1.92 | | | WT Veh vs. WT ASO | >0.9999 | ns |
| | | | Mut Veh | 15 | 38.36 ± 2.24 | | | WT Veh vs. Mut ASO | >0.9999 | ns |
| | | | Mut ASO | 16 | 19.58 ± 1.44 | | | Mut Veh vs. Mut ASO | <0.0001 | **** |
| | | Section 8 | WT Veh | 17 | 16.21 ± 1.87 | | | WT Veh vs. Mut Veh | <0.0001 | **** |
| | | | WT ASO | 15 | 15.82 ± 2.27 | | | WT Veh vs. WT ASO | >0.9999 | ns |
| | | | Mut Veh | 15 | 38.38 ± 2.22 | | | WT Veh vs. Mut ASO | >0.9999 | ns |
| | | | Mut ASO | 16 | 17.08 ± 1.20 | | | Mut Veh vs. Mut ASO | <0.0001 | **** |
| | | Section 9 | WT Veh | 17 | 16.01 ± 1.84 | | | WT Veh vs. Mut Veh | <0.0001 | **** |
| | | | WT ASO | 15 | 17.77 ± 2.53 | | | WT Veh vs. WT ASO | >0.9999 | ns |
| | | | Mut Veh | 15 | 40.96 ± 2.32 | | | WT Veh vs. Mut ASO | >0.9999 | ns |
| | | | Mut ASO | 16 | 17.38 ± 2.12 | | | Mut Veh vs. Mut ASO | <0.0001 | **** |
| | | Section 10 | WT Veh | 17 | 15.47 ± 1.48 | | | WT Veh vs. Mut Veh | <0.0001 | **** |
| | | | WT ASO | 15 | 17.84 ± 2.16 | | | WT Veh vs. WT ASO | >0.9999 | ns |
| | | | Mut Veh | 15 | 40.02 ± 2.54 | | | WT Veh vs. Mut ASO | >0.9999 | ns |
| | | | Mut ASO | 16 | 16.32 ± 1.31 | | | Mut Veh vs. Mut ASO | <0.0001 | **** |

| Extended Data Fig. 1a | Assay Performed | Parameter | Groups | N (animals) | Descriptive Statistics | Statistical Analysis | | Column statistics | |
|---|---|---|---|---|---|---|---|---|---|
| | | | | | | Two-Way ANOVA | | ferroni's multiple compar | |
| | | | | | Mean ± SEM | Statistical Test | Significance | P value | Significance |
| | MWM reversal learning test | Section 1 | WT | 16 | 36.74 ± 2.35 | | Interaction: F (9, 300) = 2.726, P=0.0045<br>Genotype: F (1, 300) = 80.52, P<0.0001<br>Section: F (9, 300) = 4.326, P<0.0001 | >0.9999 | ns |
| | | | Mut | 16 | 36.12 ± 3.14 | | | | |
| | | Section 2 | WT | 16 | 32.86 ± 3.62 | | | >0.9999 | ns |
| | | | Mut | 16 | 37.34 ± 2.08 | | | | |
| | | Section 3 | WT | 16 | 28.05 ± 3.37 | | | 0.7187 | ns |
| | | | Mut | 16 | 35.18 ± 2.87 | | | | |
| | | Section 4 | WT | 16 | 29.26 ± 2.65 | | | 0.0555 | ns |
| | | | Mut | 16 | 40.28 ± 2.99 | | | | |
| | | Section 5 | WT | 16 | 25.19 ± 2.36 | | | 0.1919 | ns |
| | | | Mut | 16 | 34.48 ± 2.01 | | | | |
| | | Section 6 | WT | 16 | 21.36 ± 2.65 | | | 0.0478 | * |
| | | | Mut | 16 | 32.57 ± 2.58 | | | | |
| | | Section 7 | WT | 16 | 18.86 ± 1.85 | | | <0.0001 | **** |
| | | | Mut | 16 | 40.06 ± 3.25 | | | | |
| | | Section 8 | WT | 16 | 19.01 ± 1.42 | | | 0.0003 | *** |
| | | | Mut | 16 | 35.81 ± 3.59 | | | | |
| | | Section 9 | WT | 16 | 17.38 ± 2.16 | | | 0.0002 | *** |
| | | | Mut | 16 | 34.63 ± 3.38 | | | | |
| | | Section 10 | WT | 16 | 17.70 ± 2.45 | | | 0.0038 | ** |
| | | | Mut | 16 | 31.88 ± 3.61 | | | | |

| Extended Data Fig. 1b | Assay Performed | Parameter | Groups | N (animals) | Descriptive Statistics | Statistical Analysis | |
|---|---|---|---|---|---|---|---|
| | | | | | | nn Whitney test - Two-ta | |
| | | | | | Mean ± SEM | P value | Significance |
| | MWM reversal probe trial | Quadrant 1 | WT | 16 | 33.40 ± 1.14 | p=0.0001 | *** |
| | | | Mut | 16 | 22.30 ± 2.05 | | |
| | | Quadrant 2 | WT | 16 | 23.06 ± 1.41 | p=0.2131 | ns |
| | | | Mut | 16 | 25.09 ± 1.32 | | |
| | | Quadrant 3 | WT | 16 | 20.97 ± 1.96 | p=0.0295 | * |
| | | | Mut | 16 | 27.54 ± 2.33 | | |
| | | Quadrant 4 | WT | 16 | 22.59 ± 1.55 | p=0.3089 | ns |
| | | | Mut | 16 | 25.02 ± 1.96 | | |

| Extended Data Fig. 1c | Assay Performed | Parameter | Groups | N (animals) | Descriptive Statistics | Statistical Analysis | |
|---|---|---|---|---|---|---|---|
| | | | | | | Mann Whitney test - Two-tailed | |
| | | | | | Mean ± SEM | P value | Significance |
| | Grooming | Grooming time | WT | 20 | 28.00 ± 2.29 | p=0.0039 | ** |
| | | | Mut | 21 | 42.79 ± 3.90 | | |

| Extended Data Fig. 1d | Assay Performed | Parameter | Groups | N (animals) | Descriptive Statistics | Statistical Analysis | |
|---|---|---|---|---|---|---|---|
| | | | | | | Mann Whitney test - Two-tailed | |
| | | | | | Mean ± SEM | P value | Significance |
| | Open field | Distance travelled | WT | 22 | 64.97 ± 2.81 | p=0.0675 | ns |
| | | | Mut | 20 | 55.45 ± 3.75 | | |

| Extended Data Fig. 1e | Assay Performed | Parameter | Groups | N (animals) | Descriptive Statistics | Statistical Analysis | |
|---|---|---|---|---|---|---|---|
| | | | | | | Mann Whitney test - Two-tailed | |
| | | | | | Mean ± SEM | P value | Significance |
| | Open field | Velocity | WT | 22 | 0.036 ± 0.001 | p=0.0523 | ns |
| | | | Mut | 20 | 0.031 ± 0.002 | | |

| Extended Data Fig. 1f | Assay Performed | Parameter | Groups | N (animals) | Descriptive Statistics | Statistical Analysis | |
|---|---|---|---|---|---|---|---|
| | | | | | | Mann Whitney test - Two-tailed | |
| | | | | | Mean ± SEM | P value | Significance |
| | Open field | Anxiety index | WT | 22 | 9.09 ± 0.89 | p=0.0159 | * |
| | | | Mut | 20 | 14.04 ± 1.48 | | |

| Extended Data Fig. 1g | Assay Performed | Parameter | Groups | N (animals) | Descriptive Statistics | Statistical Analysis | |
|---|---|---|---|---|---|---|---|
| | | | | | | Mann Whitney test - Two-tailed | |
| | | | | | Mean ± SEM | P value | Significance |
| | EPM | Anxiety index | WT | 16 | 57.67 ± 3.36 | p=0.0022 | ** |
| | | | Mut | 16 | 76.30 ± 4.77 | | |

| Extended Data Fig. 2b | Assay Performed | Parameter | Groups | N (animals) | Descriptive Statistics | Statistical Analysis | |
|---|---|---|---|---|---|---|---|
| | | | | | | Mann Whitney test - Two-tailed | |
| | | | | | Mean ± SEM | P value | Significance |
| | Western-PFC-GluA1 | Juvenile | WT | 7 | 1.00 ± 0.04 | 0.048 | * |
| | | | Mut | 5 | 0.80 ± 0.06 | | |
| | | Adult | WT | 7 | 1.00 ± 0.05 | 0.0012 | ** |
| | | | Mut | 7 | 0.65 ± 0.05 | | |

| Extended Data Fig. 2c | Assay Performed | Parameter | Groups | N (animals) | Descriptive Statistics | Statistical Analysis | |
|---|---|---|---|---|---|---|---|
| | | | | | | Mann Whitney test - Two-tailed | |
| | | | | | Mean ± SEM | P value | Significance |
| | Western-PFC-GluA2 | Juvenile | WT | 7 | 1.00 ± 0.05 | 0.1061 | ns |
| | | | Mut | 5 | 0.86 ± 0.07 | | |
| | | Adult | WT | 7 | 1.00 ± 0.03 | 0.0728 | ns |
| | | | Mut | 7 | 0.86 ± 0.043 | | |

| Extended Data Fig. 2d | Assay Performed | Parameter | Groups | N (animals) | Descriptive Statistics | Statistical Analysis | |
|---|---|---|---|---|---|---|---|
| | | | | | | Mann Whitney test - Two-tailed | |
| | | | | | Mean ± SEM | P value | Significance |
| | Western-PFC-GluA3 | Juvenile | WT | 7 | 1.00 ± 0.05 | 0.7551 | ns |
| | | | Mut | 5 | 1.05 ± 0.18 | | |
| | | Adult | WT | 7 | 1.00 ± 0.06 | 0.8048 | ns |
| | | | Mut | 7 | 1.03 ± 0.06 | | |

| Extended Data Fig. 2e | Assay Performed | Parameter | Groups | N (animals) | Descriptive Statistics | Statistical Analysis | |
|---|---|---|---|---|---|---|---|
| | | | | | | Mann Whitney test - Two-tailed | |
| | | | | | Mean ± SEM | P value | Significance |
| | Western-PFC-GluN1 | Juvenile | WT | 7 | 1.00 ± 0.16 | 0.4318 | ns |
| | | | Mut | 5 | 0.84 ± 0.20 | | |
| | | Adult | WT | 7 | 1.00 ± 0.08 | 0.2593 | ns |
| | | | Mut | 7 | 1.18 ± 0.09 | | |

| Extended Data Fig. 2g | Assay Performed | Parameter | Groups | N (animals) | Descriptive Statistics | Statistical Analysis | |
|---|---|---|---|---|---|---|---|
| | | | | | | Mann Whitney test - Two-tailed | |
| | | | | | Mean ± SEM | P value | Significance |
| | Western-SSC-GluA1 | Juvenile | WT | 7 | 1.00 ± 0.02 | 0.0012 | ** |
| | | | Mut | 6 | 0.78 ± 0.03 | | |
| | | Adult | WT | 7 | 1.00 ± 0.07 | 0.053 | ns |
| | | | Mut | 7 | 0.71 ± 0.08 | | |

| Extended Data Fig. 2h | Assay Performed | Parameter | Groups | N (animals) | Descriptive Statistics | Statistical Analysis | |
|---|---|---|---|---|---|---|---|
| | | | | | | Mann Whitney test - Two-tailed | |
| | | | | | Mean ± SEM | P value | Significance |
| | Western-SSC-GluA2 | Juvenile | WT | 7 | 1.00 ± 0.02 | 0.7308 | ns |
| | | | Mut | 6 | 0.94 ± 0.06 | | |
| | | Adult | WT | 7 | 1.00 ± 0.02 | 0.053 | ns |
| | | | Mut | 7 | 0.87 ± 0.04 | | |

| Extended Data Fig. 2i | Assay Performed | Parameter | Groups | N (animals) | Descriptive Statistics | Statistical Analysis | |
|---|---|---|---|---|---|---|---|
| | | | | | | Mann Whitney test - Two-tailed | |
| | | | | | Mean ± SEM | P value | Significance |
| | Western-SSC-GluA3 | Juvenile | WT | 7 | 1.00 ± 0.05 | 0.366 | ns |
| | | | Mut | 6 | 1.09 ± 0.06 | | |
| | | Adult | WT | 7 | 1.00 ± 0.08 | 0.0973 | ns |
| | | | Mut | 7 | 0.83 ± 0.03 | | |

| Extended Data Fig. 2j | Assay Performed | Parameter | Groups | N (animals) | Descriptive Statistics | Statistical Analysis | |
|---|---|---|---|---|---|---|---|
| | | | | | | Mann Whitney test - Two-tailed | |
| | | | | | Mean ± SEM | P value | Significance |
| | Western-SSC-GluN1 | Juvenile | WT | 7 | 1.00 ± 0.09 | 0.4452 | ns |
| | | | Mut | 6 | 1.11 ± 0.08 | | |
| | | Adult | WT | 7 | 1.00 ± 0.08 | 0.2086 | ns |
| | | | Mut | 7 | 0.85 ± 0.05 | | |

| Extended Data Fig. 2l | Assay Performed | Parameter | Groups | N (animals) | Descriptive Statistics | Statistical Analysis | |
|---|---|---|---|---|---|---|---|
| | | | | | | Mann Whitney test - Two-tailed | |
| | | | | | Mean ± SEM | P value | Significance |
| | Western-Thalamus-GluA1 | Juvenile | WT | 7 | 1.00 ± 0.17 | 0.1807 | ns |
| | | | Mut | 6 | 0.64 ± 0.10 | | |
| | | Adult | WT | 7 | 1.00 ± 0.10 | 0.014 | * |
| | | | Mut | 6 | 0.62 ± 0.07 | | |

| Extended Data Fig. 2m | Assay Performed | Parameter | Groups | N (animals) | Descriptive Statistics | Statistical Analysis | |
|---|---|---|---|---|---|---|---|
| | | | | | | Mann Whitney test - Two-tailed | |
| | | | | | Mean ± SEM | P value | Significance |
| | Western-Thalamus-GluA2 | Juvenile | WT | 10 | 1.00 ± 0.09 | 0.7802 | ns |
| | | | Mut | 9 | 0.98 ± 0.09 | | |
| | | Adult | WT | 9 | 1.00 ± 0.09 | 0.5457 | ns |
| | | | Mut | 9 | 0.88 ± 0.09 | | |

| Extended Data Fig. 2n | Assay Performed | Parameter | Groups | N (animals) | Descriptive Statistics | Statistical Analysis | |
|---|---|---|---|---|---|---|---|
| | | | | | | Mann Whitney test - Two-tailed | |
| | | | | | Mean ± SEM | P value | Significance |
| | Western-Thalamus-GluA3 | Juvenile | WT | 10 | 1.00 ± 0.11 | 0.7197 | ns |
| | | | Mut | 9 | 1.01 ± 0.11 | | |
| | | Adult | WT | 9 | 1.00 ± 0.07 | 0.9314 | ns |

| | | | | | | |
|---|---|---|---|---|---|---|
| | [illegible] | Mut | 9 | 0.99 ± 0.10 | [illegible] | [illegible] |

| Extended Data Fig. 2o | Assay Performed | Parameter | Groups | N (animals) | Descriptive Statistics | Statistical Analysis: Mann Whitney test - Two-tailed | |
|---|---|---|---|---|---|---|---|
| | | | | | Mean ± SEM | P value | Significance |
| | Western-Thalamus-GluN1 | Juvenile | WT | 10 | 1.00 ± 0.09 | >0.9999 | ns |
| | | | Mut | 9 | 1.02 ± 0.10 | | |
| | | Adult | WT | 9 | 1.00 ± 0.03 | 0.4894 | ns |
| | | | Mut | 9 | 1.02 ± 0.10 | | |

| Extended Data Fig. 2q | Assay Performed | Parameter | Groups | N (animals) | Descriptive Statistics | Statistical Analysis: Mann Whitney test - Two-tailed | |
|---|---|---|---|---|---|---|---|
| | | | | | Mean ± SEM | P value | Significance |
| | Western-Cerebellum-GluA1 | Juvenile | WT | 7 | 1.00 ± 0.06 | 0.0012 | ** |
| | | | Mut | 6 | 0.72 ± 0.080 | | |
| | | Adult | WT | 7 | 1.00 ± 0.05 | 0.0262 | * |
| | | | Mut | 7 | 0.65 ± 0.05 | | |

| Extended Data Fig. 2r | Assay Performed | Parameter | Groups | N (animals) | Descriptive Statistics | Statistical Analysis: Mann Whitney test - Two-tailed | |
|---|---|---|---|---|---|---|---|
| | | | | | Mean ± SEM | P value | Significance |
| | Western-Cerebellum-GluA2 | Juvenile | WT | 7 | 1.00 ± 0.05 | 0.366 | ns |
| | | | Mut | 6 | 1.13 ± 0.11 | | |
| | | Adult | WT | 7 | 1.00 ± 0.08 | 0.1649 | ns |
| | | | Mut | 7 | 1.20 ± 0.08 | | |

| Extended Data Fig. 2s | Assay Performed | Parameter | Groups | N (animals) | Descriptive Statistics | Statistical Analysis: Mann Whitney test - Two-tailed | |
|---|---|---|---|---|---|---|---|
| | | | | | Mean ± SEM | P value | Significance |
| | Western-Cerebellum-GluA3 | Juvenile | WT | 7 | 1.00 ± 0.09 | > 0.9999 | ns |
| | | | Mut | 6 | 0.96 ± 0.06 | | |
| | | Adult | WT | 7 | 1.00 ± 0.11 | 0.535 | ns |
| | | | Mut | 7 | 0.86 ± 0.09 | | |

| Extended Data Fig. 2t | Assay Performed | Parameter | Groups | N (animals) | Descriptive Statistics | Statistical Analysis: Mann Whitney test - Two-tailed | |
|---|---|---|---|---|---|---|---|
| | | | | | Mean ± SEM | P value | Significance |
| | Western-Cerebellum-GluN1 | Juvenile | WT | 7 | 1.00 ± 0.05 | 0.4452 | ns |
| | | | Mut | 6 | 0.91 ± 0.05 | | |
| | | Adult | WT | 7 | 1.00 ± 0.04 | 0.535 | ns |
| | | | Mut | 7 | 0.97 ± 0.07 | | |

| Extended Data Fig. 2u | Assay Performed | Parameter | Groups | N (animals) | Descriptive Statistics | Statistical Analysis: Mann Whitney test - Two-tailed | |
|---|---|---|---|---|---|---|---|
| | | | | | Mean ± SEM | P value | Significance |
| | qRT-PCR-Juvenile | *Gria1* | WT | 6 | 1.01 ± 0.07 | p=0.3680 | ns |
| | | | Mut | 6 | 0.92 ± 0.05 | | |
| | | *Gria2* | WT | 6 | 1.01 ± 0.05 | p=0.6190 | ns |
| | | | Mut | 6 | 0.98 ± 0.05 | | |
| | | *Gria3* | WT | 6 | 1.01 ± 0.06 | p=0.0714 | ns |
| | | | Mut | 6 | 0.84 ± 0.03 | | |
| | | *Gria4* | WT | 6 | 1.01 ± 0.05 | p=0.0649 | ns |
| | | | Mut | 6 | 1.12 ± 0.04 | | |
| | | *Grin1* | WT | 6 | 1.01 ± 0.07 | p>0.9999 | ns |
| | | | Mut | 6 | 1.00 ± 0.03 | | |
| | | *Grin2a* | WT | 6 | 1.01 ± 0.05 | p=0.2403 | ns |
| | | | Mut | 6 | 1.13 ± 0.07 | | |
| | | *Grin2b* | WT | 6 | 1.01 ± 0.06 | p=0.1797 | ns |
| | | | Mut | 6 | 1.19 ± 0.09 | | |
| | | *Cacng2* | WT | 6 | 1.01 ± 0.06 | p=0.0931 | ns |
| | | | Mut | 6 | 0.82 ± 0.06 | | |
| | | *Cacng8* | WT | 6 | 1.01 ± 0.07 | p=0.3615 | ns |
| | | | Mut | 6 | 1.03 ± 0.06 | | |

| Extended Data Fig. 2v | Assay Performed | Parameter | Groups | N (animals) | Descriptive Statistics | Statistical Analysis: Mann Whitney test - Two-tailed | |
|---|---|---|---|---|---|---|---|
| | | | | | Mean ± SEM | P value | Significance |
| | qRT-PCR-Adult | *Gria1* | WT | 6 | 1.00 ± 0.04 | p=0.2403 | ns |
| | | | Mut | 6 | 0.93 ± 0.02 | | |
| | | *Gria2* | WT | 6 | 1.01 ± 0.06 | p=0.8182 | ns |
| | | | Mut | 6 | 1.05 ± 0.04 | | |
| | | *Gria3* | WT | 6 | 1.01 ± 0.06 | p=0.3095 | ns |
| | | | Mut | 6 | 0.89 ± 0.03 | | |
| | | *Gria4* | WT | 6 | 1.02 ± 0.09 | p=0.1926 | ns |
| | | | Mut | 6 | 1.19 ± 0.03 | | |
| | | *Grin1* | WT | 6 | 1.02 ± 0.10 | p=0.9372 | ns |
| | | | Mut | 6 | 1.02 ± 0.07 | | |
| | | *Grin2a* | WT | 6 | 1.01 ± 0.05 | p=0.9372 | ns |
| | | | Mut | 6 | 0.99 ± 0.06 | | |
| | | *Grin2b* | WT | 6 | 1.01 ± 0.05 | p=0.7835 | ns |
| | | | Mut | 6 | 0.99 ± 0.02 | | |
| | | *Cacng2* | WT | 6 | 1.04 ± 0.13 | p=0.3095 | ns |
| | | | Mut | 6 | 0.87 ± 0.07 | | |
| | | *Cacng8* | WT | 6 | 1.03 ± 0.12 | p=0.2597 | ns |
| | | | Mut | 6 | 1.01 ± 0.08 | | |

| Extended Data Fig. 3c | Assay Performed | Parameter | Groups | N (animals) | Descriptive Statistics | Statistical Analysis | | Column statistics | | |
|---|---|---|---|---|---|---|---|---|---|---|
| | | | | | | Two-Way ANOVA | | Bonferroni's multiple comparisons | | |
| | | | | | Mean ± SEM | Statistical Test | Significance | Comparison | P value | Significance |
| | MRI-Whole brain | P21 | WT | 5 | 287.00 ± 9.03 | Interaction: F (1, 17) = 1.062, P=3173 | | p21-WT vs. p21-Mut | 0.536 | ns |
| | | | Mut | 6 | 270.56 ± 6.45 | Genotype: F (1, 17) = 12.41, P=0.0026 | | 2m-WT vs. 2m-Mut | 0.0349 | * |
| | | 2 mon | WT | 5 | 307.81 ± 6.14 | Age: F (1, 17) = 4.598, P=0.0468 | | p21-WT vs. 2m-WT | 0.2528 | ns |
| | | | Mut | 5 | 277.96 ± 2.97 | | | p21-Mut vs. 2m-Mut | >0.9999 | ns |

| Extended Data Fig. 3d | Assay Performed | Parameter | Groups | N (animals) | Descriptive Statistics | Statistical Analysis | | Column statistics | | |
|---|---|---|---|---|---|---|---|---|---|---|
| | | | | | | Two-Way ANOVA | | Bonferroni's multiple comparisons | | |
| | | | | | Mean ± SEM | Statistical Test | Significance | Comparison | P value | Significance |
| | MRI-Thalamus | P21 | WT | 5 | 16.23 ± 1.14 | Interaction: F (1, 17) = 0.7619, P=0.3949 | | p21-WT vs. p21-Mut | >0.9999 | ns |
| | | | Mut | 6 | 15.03 ± 0.73 | Genotype: F (1, 17) = 5.059, P=0.0380 | | 2m-WT vs. 2m-Mut | 0.2715 | ns |
| | | 2 mon | WT | 5 | 20.78 ± 0.94 | Age: F (1, 17) = 18.68, P= 0.0005 | | p21-WT vs. 2m-WT | 0.0134 | * |
| | | | Mut | 5 | 18.05 ± 0.66 | | | p21-Mut vs. 2m-Mut | 0.1395 | ns |

| Extended Data Fig. 3e | Assay Performed | Parameter | Groups | N (animals) | Descriptive Statistics | Statistical Analysis | | Column statistics | | |
|---|---|---|---|---|---|---|---|---|---|---|
| | | | | | | Two-Way ANOVA | | Bonferroni's multiple comparisons | | |
| | | | | | Mean ± SEM | Statistical Test | Significance | Comparison | P value | Significance |
| | MRI-Striatum | P21 | WT | 5 | 15.84 ± 0.63 | Interaction: F (1, 17) = 0.06335, P=0.8043 | | p21-WT vs. p21-Mut | >0.9999 | ** |
| | | | Mut | 6 | 12.48 ± 0.30 | Genotype: F (1, 17) = 0.3572, P=0.5580 | | p21-WT vs. p21-Mut | >0.9999 | **** |
| | | 2 mon | WT | 5 | 17.57 ± 1.02 | Age: F (1, 17) = 17.68, P=0.0006 | | p21-WT vs. 2m-WT | 0.0844 | ns |
| | | | Mut | 5 | 9.7 ± 0.44 | | | p21-Mut vs. 2m-Mut | 0.0301 | * |

| Extended Data Fig. 3f | Assay Performed | Parameter | Groups | N (animals) | Descriptive Statistics | Statistical Analysis | | Column statistics | | |
|---|---|---|---|---|---|---|---|---|---|---|
| | | | | | | Two-Way ANOVA | | Bonferroni's multiple comparisons | | |
| | | | | | Mean ± SEM | Statistical Test | Significance | Comparison | P value | Significance |
| | MRI-Ventricle | P21 | WT | 5 | 7.96 ± 0.96 | Interaction: F (1, 17) = 3.000, P=0.1014 | | p21-WT vs. p21-Mut | 0.7060 | ns |
| | | | Mut | 6 | 6.04 ± 0.19 | Genotype: F (1, 17) = 0.3000, P=0.5910 | | 2m-WT vs. 2m-Mut | >0.9999 | ns |
| | | 2 mon | WT | 5 | 11.43 ± 0.35 | Age: F (1, 17) = 34.15, P<0.0001 | | p21-WT vs. 2m-WT | 0.0670 | ns |
| | | | Mut | 5 | 12.43 ± 1.43 | | | p21-Mut vs. 2m-Mut | 0.0002 | *** |

| Extended Data Fig. 4a | Assay Performed | Parameter | Groups | N (animals) | Descriptive Statistics | Statistical Analysis | |
|---|---|---|---|---|---|---|---|
| | | | | | | Mann Whitney test- Two-tailed | |
| | | | | | Mean ± SEM | P value | Significance |
| | Branch legth- P14 | Total | WT | 5 | 3484 ± 63 | 0.0079 | ** |
| | | | Mut | 5 | 2798 ± 128 | | |
| | | Basal | WT | 5 | 1619 ± 50 | 0.0159 | * |
| | | | Mut | 5 | 1333 ± 54 | | |
| | | Apical | WT | 5 | 1864 ± 64 | 0.0317 | * |
| | | | Mut | 5 | 1465 ± 127 | | |

| Extended Data Fig. 4b | Assay Performed | Parameter | Groups | N (animals) | Descriptive Statistics | Statistical Analysis | |
|---|---|---|---|---|---|---|---|
| | | | | | | Mann Whitney test- Two-tailed | |
| | | | | | Mean ± SEM | P value | Significance |
| | Branch legth- P21 | Total | WT | 5 | 4585 ± 206 | 0.0159 | * |
| | | | Mut | 5 | 3528 ± 193 | | |
| | | Basal | WT | 5 | 2098 ± 78 | 0.0079 | ** |
| | | | Mut | 5 | 1449 ± 67 | | |
| | | Apical | WT | 5 | 2487 ± 168 | 0.2222 | ns |
| | | | Mut | 5 | 2078 ± 148 | | |

| Extended Data Fig. 4c | Assay Performed | Parameter | Groups | N (animals) | Descriptive Statistics | Statistical Analysis | |
|---|---|---|---|---|---|---|---|
| | | | | | | Mann Whitney test- Two-tailed | |
| | | | | | Mean ± SEM | P value | Significance |
| | Branch legth- 2 months | Total | WT | 5 | 5238 ± 276 | 0.0079 | ** |
| | | | Mut | 5 | 3525 ± 281 | | |
| | | Basal | WT | 5 | 2430 ± 144 | 0.0079 | ** |
| | | | Mut | 5 | 1345 ± 170 | | |
| | | Apical | WT | 5 | 2904 ± 152 | 0.0317 | * |
| | | | Mut | 5 | 2347 ± 143 | | |

| Extended Data Fig. 4d | Assay Performed | Parameter | Groups | N (animals) | Descriptive Statistics | Statistical Analysis | |
|---|---|---|---|---|---|---|---|
| | | | | | | Mann Whitney test- Two-tailed | |
| | | | | | Mean ± SEM | P value | Significance |
| | Relative Dendritic brench length | P14 | WT | 5 | 100.0 ± 1.8 | 0.0079 | ** |
| | | | Mut | 5 | 80.2 ± 3.7 | | |
| | | P21 | WT | 5 | 100.0 ± 4.6 | 0.0159 | * |
| | | | Mut | 5 | 76.8 ± 4.3 | | |
| | | Adult | WT | 5 | 100.0 ± 5.3 | 0.0079 | ** |
| | | | Mut | 5 | 67.4 ± 5.3 | | |

| Extended Data Fig. 4e | Assay Performed | Parameter | Groups | N (animals) | Descriptive Statistics | Statistical Analysis | |
|---|---|---|---|---|---|---|---|
| | | | | | | Mann Whitney test- Two-tailed | |
| | | | | | Mean ± SEM | P value | Significance |
| | Branch points- P14 | Total | WT | 5 | 40.44 ± 1.16 | 0.0079 | ** |
| | | | Mut | 5 | 34.88 ± 1.16 | | |
| | | Basal | WT | 5 | 18.12 ± 0.74 | 0.0635 | ns |
| | | | Mut | 5 | 16.12 ± 0.94 | | |
| | | Apical | WT | 5 | 22.32 ± 0.91 | 0.0317 | * |
| | | | Mut | 5 | 18.76 ± 1.24 | | |

| Extended Data Fig. 4f | Assay Performed | Parameter | Groups | N (animals) | Descriptive Statistics | Statistical Analysis | |
|---|---|---|---|---|---|---|---|
| | | | | | | Mann Whitney test- Two-tailed | |
| | | | | | Mean ± SEM | P value | Significance |
| | Branch points- P21 | Total | WT | 5 | 57.72 ± 1.99 | 0.9524 | ns |
| | | | Mut | 5 | 56.84 ± 3.89 | | |
| | | Basal | WT | 5 | 24.20 ± 1.06 | 0.0794 | ns |
| | | | Mut | 5 | 21.08 ± 1.15 | | |
| | | Apical | WT | 5 | 33.52 ± 1.99 | 0.5952 | ns |
| | | | Mut | 5 | 35.76 ± 3.31 | | |

| Extended Data Fig. 4g | Assay Performed | Parameter | Groups | N (animals) | Descriptive Statistics | Statistical Analysis | |
|---|---|---|---|---|---|---|---|
| | | | | | | Mann Whitney test- Two-tailed | |
| | | | | | Mean ± SEM | P value | Significance |
| | Branch points- 2 months | Total | WT | 5 | 61.05 ± 2.71 | 0.6905 | ns |
| | | | Mut | 5 | 59.49 ± 3.20 | | |
| | | Basal | WT | 5 | 25.68 ± 1.80 | 0.0952 | ns |
| | | | Mut | 5 | 20.69 ± 1.71 | | |
| | | Apical | WT | 5 | 38.86 ± 1.25 | 0.0307 | ns |
| | | | Mut | 5 | 38.34 ± 2.61 | | |

| Extended Data Fig. 4h | Assay Performed | Parameter | Groups | N (animals) | Descriptive Statistics | Statistical Analysis | |
|---|---|---|---|---|---|---|---|
| | | | | | | Mann Whitney test- Two-tailed | |
| | | | | | Mean ± SEM | P value | Significance |
| | Relative brench point | P14 | WT | 5 | 99.8 ± 2.8 | 0.079 | ** |
| | | | Mut | 5 | 86.4 ± 2.8 | | |
| | | P21 | WT | 5 | 100.0 ± 3.4 | 0.9524 | ns |
| | | | Mut | 5 | 98.4 ± 6.7 | | |
| | | Adult | WT | 5 | 100.2 ± 4.4 | 0.6905 | ns |
| | | | Mut | 5 | 97.4 ± 54 | | |

| Extended Data Fig. 5b | Assay Performed | Parameter | Groups | N (animals) | Descriptive Statistics | Statistical Analysis | | Column statistics | |
|---|---|---|---|---|---|---|---|---|---|
| | | | | | | Two-Way ANOVA | | Bonferroni's multiple comparisons | |
| | | | | | Mean ± SEM | Statistical Test | Significance | P value | Significance |
| | Sholl Analysis- P21 Basal dendrites | Distance- 0 | WT | 3 | 0.00 ± 0.00 | Distance x Genotype: F (26, 104) = 0.7612, P=0.7851<br>Distance: F (26, 104) = 199.4, P <0.0001<br>Genotype: F (1, 4) = 5.548, P=0.0780 | | >0.9999 | ns |
| | | | Mut | 3 | 0.00 ± 0.00 | | | | |
| | | Distance- 10 | WT | 3 | 10.67 ± 0.44 | | | >0.9999 | ns |
| | | | Mut | 3 | 10.42 ± 1.01 | | | | |
| | | Distance- 20 | WT | 3 | 13.41 ± 0.14 | | | >0.9999 | ns |
| | | | Mut | 3 | 13.33 ± 1.20 | | | | |
| | | Distance- 30 | WT | 3 | 18.58 ± 0.13 | | | >0.9999 | ns |
| | | | Mut | 3 | 17.15 ± 1.35 | | | | |
| | | Distance- 40 | WT | 3 | 21.62 ± 0.71 | | | >0.9999 | ns |
| | | | Mut | 3 | 20.17 ± 0.72 | | | | |
| | | Distance- 50 | WT | 3 | 23.37 ± 0.51 | | | 0.7154 | ns |
| | | | Mut | 3 | 20.52 ± 0.26 | | | | |
| | | Distance- 60 | WT | 3 | 23.18 ± 1.17 | | | >0.9999 | ns |
| | | | Mut | 3 | 20.68 ± 0.36 | | | | |
| | | Distance- 70 | WT | 3 | 22.67 ± 1.02 | | | 0.587 | ns |
| | | | Mut | 3 | 19.72 ± 1.37 | | | | |
| | | Distance- 80 | WT | 3 | 20.53 ± 0.67 | | | >0.9999 | ns |
| | | | Mut | 3 | 17.95 ± 1.48 | | | | |
| | | Distance- 90 | WT | 3 | 18.20 ± 1.73 | | | >0.9999 | ns |
| | | | Mut | 3 | 16.57 ± 1.16 | | | | |
| | | Distance- 100 | WT | 3 | 15.84 ± 1.52 | | | >0.9999 | ns |
| | | | Mut | 3 | 13.55 ± 1.21 | | | | |
| | | Distance- 110 | WT | 3 | 13.60 ± 2.29 | | | >0.9999 | ns |
| | | | Mut | 3 | 11.95 ± 1.18 | | | | |
| | | Distance- 120 | WT | 3 | 12.13 ± 1.95 | | | >0.9999 | ns |
| | | | Mut | 3 | 9.50 ± 0.61 | | | | |
| | | Distance- 130 | WT | 3 | 9.99 ± 1.58 | | | >0.9999 | ns |
| | | | Mut | 3 | 7.57 ± 0.47 | | | | |
| | | Distance- 140 | WT | 3 | 7.93 ± 1.58 | | | >0.9999 | ns |
| | | | Mut | 3 | 5.87 ± 0.77 | | | | |
| | | Distance- 150 | WT | 3 | 5.99 ± 1.45 | | | >0.9999 | ns |
| | | | Mut | 3 | 430 ± 0.67 | | | | |
| | | Distance- 160 | WT | 3 | 4.66 ± 1.19 | | | >0.9999 | ns |
| | | | Mut | 3 | 3.05 ± 0.63 | | | | |
| | | Distance- 170 | WT | 3 | 3.48 ± 0.66 | | | >0.9999 | ns |
| | | | Mut | 3 | 2.02 ± 0.42 | | | | |
| | | Distance- 180 | WT | 3 | 2.63 ± 0.45 | | | >0.9999 | ns |
| | | | Mut | 3 | 1.45 ± 0.43 | | | | |
| | | Distance- 190 | WT | 3 | 1.78 ± 0.22 | | | >0.9999 | ns |
| | | | Mut | 3 | 0.93 ± 0.29 | | | | |
| | | Distance- 200 | WT | 3 | 1.02 ± 0.19 | | | >0.9999 | ns |
| | | | Mut | 3 | 0.48 ± 0.26 | | | | |
| | | Distance- 210 | WT | 3 | 0.63 ± 0.13 | | | >0.9999 | ns |
| | | | Mut | 3 | 0.28 ± 0.17 | | | | |
| | | Distance- 220 | WT | 3 | 0.57 ± 0.19 | | | >0.9999 | ns |
| | | | Mut | 3 | 0.22 ± 0.12 | | | | |
| | | Distance- 230 | WT | 3 | 0.29 ± 0.11 | | | >0.9999 | ns |
| | | | Mut | 3 | 0.80 ± 0.08 | | | | |
| | | Distance- 240 | WT | 3 | 0.06 ± 0.06 | | | >0.9999 | ns |
| | | | Mut | 3 | 0.80 ± 0.08 | | | | |
| | | Distance- 250 | WT | 3 | 0.00 ± 0.00 | | | >0.9999 | ns |
| | | | Mut | 3 | 0.80 ± 0.08 | | | | |
| | | Distance- 260 | WT | 3 | 0.00 ± 0.00 | | | >0.9999 | ns |
| | | | Mut | 3 | 0.00 ± 0.00 | | | | |

| Assay Performed | Parameter | Groups | N (animals) | Descriptive Statistics | Statistical Analysis | | Column statistics | |
|---|---|---|---|---|---|---|---|---|
| | | | | | Two-Way ANOVA | | Bonferroni's multiple comparisons | |
| | | | | Mean ± SEM | Statistical Test | Significance | P value | Significance |
| | Distance- 0 | WT | 3 | 0.00 ± 0.00 | | | >0.9999 | ns |
| | | Mut | 3 | 0.00 ± 0.00 | | | | |
| | Distance- 10 | WT | 3 | 1.07 ± 0.07 | | | >0.9999 | ns |
| | | Mut | 3 | 1.00 ± 0.00 | | | | |
| | Distance- 20 | WT | 3 | 2.79 ± 0.30 | | | >0.9999 | ns |
| | | Mut | 3 | 2.48 ± 0.56 | | | | |
| | Distance- 30 | WT | 3 | 5.26 ± 0.33 | | | >0.9999 | ns |
| | | Mut | 3 | 4.23 ± 0.54 | | | | |
| | Distance- 40 | WT | 3 | 8.50 ± 0.76 | | | >0.9999 | ns |
| | | Mut | 3 | 5.75 ± 1.10 | | | | |
| | Distance- 50 | WT | 3 | 9.89 ± 0.29 | | | >0.9999 | ns |
| | | Mut | 3 | 7.57 ± 1.09 | | | | |
| | Distance- 60 | WT | 3 | 11.53 ± 0.39 | | | 0.5878 | ns |
| | | Mut | 3 | 8.53 ± 1.01 | | | | |
| | Distance- 70 | WT | 3 | 12.83 ± 0.26 | | | 0.2774 | ns |
| | | Mut | 3 | 9.57 ± 1.30 | | | | |
| | Distance- 80 | WT | 3 | 12.29 ± 0.20 | | | >0.9999 | ns |
| | | Mut | 3 | 10.00 ± 1.44 | | | | |
| | Distance- 90 | WT | 3 | 12.51 ± 0.40 | | | 0.8923 | ns |
| | | Mut | 3 | 9.67 ± 1.23 | | | | |
| | Distance- 100 | WT | 3 | 12.72 ± 1.02 | | | >0.9999 | ns |
| | | Mut | 3 | 10.72 ± 0.62 | | | | |
| | Distance- 110 | WT | 3 | 11.63 ± 1.40 | | | >0.9999 | ns |
| | | Mut | 3 | 10.07 ± 0.48 | | | | |
| | Distance- 120 | WT | 3 | 10.82 ± 0.77 | | | >0.9999 | ns |
| | | Mut | 3 | 8.73 ± 0.64 | | | | |
| | Distance- 130 | WT | 3 | 9.56 ± 0.99 | | | >0.9999 | ns |
| | | Mut | 3 | 8.33 ± 0.33 | | | | |
| | Distance- 140 | WT | 3 | 8.78 ± 0.68 | | | >0.9999 | ns |
| | | Mut | 3 | 6.98 ± 0.69 | | | | |
| | Distance- 150 | WT | 3 | 7.69 ± 1.03 | | | >0.9999 | ns |
| | | Mut | 3 | 7.07 ± 0.98 | | | | |
| | Distance- 160 | WT | 3 | 6.77 ± 0.89 | | | >0.9999 | ns |
| | | Mut | 3 | 6.22 ± 0.52 | | | | |
| | Distance- 170 | WT | 3 | 6.40 ± 1.07 | | | >0.9999 | ns |
| | | Mut | 3 | 6.08 ± 1.08 | | | | |
| | Distance- 180 | WT | 3 | 5.17 ± 1.11 | | | >0.9999 | ns |
| | | Mut | 3 | 5.22 ± 1.04 | | | | |
| | Distance- 190 | WT | 3 | 4.63 ± 1.14 | | | >0.9999 | ns |
| | | Mut | 3 | 5.15 ± 1.08 | | | | |
| | Distance- 200 | WT | 3 | 4.09 ± 0.85 | | | >0.9999 | ns |
| | | Mut | 3 | 3.72 ± 0.62 | | | | |
| | Distance- 210 | WT | 3 | 3.39 ± 0.84 | | | >0.9999 | ns |
| | | Mut | 3 | 3.77 ± 0.50 | | | | |
| | Distance- 220 | WT | 3 | 2.94 ± 0.58 | | | >0.9999 | ns |
| | | Mut | 3 | 3.42 ± 0.30 | | | | |
| | Distance- 230 | WT | 3 | 2.19 ± 0.18 | | | >0.9999 | ns |
| | | Mut | 3 | 2.68 ± 0.46 | | | | |
| | Distance- 240 | WT | 3 | 2.12 ± 0.06 | | | >0.9999 | ns |
| | | Mut | 3 | 2.17 ± 0.44 | | | | |
| | Distance- 250 | WT | 3 | 1.91 ± 0.31 | | | >0.9999 | ns |
| | | Mut | 3 | 1.95 ± 0.39 | | | | |
| | Distance- 260 | WT | 3 | 1.42 ± 0.13 | | | >0.9999 | ns |
| | | Mut | 3 | 1.82 ± 0.49 | | | | |
| | Distance- 270 | WT | 3 | 1.30 ± 0.07 | | | >0.9999 | ns |
| | | Mut | 3 | 1.62 ± 0.39 | | | | |
| | Distance- 280 | WT | 3 | 1.38 ± 0.21 | | | >0.9999 | ns |
| | | Mut | 3 | 1.42 ± 0.30 | | | | |
| | Distance- 290 | WT | 3 | 1.47 ± 0.19 | | | >0.9999 | ns |

| | | | | | | | |
|---|---|---|---|---|---|---|---|
| Sholl Analysis- p21 Apical dendrites | Distance- 290 | Mut | 3 | 1.68 ± 0.56 | Distance x Genotype: F (90, 360) = 0.7489, P=0.9504<br>Distance: F (90, 360) = 29.65, P <0.0001<br>Genotype: F (1, 4) = 1.299, P=0.3180 | >0.9999 | ns |
| | Distance- 300 | WT | 3 | 1.30 ± 0.15 | | >0.9999 | ns |
| | | Mut | 3 | 1.35 ± 0.24 | | | |
| | Distance- 310 | WT | 3 | 1.42 ± 0.13 | | >0.9999 | ns |
| | | Mut | 3 | 1.50 ± 0.29 | | | |
| | Distance- 320 | WT | 3 | 1.42 ± 0.09 | | >0.9999 | ns |
| | | Mut | 3 | 1.35 ± 0.24 | | | |
| | Distance- 330 | WT | 3 | 1.30 ± 0.07 | | >0.9999 | ns |
| | | Mut | 3 | 1.28 ± 0.17 | | | |
| | Distance- 340 | WT | 3 | 1.19 ± 0.12 | | >0.9999 | ns |
| | | Mut | 3 | 1.48 ± 0.16 | | | |
| | Distance- 350 | WT | 3 | 1.19 ± 0.12 | | >0.9999 | ns |
| | | Mut | 3 | 1.47 ± 0.24 | | | |
| | Distance- 360 | WT | 3 | 1.26 ± 0.18 | | >0.9999 | ns |
| | | Mut | 3 | 1.55 ± 0.16 | | | |
| | Distance- 370 | WT | 3 | 1.31 ± 0.17 | | >0.9999 | ns |
| | | Mut | 3 | 1.62 ± 0.18 | | | |
| | Distance- 380 | WT | 3 | 1.37 ± 0.13 | | >0.9999 | ns |
| | | Mut | 3 | 1.62 ± 0.18 | | | |
| | Distance- 390 | WT | 3 | 1.48 ± 0.16 | | >0.9999 | ns |
| | | Mut | 3 | 1.83 ± 0.17 | | | |
| | Distance- 400 | WT | 3 | 1.82 ± 0.10 | | >0.9999 | ns |
| | | Mut | 3 | 1.83 ± 0.20 | | | |
| | Distance- 410 | WT | 3 | 1.66 ± 0.09 | | >0.9999 | ns |
| | | Mut | 3 | 1.70 ± 0.15 | | | |
| | Distance-420 | WT | 3 | 1.60 ± 0.14 | | >0.9999 | ns |
| | | Mut | 3 | 1.70 ± 0.15 | | | |
| | Distance- 430 | WT | 3 | 1.60 ± 0.14 | | >0.9999 | ns |
| | | Mut | 3 | 1.70 ± 0.15 | | | |
| | Distance- 440 | WT | 3 | 1.71 ± 0.11 | | >0.9999 | ns |
| | | Mut | 3 | 1.77 ± 0.15 | | | |
| | Distance- 450 | WT | 3 | 1.77 ± 0.05 | | >0.9999 | ns |
| | | Mut | 3 | 1.92 ± 0.14 | | | |
| | Distance- 460 | WT | 3 | 1.77 ± 0.05 | | >0.9999 | ns |
| | | Mut | 3 | 2.25 ± 0.30 | | | |
| | Distance- 470 | WT | 3 | 1.75 ± 0.08 | | >0.9999 | ns |
| | | Mut | 3 | 2.12 ± 0.34 | | | |
| | Distance- 480 | WT | 3 | 1.92 ± 0.17 | | >0.9999 | ns |
| | | Mut | 3 | 2.12 ± 0.44 | | | |
| | Distance- 490 | WT | 3 | 2.09 ± 0.26 | | >0.9999 | ns |
| | | Mut | 3 | 2.20 ± 0.42 | | | |
| | Distance- 500 | WT | 3 | 2.16 ± 0.20 | | >0.9999 | ns |
| | | Mut | 3 | 2.27 ± 0.48 | | | |
| | Distance- 510 | WT | 3 | 2.48 ± 0.08 | | >0.9999 | ns |
| | | Mut | 3 | 2.35 ± 0.46 | | | |
| | Distance- 520 | WT | 3 | 2.59 ± 0.05 | | >0.9999 | ns |
| | | Mut | 3 | 2.35 ± 0.46 | | | |
| | Distance- 530 | WT | 3 | 2.87 ± 0.13 | | >0.9999 | ns |
| | | Mut | 3 | 2.52 ± 0.48 | | | |
| | Distance- 540 | WT | 3 | 2.98 ± 0.19 | | >0.9999 | ns |
| | | Mut | 3 | 2.70 ± 0.59 | | | |
| | Distance- 550 | WT | 3 | 3.14 ± 0.28 | | >0.9999 | ns |
| | | Mut | 3 | 3.00 ± 0.69 | | | |
| | Distance- 560 | WT | 3 | 3.53 ± 0.50 | | >0.9999 | ns |
| | | Mut | 3 | 3.12 ± 0.77 | | | |
| | Distance- 570 | WT | 3 | 3.98 ± 0.71 | | >0.9999 | ns |
| | | Mut | 3 | 3.60 ± 0.83 | | | |
| | Distance- 580 | WT | 3 | 4.03 ± 0.73 | | >0.9999 | ns |
| | | Mut | 3 | 3.18 ± 0.84 | | | |
| | Distance- 590 | WT | 3 | 4.32 ± 0.76 | | >0.9999 | ns |
| | | Mut | 3 | 3.10 ± 0.86 | | | |
| | Distance- 600 | WT | 3 | 4.43 ± 0.83 | | >0.9999 | ns |
| | | Mut | 3 | 3.45 ± 1.10 | | | |
| | Distance- 610 | WT | 3 | 4.39 ± 0.74 | | >0.9999 | ns |
| | | Mut | 3 | 3.63 ±1.29 | | | |
| | Distance- 620 | WT | 3 | 4.87 ± 0.70 | | >0.9999 | ns |
| | | Mut | 3 | 3.72 ± 1.04 | | | |
| | Distance- 630 | WT | 3 | 4.43 ± 0.54 | | >0.9999 | ns |
| | | Mut | 3 | 3.85 ± 1.18 | | | |
| | Distance- 640 | WT | 3 | 4.48 ± 0.53 | | >0.9999 | ns |
| | | Mut | 3 | 3.85 ± 1.08 | | | |
| | Distance- 650 | WT | 3 | 4.30 ± 0.55 | | >0.9999 | ns |
| | | Mut | 3 | 4.22 ± 0.89 | | | |
| | Distance- 660 | WT | 3 | 5.00 ± 0.54 | | >0.9999 | ns |
| | | Mut | 3 | 4.72 ± 0.75 | | | |
| | Distance- 670 | WT | 3 | 4.63 ± 0.58 | | >0.9999 | ns |
| | | Mut | 3 | 5.17 ± 0.38 | | | |
| | Distance- 680 | WT | 3 | 4.98 ± 0.92 | | >0.9999 | ns |
| | | Mut | 3 | 4.85 ± 0.64 | | | |
| | Distance- 690 | WT | 3 | 4.73 ± 0.81 | | >0.9999 | ns |
| | | Mut | 3 | 5.03 ± 0.73 | | | |
| | Distance- 700 | WT | 3 | 5.81 ± 1.37 | | >0.9999 | ns |
| | | Mut | 3 | 5.58 ± 0.99 | | | |
| | Distance- 710 | WT | 3 | 6.33 ± 2.09 | | >0.9999 | ns |
| | | Mut | 3 | 5.65 ± 1.39 | | | |
| | Distance- 720 | WT | 3 | 6.80 ± 1.80 | | >0.9999 | ns |
| | | Mut | 3 | 5.97 ± 2.09 | | | |
| | Distance- 730 | WT | 3 | 6.27 ± 1.78 | | >0.9999 | ns |
| | | Mut | 3 | 5.43 ± 2.19 | | | |
| | Distance- 740 | WT | 3 | 5.82 ± 1.65 | | >0.9999 | ns |
| | | Mut | 3 | 4.88 ± 1.42 | | | |
| | Distance- 750 | WT | 3 | 5.58 ± 1.69 | | >0.9999 | ns |
| | | Mut | 3 | 4.68 ± 1.35 | | | |
| | Distance- 760 | WT | 3 | 4.80 ± 1.26 | | >0.9999 | ns |
| | | Mut | 3 | 3.67 ± 1.11 | | | |
| | Distance- 770 | WT | 3 | 4.34 ± 1.54 | | >0.9999 | ns |
| | | Mut | 3 | 3.85 ± 1.33 | | | |
| | Distance- 780 | WT | 3 | 3.11 ± 1.43 | | >0.9999 | ns |
| | | Mut | 3 | 3.42 ± 1.10 | | | |
| | Distance- 790 | WT | 3 | 2.62 ± 1.44 | | >0.9999 | ns |
| | | Mut | 3 | 3.85 ± 1.18 | | | |
| | Distance- 800 | WT | 3 | 1.53 ± 0.98 | | >0.9999 | ns |
| | | Mut | 3 | 3.08 ± 1.76 | | | |
| | Distance- 810 | WT | 3 | 1.15 ± 0.77 | | >0.9999 | ns |
| | | Mut | 3 | 2.15 ± 0.83 | | | |
| | Distance- 820 | WT | 3 | 0.82 ± 0.48 | | >0.9999 | ns |
| | | Mut | 3 | 1.72 ± 0.75 | | | |
| | Distance- 830 | WT | 3 | 0.39 ± 0.39 | | >0.9999 | ns |
| | | Mut | 3 | 1.58 ± 0.87 | | | |
| | Distance- 840 | WT | 3 | 0.28 ± 0.28 | | >0.9999 | ns |
| | | Mut | 3 | 0.95 ± 0.83 | | | |
| | Distance- 850 | WT | 3 | 0.22 ± 0.22 | | >0.9999 | ns |
| | | Mut | 3 | 0.53 ± 0.53 | | | |
| | Distance- 860 | WT | 3 | 0.22 ± 0.22 | | >0.9999 | ns |
| | | Mut | 3 | 0.07 ± 0.07 | | | |
| | Distance- 870 | WT | 3 | 0.11 ± 0.11 | | >0.9999 | ns |
| | | Mut | 3 | 0.00 ± 0.00 | | | |
| | Distance- 880 | WT | 3 | 0.06 ± 0.06 | | >0.9999 | ns |
| | | Mut | 3 | 0.00 ± 0.00 | | | |

| | | | | | | | | | |
|---|---|---|---|---|---|---|---|---|---|
| | | Distance- 890 | WT | 3 | 0.00 ± 0.00 | | | >0.9999 | ns |
| | | | Mut | 3 | 0.00 ± 0.00 | | | | |
| | | Distance- 900 | WT | 3 | 0.00 ± 0.00 | | | >0.9999 | ns |
| | | | Mut | 3 | 0.00 ± 0.00 | | | | |

| Extended Data Fig. 5d | Assay Performed | Parameter | Groups | N (animals) | Descriptive Statistics | Statistical Analysis | | Column statistics | |
|---|---|---|---|---|---|---|---|---|---|
| | | | | | | Two-Way ANOVA | | Bonferroni's multiple comparisons | |
| | | | | | Mean ± SEM | Statistical Test | Significance | P value | Significance |
| | | Distance- 0 | WT | 3 | 0.00 ± 0.00 | | | >0.9999 | ns |
| | | | Mut | 3 | 0.00 ± 0.00 | | | | |
| | | Distance- 10 | WT | 3 | 9.46 ± 0.34 | | | >0.9999 | ns |
| | | | Mut | 3 | 9.90 ± 0.46 | | | | |
| | | Distance- 20 | WT | 3 | 14.25 ± 0.66 | | | >0.9999 | ns |
| | | | Mut | 3 | 14.29 ± 0.51 | | | | |
| | | Distance- 30 | WT | 3 | 19.46 ± 1.11 | | | >0.9999 | ns |
| | | | Mut | 3 | 19.01 ± 1.26 | | | | |
| | | Distance- 40 | WT | 3 | 20.71 ± 0.77 | | | >0.9999 | ns |
| | | | Mut | 3 | 21.31 ± 0.04 | | | | |
| | | Distance- 50 | WT | 3 | 22.13 ± 1.34 | | | >0.9999 | ns |
| | | | Mut | 3 | 22.59 ± 0.62 | | | | |
| | | Distance- 60 | WT | 3 | 23.04 ± 0.51 | | | >0.9999 | ns |
| | | | Mut | 3 | 22.00 ± 1.40 | | | | |
| | | Distance- 70 | WT | 3 | 21.46 ± 1.12 | | | >0.9999 | ns |
| | | | Mut | 3 | 20.85 ± 1.83 | | | | |
| | | Distance- 80 | WT | 3 | 18.83 ± 1.17 | | | >0.9999 | ns |
| | | | Mut | 3 | 19.09 ± 2.31 | | | | |
| | | Distance- 90 | WT | 3 | 17.54 ± 1.49 | | | >0.9999 | ns |
| | | | Mut | 3 | 16.80 ± 2.33 | | | | |
| | | Distance- 100 | WT | 3 | 15.54 ± 1.72 | | | >0.9999 | ns |
| | | | Mut | 3 | 14.75 ± 2.81 | | | | |
| | | Distance- 110 | WT | 3 | 14.54 ± 2.27 | | | >0.9999 | ns |
| | | | Mut | 3 | 12.70 ± 3.13 | | | | |
| | | Distance- 120 | WT | 3 | 12.50 ± 1.89 | | | >0.9999 | ns |
| | | | Mut | 3 | 10.09 ± 2.89 | | | | |
| | | Distance- 130 | WT | 3 | 10.04 ± 2.13 | | | >0.9999 | ns |
| | | | Mut | 3 | 8.21 ± 2.54 | | | | |
| | | Distance- 140 | WT | 3 | 7.21 ± 1.69 | | | >0.9999 | ns |
| | | | Mut | 3 | 6.15 ± 2.22 | | | | |
| | | Distance- 150 | WT | 3 | 5.50 ± 1.28 | | | >0.9999 | ns |
| | | | Mut | 3 | 4.44 ± 2.07 | | | | |
| | | Distance- 160 | WT | 3 | 3.75 ± 1.01 | | | >0.9999 | ns |
| | | | Mut | 3 | 3.48 ± 1.72 | | | | |
| | Sholl Analysis- 2 months Basal dendrites | Distance- 170 | WT | 3 | 2.42 ± 0.44 | Distance x Genotype: F (34, 136) = 0.2278, P>0.9999<br>Distance: F (34, 136) = 129.2, P <0.0001<br>Genotype: F (1, 4) = 0.1159, P=0.7506 | | >0.9999 | ns |
| | | | Mut | 3 | 2.64 ± 1.31 | | | | |
| | | Distance- 180 | WT | 3 | 1.71 ± 0.43 | | | >0.9999 | ns |
| | | | Mut | 3 | 2.05 ± 1.02 | | | | |
| | | Distance- 190 | WT | 3 | 0.75 ± 0.14 | | | >0.9999 | ns |
| | | | Mut | 3 | 1.44 ± 0.76 | | | | |
| | | Distance- 200 | WT | 3 | 0.63 ± 0.19 | | | >0.9999 | ns |
| | | | Mut | 3 | 0.69 ± 0.46 | | | | |
| | | Distance- 210 | WT | 3 | 0.54 ± 0.36 | | | >0.9999 | ns |
| | | | Mut | 3 | 0.37 ± 0.26 | | | | |
| | | Distance- 220 | WT | 3 | 0.33 ± 0.33 | | | >0.9999 | ns |
| | | | Mut | 3 | 0.24 ± 0.24 | | | | |
| | | Distance- 230 | WT | 3 | 0.33 ± 0.33 | | | >0.9999 | ns |
| | | | Mut | 3 | 0.19 ± 0.19 | | | | |
| | | Distance- 240 | WT | 3 | 0.33 ± 0.33 | | | >0.9999 | ns |
| | | | Mut | 3 | 0.14 ± 0.14 | | | | |
| | | Distance- 250 | WT | 3 | 0.25 ± 0.25 | | | >0.9999 | ns |
| | | | Mut | 3 | 0.05 ± 0.05 | | | | |
| | | Distance- 260 | WT | 3 | 0.17 ± 0.17 | | | >0.9999 | ns |
| | | | Mut | 3 | 0.05 ± 0.05 | | | | |
| | | Distance- 270 | WT | 3 | 0.25 ± 0.25 | | | >0.9999 | ns |
| | | | Mut | 3 | 0.05 ± 0.05 | | | | |
| | | Distance- 280 | WT | 3 | 0.08 ± 0.08 | | | >0.9999 | ns |
| | | | Mut | 3 | 0.05 ± 0.05 | | | | |
| | | Distance- 290 | WT | 3 | 0.08 ± 0.08 | | | >0.9999 | ns |
| | | | Mut | 3 | 0.00 ± 0.00 | | | | |
| | | Distance- 300 | WT | 3 | 0.08 ± 0.08 | | | >0.9999 | ns |
| | | | Mut | 3 | 0.00 ± 0.00 | | | | |
| | | Distance- 310 | WT | 3 | 0.08 ± 0.08 | | | >0.9999 | ns |
| | | | Mut | 3 | 0.00 ± 0.00 | | | | |
| | | Distance- 320 | WT | 3 | 0.08 ± 0.08 | | | >0.9999 | ns |
| | | | Mut | 3 | 0.00 ± 0.00 | | | | |
| | | Distance- 330 | WT | 3 | 0.08 ± 0.08 | | | >0.9999 | ns |
| | | | Mut | 3 | 0.00 ± 0.00 | | | | |
| | | Distance- 340 | WT | 3 | 0.00 ± 0.00 | | | >0.9999 | ns |
| | | | Mut | 3 | 0.00 ± 0.00 | | | | |

| Assay Performed | Parameter | Groups | N (animals) | Descriptive Statistics | Statistical Analysis | | Column statistics | |
|---|---|---|---|---|---|---|---|---|
| | | | | | Two-Way ANOVA | | Bonferroni's multiple comparisons | |
| | | | | Mean ± SEM | Statistical Test | Significance | P value | Significance |
| | Distance- 0 | WT | 3 | 0.00 ± 0.00 | | | >0.9999 | ns |
| | | Mut | 3 | 0.00 ± 0.00 | | | | |
| | Distance- 10 | WT | 3 | 1.21 ± 0.04 | | | >0.9999 | ns |
| | | Mut | 3 | 1.15 ± 0.08 | | | | |
| | Distance- 20 | WT | 3 | 2.83 ± 0.74 | | | >0.9999 | ns |
| | | Mut | 3 | 2.13 ± 0.47 | | | | |
| | Distance- 30 | WT | 3 | 4.83 ± 0.33 | | | >0.9999 | ns |
| | | Mut | 3 | 4.74 ± 0.53 | | | | |
| | Distance- 40 | WT | 3 | 7.17 ± 0.51 | | | >0.9999 | ns |
| | | Mut | 3 | 6.16 ± 0.76 | | | | |
| | Distance- 50 | WT | 3 | 8.29 ± 0.36 | | | >0.9999 | ns |
| | | Mut | 3 | 8.19 ± 1.28 | | | | |
| | Distance- 60 | WT | 3 | 9.08 ± 0.42 | | | >0.9999 | ns |
| | | Mut | 3 | 8.77 ± 1.03 | | | | |
| | Distance- 70 | WT | 3 | 10.71 ± 1.05 | | | >0.9999 | ns |
| | | Mut | 3 | 10.93 ± 1.57 | | | | |
| | Distance- 80 | WT | 3 | 11.38 ± 0.92 | | | >0.9999 | ns |
| | | Mut | 3 | 10.09 ± 1.97 | | | | |
| | Distance- 90 | WT | 3 | 10.71 ± 0.29 | | | >0.9999 | ns |
| | | Mut | 3 | 10.60 ± 2.66 | | | | |
| | Distance- 100 | WT | 3 | 10.50 ± 0.95 | | | >0.9999 | ns |
| | | Mut | 3 | 9.45 ± 2.33 | | | | |
| | Distance- 110 | WT | 3 | 10.38 ± 0.99 | | | >0.9999 | ns |
| | | Mut | 3 | 9.34 ± 2.36 | | | | |
| | Distance- 120 | WT | 3 | 9.63 ± 0.73 | | | >0.9999 | ns |
| | | Mut | 3 | 9.62 ± 2.37 | | | | |
| | Distance- 130 | WT | 3 | 8.71 ± 0.32 | | | >0.9999 | ns |
| | | Mut | 3 | 9.42 ± 2.57 | | | | |
| | Distance- 140 | WT | 3 | 7.96 ± 0.29 | | | >0.9999 | ns |
| | | Mut | 3 | 8.66 ± 2.63 | | | | |
| | Distance- 150 | WT | 3 | 6.67 ± 0.08 | | | >0.9999 | ns |
| | | Mut | 3 | 8.01 ± 2.81 | | | | |
| | Distance- 160 | WT | 3 | 5.79 ± 0.23 | | | >0.9999 | ns |
| | | Mut | 3 | 6.86 ± 2.26 | | | | |
| | Distance- 170 | WT | 3 | 5.29 ± 0.36 | | | >0.9999 | ns |
| | | Mut | 3 | 6.16 ± 1.98 | | | | |
| | Distance- 180 | WT | 3 | 4.42 ± 0.58 | | | >0.9999 | ns |

| | | | | | | | |
|---|---|---|---|---|---|---|---|
| | Distance- 180 | Mut | 3 | 5.82 ± 1.71 | | >0.9999 | ns |
| | Distance- 190 | WT | 3 | 4.04 ± 0.83 | | >0.9999 | ns |
| | | Mut | 3 | 5.03 ± 1.43 | | | |
| | Distance- 200 | WT | 3 | 3.58 ± 0.88 | | >0.9999 | ns |
| | | Mut | 3 | 4.82 ± 1.31 | | | |
| | Distance- 210 | WT | 3 | 3.00 ± 0.50 | | >0.9999 | ns |
| | | Mut | 3 | 3.96 ± 0.99 | | | |
| | Distance- 220 | WT | 3 | 3.04 ± 0.76 | | >0.9999 | ns |
| | | Mut | 3 | 3.41 ± 0.81 | | | |
| | Distance- 230 | WT | 3 | 2.54 ± 0.68 | | >0.9999 | ns |
| | | Mut | 3 | 2.70 ± 0.46 | | | |
| | Distance- 240 | WT | 3 | 2.67 ± 0.79 | | >0.9999 | ns |
| | | Mut | 3 | 2.37 ± 0.26 | | | |
| | Distance- 250 | WT | 3 | 2.67 ± 0.30 | | >0.9999 | ns |
| | | Mut | 3 | 1.97 ± 0.30 | | | |
| | Distance- 260 | WT | 3 | 2.63 ± 0.26 | | >0.9999 | ns |
| | | Mut | 3 | 1.83 ± 0.16 | | | |
| | Distance- 270 | WT | 3 | 2.08 ± 0.36 | | >0.9999 | ns |
| | | Mut | 3 | 1.75 ± 0.20 | | | |
| | Distance- 280 | WT | 3 | 1.83 ± 0.08 | | >0.9999 | ns |
| | | Mut | 3 | 1.60 ± 0.06 | | | |
| | Distance- 290 | WT | 3 | 1.96 ± 0.27 | | >0.9999 | ns |
| | | Mut | 3 | 1.57 ± 0.18 | | | |
| | Distance- 300 | WT | 3 | 2.13 ± 0.19 | | >0.9999 | ns |
| | | Mut | 3 | 1.39 ± 0.20 | | | |
| | Distance- 310 | WT | 3 | 2.13 ± 0.19 | | >0.9999 | ns |
| | | Mut | 3 | 1.49 ± 0.25 | | | |
| | Distance- 320 | WT | 3 | 2.29 ± 0.36 | | >0.9999 | ns |
| | | Mut | 3 | 1.44 ± 0.22 | | | |
| | Distance- 330 | WT | 3 | 2.00 ± 0.25 | | >0.9999 | ns |
| | | Mut | 3 | 1.60 ± 0.28 | | | |
| | Distance- 340 | WT | 3 | 1.83 ± 0.22 | | >0.9999 | ns |
| | | Mut | 3 | 1.66 ± 0.26 | | | |
| | Distance- 350 | WT | 3 | 1.83 ± 0.22 | | >0.9999 | ns |
| | | Mut | 3 | 1.47 ± 0.11 | | | |
| | Distance- 360 | WT | 3 | 1.79 ± 0.04 | | >0.9999 | ns |
| | | Mut | 3 | 1.56 ± 0.23 | | | |
| | Distance- 370 | WT | 3 | 1.58 ± 0.08 | | >0.9999 | ns |
| | | Mut | 3 | 1.56 ± 0.23 | | | |
| | Distance- 380 | WT | 3 | 1.58 ± 0.08 | | >0.9999 | ns |
| | | Mut | 3 | 1.82 ± 0.29 | | | |
| | Distance- 390 | WT | 3 | 1.71 ± 0.11 | | >0.9999 | ns |
| | | Mut | 3 | 1.65 ± 0.38 | | | |
| | Distance- 400 | WT | 3 | 1.92 ± 0.08 | | >0.9999 | ns |
| | | Mut | 3 | 1.56 ± 0.43 | | | |
| | Distance- 410 | WT | 3 | 1.83 ± 0.08 | | >0.9999 | ns |
| | | Mut | 3 | 1.61 ± 0.41 | | | |
| | Distance-420 | WT | 3 | 1.92 ± 0.17 | | >0.9999 | ns |
| | | Mut | 3 | 1.76 ± 0.42 | | | |
| | Distance- 430 | WT | 3 | 2.00 ± 0.14 | | >0.9999 | ns |
| | | Mut | 3 | 1.87 ± 0.47 | | | |
| | Distance- 440 | WT | 3 | 2.00 ± 0.14 | | >0.9999 | ns |
| | | Mut | 3 | 1.94 ± 0.54 | | | |
| | Distance- 450 | WT | 3 | 2.25 ± 0.14 | | >0.9999 | ns |
| | | Mut | 3 | 2.07 ± 0.47 | | | |
| **Sholl Analysis- 2 months Apical dendrites** | Distance- 460 | WT | 3 | 2.50 ± 0.29 | Distance x Genotype: F (93, 372) = 0.3025, P>0.9999<br>Distance: F (93, 372) = 16.95, P <0.0001<br>Genotype: F (1, 4) = 0.3896, P=0.5663 | >0.9999 | ns |
| | | Mut | 3 | 2.19 ± 0.52 | | | |
| | Distance- 470 | WT | 3 | 2.71 ± 0.32 | | >0.9999 | ns |
| | | Mut | 3 | 2.20 ± 0.60 | | | |
| | Distance- 480 | WT | 3 | 2.75 ± 0.25 | | >0.9999 | ns |
| | | Mut | 3 | 2.18 ± 0.71 | | | |
| | Distance- 490 | WT | 3 | 2.96 ± 0.32 | | >0.9999 | ns |
| | | Mut | 3 | 2.31 ± 0.85 | | | |
| | Distance- 500 | WT | 3 | 3.25 ± 0.38 | | >0.9999 | ns |
| | | Mut | 3 | 2.42 ± 0.89 | | | |
| | Distance- 510 | WT | 3 | 3.29 ± 0.40 | | >0.9999 | ns |
| | | Mut | 3 | 2.57 ± 0.91 | | | |
| | Distance- 520 | WT | 3 | 3.38 ± 0.40 | | >0.9999 | ns |
| | | Mut | 3 | 2.57 ± 0.91 | | | |
| | Distance- 530 | WT | 3 | 3.46 ± 0.42 | | >0.9999 | ns |
| | | Mut | 3 | 2.69 ± 0.96 | | | |
| | Distance- 540 | WT | 3 | 3.63 ± 0.50 | | >0.9999 | ns |
| | | Mut | 3 | 2.80 ± 1.00 | | | |
| | Distance- 550 | WT | 3 | 4.25 ± 0.52 | | >0.9999 | ns |
| | | Mut | 3 | 2.87 ± 1.07 | | | |
| | Distance- 560 | WT | 3 | 5.46 ± 0.90 | | >0.9999 | ns |
| | | Mut | 3 | 3.15 ± 1.14 | | | |
| | Distance- 570 | WT | 3 | 5.38 ± 0.99 | | >0.9999 | ns |
| | | Mut | 3 | 3.54 ± 1.33 | | | |
| | Distance- 580 | WT | 3 | 5.38 ± 0.82 | | >0.9999 | ns |
| | | Mut | 3 | 3.78 ± 1.31 | | | |
| | Distance- 590 | WT | 3 | 5.21 ± 0.47 | | >0.9999 | ns |
| | | Mut | 3 | 4.14 ± 1.53 | | | |
| | Distance- 600 | WT | 3 | 5.00 ± 0.25 | | >0.9999 | ns |
| | | Mut | 3 | 4.30 ± 1.56 | | | |
| | Distance- 610 | WT | 3 | 4.71 ± 0.25 | | >0.9999 | ns |
| | | Mut | 3 | 4.81 ± 1.40 | | | |
| | Distance- 620 | WT | 3 | 4.33 ± 0.36 | | >0.9999 | ns |
| | | Mut | 3 | 4.78 ± 1.05 | | | |
| | Distance- 630 | WT | 3 | 4.42 ± 0.08 | | >0.9999 | ns |
| | | Mut | 3 | 4.90 ± 0.67 | | | |
| | Distance- 640 | WT | 3 | 4.63 ± 0.19 | | >0.9999 | ns |
| | | Mut | 3 | 4.61 ± 0.41 | | | |
| | Distance- 650 | WT | 3 | 4.96 ± 0.36 | | >0.9999 | ns |
| | | Mut | 3 | 4.66 ± 0.12 | | | |
| | Distance- 660 | WT | 3 | 5.42 ± 0.96 | | >0.9999 | ns |
| | | Mut | 3 | 5.53 ± 0.36 | | | |
| | Distance- 670 | WT | 3 | 4.92 ± 1.09 | | >0.9999 | ns |
| | | Mut | 3 | 5.22 ± 0.33 | | | |
| | Distance- 680 | WT | 3 | 5.71 ± 1.61 | | >0.9999 | ns |
| | | Mut | 3 | 5.09 ± 0.42 | | | |
| | Distance- 690 | WT | 3 | 6.38 ± 1.81 | | >0.9999 | ns |
| | | Mut | 3 | 4.95 ± 1.08 | | | |
| | Distance- 700 | WT | 3 | 6.29 ± 1.52 | | >0.9999 | ns |
| | | Mut | 3 | 4.09 ± 1.43 | | | |
| | Distance- 710 | WT | 3 | 4.96 ± 1.43 | | >0.9999 | ns |
| | | Mut | 3 | 4.27 ± 0.91 | | | |
| | Distance- 720 | WT | 3 | 4.58 ± 1.23 | | >0.9999 | ns |
| | | Mut | 3 | 4.31 ± 1.16 | | | |
| | Distance- 730 | WT | 3 | 4.38 ± 1.41 | | >0.9999 | ns |
| | | Mut | 3 | 4.11 ± 1.64 | | | |
| | Distance- 740 | WT | 3 | 4.50 ± 1.51 | | >0.9999 | ns |
| | | Mut | 3 | 4.09 ± 1.92 | | | |
| | Distance- 750 | WT | 3 | 3.33 ± 0.98 | | >0.9999 | ns |
| | | Mut | 3 | 4.18 ± 1.99 | | | |
| | Distance- 760 | WT | 3 | 3.13 ± 1.27 | | >0.9999 | ns |
| | | Mut | 3 | 3.70 ± 1.69 | | | |
| | Distance- 770 | WT | 3 | 2.08 ± 0.79 | | >0.9999 | ns |
| | | Mut | 3 | 2.63 ± 1.68 | | | |

| | | | | | | |
|---|---|---|---|---|---|---|
| | Distance- 780 | WT | 3 | 1.33 ± 0.55 | >0.9999 | ns |
| | | Mut | 3 | 2.18 ± 1.58 | | |
| | Distance- 790 | WT | 3 | 1.13 ± 0.59 | >0.9999 | ns |
| | | Mut | 3 | 1.15 ± 0.82 | | |
| | Distance- 800 | WT | 3 | 1.71 ± 1.19 | >0.9999 | ns |
| | | Mut | 3 | 1.68 ± 1.10 | | |
| | Distance- 810 | WT | 3 | 1.00 ± 0.66 | >0.9999 | ns |
| | | Mut | 3 | 0.99 ± 0.66 | | |
| | Distance- 820 | WT | 3 | 0.83 ± 0.60 | >0.9999 | ns |
| | | Mut | 3 | 0.88 ± 0.81 | | |
| | Distance- 830 | WT | 3 | 0.54 ± 0.48 | >0.9999 | ns |
| | | Mut | 3 | 1.00 ± 1.00 | | |
| | Distance- 840 | WT | 3 | 0.50 ± 0.50 | >0.9999 | ns |
| | | Mut | 3 | 1.17 ± 1.17 | | |
| | Distance- 850 | WT | 3 | 0.42 ± 0.42 | >0.9999 | ns |
| | | Mut | 3 | 1.42 ± 1.42 | | |
| | Distance- 860 | WT | 3 | 0.67 ± 0.67 | >0.9999 | ns |
| | | Mut | 3 | 0.75 ± 0.75 | | |
| | Distance- 870 | WT | 3 | 0.58 ± 0.58 | >0.9999 | ns |
| | | Mut | 3 | 0.42 ± 0.42 | | |
| | Distance- 880 | WT | 3 | 0.67 ± 0.67 | >0.9999 | ns |
| | | Mut | 3 | 0.00 ± 0.00 | | |
| | Distance- 890 | WT | 3 | 0.58 ± 0.58 | >0.9999 | ns |
| | | Mut | 3 | 0.00 ± 0.00 | | |
| | Distance- 900 | WT | 3 | 0.42 ± 0.42 | >0.9999 | ns |
| | | Mut | 3 | 0.00 ± 0.00 | | |
| | Distance- 910 | WT | 3 | 0.33 ± 0.33 | >0.9999 | ns |
| | | Mut | 3 | 0.00 ± 0.00 | | |
| | Distance- 920 | WT | 3 | 0.33 ± 0.33 | >0.9999 | ns |
| | | Mut | 3 | 0.00 ± 0.00 | | |
| | Distance- 930 | WT | 3 | 0.00 ± 0.00 | >0.9999 | ns |
| | | Mut | 3 | 0.00 ± 0.00 | | |

| **Extended Data Fig. 5e** | **Assay Performed** | **Parameter** | **Groups** | **N (animals)** | **Descriptive Statistics** | **Statistical Analysis** | |
|---|---|---|---|---|---|---|---|
| | | | | | | **Mann Whitney test- Two-tailed** | |
| | | | | | **Mean ± SEM** | **P value** | **Significance** |
| | **Branch length-P21** | Total | WT | 5 | 8422 ± 303 | 0.2000 | ns |
| | | | Mut | 5 | 7382 ± 469 | | |
| | | Basal | WT | 5 | 3099 ± 150 | 0.1000 | ns |
| | | | Mut | 5 | 2660 ± 55 | | |
| | | Apical | WT | 5 | 5323 ± 417 | 0.7000 | ns |
| | | | Mut | 5 | 4723 ± 487 | | |

| **Extended Data Fig. 5f** | **Assay Performed** | **Parameter** | **Groups** | **N (animals)** | **Descriptive Statistics** | **Statistical Analysis** | |
|---|---|---|---|---|---|---|---|
| | | | | | | **Mann Whitney test- Two-tailed** | |
| | | | | | **Mean ± SEM** | **P value** | **Significance** |
| | **Branch length-2 months** | Total | WT | 5 | 8062 ± 279 | >0.9999 | ns |
| | | | Mut | 5 | 7747 ± 835 | | |
| | | Basal | WT | 5 | 2956 ± 141 | >0.9999 | ns |
| | | | Mut | 5 | 2887 ± 297 | | |
| | | Apical | WT | 5 | 5106 ± 144 | 0.7000 | ns |
| | | | Mut | 5 | 4862 ± 540 | | |

| **Extended Data Fig. 5g** | **Assay Performed** | **Parameter** | **Groups** | **N (animals)** | **Descriptive Statistics** | **Statistical Analysis** | |
|---|---|---|---|---|---|---|---|
| | | | | | | **Mann Whitney test- Two-tailed** | |
| | | | | | **Mean ± SEM** | **P value** | **Significance** |
| | **Branch points-P21** | Total | WT | 5 | 67.91 ± 3.26 | 0.2000 | ns |
| | | | Mut | 5 | 56.57 ± 6.14 | | |
| | | Basal | WT | 5 | 23.79 ± 1.16 | 0.1000 | ns |
| | | | Mut | 5 | 19.28 ± 0.40 | | |
| | | Apical | WT | 5 | 44.01 ± 4.03 | 0.4000 | ns |
| | | | Mut | 5 | 37.22 ± 5.68 | | |

| **Extended Data Fig. 5h** | **Assay Performed** | **Parameter** | **Groups** | **N (animals)** | **Descriptive Statistics** | **Statistical Analysis** | |
|---|---|---|---|---|---|---|---|
| | | | | | | **Mann Whitney test- Two-tailed** | |
| | | | | | **Mean ± SEM** | **P value** | **Significance** |
| | **Branch points-2months** | Total | WT | 5 | 68.96 ± 3.68 | 0.4000 | ns |
| | | | Mut | 5 | 62.84 ± 3.60 | | |
| | | Basal | WT | 5 | 25.88 ± 1.82 | 0.4000 | ns |
| | | | Mut | 5 | 23.08 ± 1.69 | | |
| | | Apical | WT | 5 | 43.00 ± 2.18 | >0.9999 | ns |
| | | | Mut | 5 | 39.76 ± 4.29 | | |

| Extended Data Fig. 6b | Assay Performed | Parameter | Groups | N (slices) | Descriptive Statistics | Statistical Analysis | |
|---|---|---|---|---|---|---|---|
| | | | | | | Mann Whitney test- Two-tailed | |
| | | | | | Mean ± SEM | P value | Significance |
| | AMPAR-mEPSC | Holding current | WT | 20 | 30.10 ± 4.64 | 0.8875 | ns |
| | | | Mut | 16 | 27.99 ± 3.65 | | |

| Extended Data Fig. 6c | Assay Performed | Parameter | Groups | N (slices) | Descriptive Statistics | Statistical Analysis | |
|---|---|---|---|---|---|---|---|
| | | | | | | Mann Whitney test- Two-tailed | |
| | | | | | Mean ± SEM | P value | Significance |
| | AMPAR-mEPSC | Membrane Resistance | WT | 20 | 1407.2 ± 104.49 | 0.2234 | ns |
| | | | Mut | 16 | 1534.5 ± 79.07 | | |

| Extended Data Fig. 6d | Assay Performed | Parameter | Groups | N (slices) | Descriptive Statistics | Statistical Analysis | |
|---|---|---|---|---|---|---|---|
| | | | | | | Mann Whitney test- Two-tailed | |
| | | | | | Mean ± SEM | P value | Significance |
| | AMPAR-mEPSC | Capacitance | WT | 20 | 49.02 ± 2.00 | 0.7176 | ns |
| | | | Mut | 16 | 46.68 ± 2.75 | | |

| Extended Data Fig. 6e | Assay Performed | Parameter | Groups | N (slices) | Descriptive Statistics | Statistical Analysis | |
|---|---|---|---|---|---|---|---|
| | | | | | | Mann Whitney test- Two-tailed | |
| | | | | | Mean ± SEM | P value | Significance |
| | AMPAR-mEPSC | Charge | WT | 21 | 74.00 ± 4.35 | 0.0015 | ** |
| | | | Mut | 17 | 101.80 ± 5.35 | | |

| Extended Data Fig. 6f | Assay Performed | Parameter | Groups | N (slices) | Descriptive Statistics | Statistical Analysis | |
|---|---|---|---|---|---|---|---|
| | | | | | | Mann Whitney test- Two-tailed | |
| | | | | | Mean ± SEM | P value | Significance |
| | AMPAR-mEPSC | Amplitude | WT | 20 | 12.90 ± 0.41 | 0.3359 | ns |
| | | | Mut | 16 | 13.70 ± 0.51 | | |

| Extended Data Fig. 6g | Assay Performed | Parameter | Groups | N (slices) | Descriptive Statistics | Statistical Analysis | |
|---|---|---|---|---|---|---|---|
| | | | | | | Mann Whitney test- Two-tailed | |
| | | | | | Mean ± SEM | P value | Significance |
| | AMPAR-mEPSC | Decay tau | WT | 20 | 5.45 ± 0.19 | p<0.0001 | **** |
| | | | Mut | 16 | 7.01 ± 0.21 | | |

| Extended Data Fig. 6h | Assay Performed | Parameter | Groups | N (slices) | Descriptive Statistics | Statistical Analysis | |
|---|---|---|---|---|---|---|---|
| | | | | | | Mann Whitney test- Two-tailed | |
| | | | | | Mean ± SEM | P value | Significance |
| | AMPAR-mEPSC | Frequency | WT | 20 | 0.10 ± 0.01 | 0.0239 | * |
| | | | Mut | 16 | 0.16 ± 0.02 | | |

| Extended Data Fig. 6i | Assay Performed | Parameter | Groups | N (slices) | Descriptive Statistics | Statistical Analysis | |
|---|---|---|---|---|---|---|---|
| | | | | | | Mann Whitney test- Two-tailed | |
| | | | | | Mean ± SEM | P value | Significance |
| | AMPAR/NMDAR ratio | Ratio | WT | 17 | 0.91 ± 0.09 | 0.3771 | ns |
| | | | Mut | 14 | 1.09 ± 0.14 | | |

| Extended Data Fig. 7e | Assay Performed | Parameter | Groups | N (animals) | Descriptive Statistics | Statistical Analysis | | Column statistics | |
|---|---|---|---|---|---|---|---|---|---|
| | | | | | | Two-Way ANOVA | | Bonferroni's multiple comparisons | |
| | | | | | Mean ± SEM | Statistical Test | Significance | P value | Significance |
| | DAPI staining-Hippocampus thickness | P7 | WT | 4 | 1037 ± 14.41 | Interaction: F (3, 32) = 7.430, P value = 0.0007<br>Genotype: F (1, 32) = 44.30, P value = <0.0001<br>Age: F (3, 32) = 2.948, P value = 0.0475 | | 0.6948 | ns |
| | | | Mut | 4 | 978 ± 26.43 | | | | |
| | | P14 | WT | 5 | 1064 ± 24.90 | | | 0.8059 | ns |
| | | | Mut | 5 | 1019 ± 14.98 | | | | |
| | | P21 | WT | 7 | 1122 ± 36.07 | | | 0.0002 | *** |
| | | | Mut | 5 | 921 ± 30.40 | | | | |
| | | Adult | WT | 5 | 1255 ± 42.94 | | | <0.0001 | **** |
| | | | Mut | 5 | 942 ± 38.92 | | | | |

| Extended Data Fig. 7f | Assay Performed | Parameter | Groups | N (animals) | Descriptive Statistics | Statistical Analysis | | Column statistics | |
|---|---|---|---|---|---|---|---|---|---|
| | | | | | | Two-Way ANOVA | | Bonferroni's multiple comparisons | |
| | | | | | Mean ± SEM | Statistical Test | Significance | P value | Significance |
| | TUNEL staining-CA1 | P7 | WT | 4 | 25.19 ± 4.80 | Interaction: F (3, 31) = 2.092, P=0.1215<br>Genotype: F (1, 31) = 46.24, P<0.0001<br>Age: F (3, 31) = 0.9250, P=0.4403 | | 0.4293 | ns |
| | | | Mut | 4 | 46.83 ± 3.77 | | | | |
| | | P14 | WT | 5 | 17.52 ± 4.08 | | | 0.0155 | * |
| | | | Mut | 5 | 53.91 ± 11.04 | | | | |
| | | P21 | WT | 7 | 12.87 ± 2.52 | | | <0.0001 | **** |
| | | | Mut | 5 | 75.44 ± 15.76 | | | | |
| | | Adult | WT | 4 | 9.63 ± 2.26 | | | 0.0066 | ** |
| | | | Mut | 5 | 52.28 ± 10.11 | | | | |

| Extended Data Fig. 7g | Assay Performed | Parameter | Groups | N (animals) | Descriptive Statistics | Statistical Analysis | | Column statistics | |
|---|---|---|---|---|---|---|---|---|---|
| | | | | | | Two-Way ANOVA | | Bonferroni's multiple comparisons | |
| | | | | | Mean ± SEM | Statistical Test | Significance | P value | Significance |
| | TUNEL staining-CA3 | P7 | WT | 4 | 33.39 ± 5.85 | Interaction: F (3, 32) = 3.134, P=0.390<br>Genotype: F (1, 32) = 14.69, P=0.0006<br>Age: F (3, 32) = 1.991, P=0.1352 | | 0.9996 | ns |
| | | | Mut | 4 | 36.93 ± 6.35 | | | | |
| | | P14 | WT | 5 | 19.85 ± 2.92 | | | 0.9053 | ns |
| | | | Mut | 5 | 33.85 ± 5.540 | | | | |
| | | P21 | WT | 7 | 16.74 ± 2.83 | | | 0.0068 | *** |
| | | | Mut | 5 | 93.28 ± 28.87 | | | | |
| | | Adult | WT | 5 | 23.64 ± 10.86 | | | 0.0387 | * |
| | | | Mut | 5 | 82.38 ± 22.28 | | | | |

| Extended Data Fig. 7h | Assay Performed | Parameter | Groups | N (animals) | Descriptive Statistics | Statistical Analysis | | Column statistics | |
|---|---|---|---|---|---|---|---|---|---|
| | | | | | | Two-Way ANOVA | | Bonferroni's multiple comparisons | |
| | | | | | Mean ± SEM | Statistical Test | Significance | P value | Significance |
| | TUNEL staining-DG | P7 | WT | 4 | 40.42 ± 11.79 | Interaction: F (3, 32) = 0.6236, P=0.6050<br>Genotype: F (1, 32) = 7.455, P=0.0102<br>Age: F (3, 32) = 0.3150, P=0.8144 | | >0.9999 | ns |
| | | | Mut | 4 | 48.25 ± 15.84 | | | | |
| | | P14 | WT | 5 | 35.73 ± 4.62 | | | 0.939 | ns |
| | | | Mut | 5 | 60.05 ± 10.31 | | | | |
| | | P21 | WT | 7 | 34.17 ± 10.14 | | | 0.0692 | ns |
| | | | Mut | 5 | 80.85 ± 22.75 | | | | |
| | | Adult | WT | 5 | 35.59 ± 11.01 | | | 0.4708 | ns |
| | | | Mut | 5 | 67.88 ± 19.93 | | | | |

| Extended Data Fig. 8c | Assay Performed | Parameter | Groups | N (animals) | Descriptive Statistics | Statistical Analysis | | Column statistics | | |
|---|---|---|---|---|---|---|---|---|---|---|
| | | | | | | Two-Way ANOVA | | Bonferroni's multiple comparisons | | |
| | | | | | Mean ± SEM | Statistical Test | Significance | Comparison | P value | Significance |
| | TUNEL staining-CA1 | WT | Veh | 3 | 22.97 ± 4.84 | Interaction: F (1, 10) = 3.700, P=0.0833<br>Treatment: F (1, 10) = 21.26, P= 0.0010<br>Genotype: F (1, 10) = 85.45, P<0.0001 | | WT Veh vs. Mut Veh | <0.0001 | **** |
| | | | JNJ | 3 | 6.41 ± 3.52 | | | WT Veh vs. WT JNJ | 0.6354 | ns |
| | | Mut | Veh | 4 | 91.79 ± 5.64 | | | WT Veh vs. Mut JNJ | 0.0500 | * |
| | | | JNJ | 4 | 51.52 ± 7.67 | | | Mut Veh vs. Mut JNJ | 0.0033 | ** |

| Extended Data Fig. 8d | Assay Performed | Parameter | Groups | N (animals) | Descriptive Statistics | Statistical Analysis | | Column statistics | | |
|---|---|---|---|---|---|---|---|---|---|---|
| | | | | | | Two-Way ANOVA | | Bonferroni's multiple comparisons | | |
| | | | | | Mean ± SEM | Statistical Test | Significance | Comparison | P value | Significance |
| | DAPI staining-stratum pyramidale thickness | WT | Veh | 3 | 63.48 ± 1.94 | Interaction: F (1, 10) = 0.3517, P=0.5663<br>Treatment: F (1, 10) = 0.8377, P= 0.3816<br>Genotype: F (1, 10) = 28.87, P=0.0003 | | WT Veh vs. Mut Veh | 0.00106 | * |
| | | | JNJ | 3 | 58.90 ± 1.57 | | | WT Veh vs. WT JNJ | >0.9999 | ns |
| | | Mut | Veh | 4 | 45.37 ± 3.70 | | | WT Veh vs. Mut JNJ | 0.0074 | ** |
| | | | JNJ | 4 | 44.39 ± 3.10 | | | Mut Veh vs. Mut JNJ | >0.9999 | ns |

| Extended Data Fig. 8e | Assay Performed | Parameter | Groups | N (animals) | Descriptive Statistics | Statistical Analysis | | Column statistics | | |
|---|---|---|---|---|---|---|---|---|---|---|
| | | | | | | Two-Way ANOVA | | Bonferroni's multiple comparisons | | |
| | | | | | Mean ± SEM | Statistical Test | Significance | Comparison | P value | Significance |
| | TUNEL staining-CA1 | WT | Veh | 4 | 7.95 ± 1.69 | Interaction: F (1, 12) = 25.48, P=0.0003<br>Treatment: F (1, 12) = 20.20, P=0.0007<br>Genotype: F (1, 12) = 106.8, P<0.0001 | | WT Veh vs. Mut Veh | <0.0001 | **** |
| | | | PER | 4 | 11.63 ± 1.64 | | | WT Veh vs. WT PER | >0.9999 | ns |
| | | Mut | Veh | 4 | 110.32 ± 5.81 | | | WT Veh vs. Mut PER | 0.0084 | ** |
| | | | PER | 4 | 46.82 ± 11.74 | | | Mut Veh vs. Mut PER | 0.0001 | *** |

| Extended Data Fig. 8f | Assay Performed | Parameter | Groups | N (animals) | Descriptive Statistics | Statistical Analysis | | Column statistics | | |
|---|---|---|---|---|---|---|---|---|---|---|
| | | | | | | Two-Way ANOVA | | Bonferroni's multiple comparisons | | |
| | | | | | Mean ± SEM | Statistical Test | Significance | Comparison | P value | Significance |
| | DAPI staining-stratum pyramidale thickness | WT | Veh | 4 | 64.84 ± 1.15 | Interaction: F (1, 12) = 6.012, P=0.0305<br>Treatment: F (1, 12) = 0.2245, P= 0.6441<br>Genotype: F (1, 12) = 114.7, P<0.0001 | | WT Veh vs. Mut Veh | P<0.0001 | *** |
| | | | PER | 4 | 61.18 ± 1.75 | | | WT Veh vs. WT PER | >0.9999 | ns |
| | | Mut | Veh | 4 | 40.54 ± 1.03 | | | WT Veh vs. Mut PER | P<0.0001 | *** |
| | | | PER | 4 | 45.94 ± 2.87 | | | Mut Veh vs. Mut PER | 0.3649 | ns |

| Extended Data Fig. 9a | Assay Performed | Parameter | Groups | N (animals) | Descriptive Statistics | Statistical Analysis | | Column statistics | | |
|---|---|---|---|---|---|---|---|---|---|---|
| | | | | | | One-Way ANOVA | | Bonferroni's multiple comparisons | | |
| | | | | | Mean ± SEM | Statistical Test | Significance | Comparison | P value | Significance |
| | Reciprocal social interaction | Sniffing | WT Veh | 17 | 47.82 ± 4.29 | Group (Genotype+treatment): F (3, 59) = 9.956, P<0.0001 | | WT Veh vs. Mut Veh | <0.0001 | *** |
| | | | WT ASO | 15 | 42.45 ± 2.65 | | | WT Veh vs. WT ASO | >0.9999 | ns |
| | | | Mut Veh | 15 | 24.36 ± 2.69 | | | WT ASO vs. Mut Veh | 0.004 | ** |
| | | | Mut ASO | 16 | 47.65 ± 3.66 | | | Mut Veh vs. Mut ASO | <0.0001 | *** |
| | | Following | WT Veh | 17 | 6.01 ± 0.60 | Group (Genotype+treatment): F (3, 59) = 6.442, P=0.0008 | | WT Veh vs. Mut Veh | 0.02 | * |
| | | | WT ASO | 15 | 5.25 ± 0.73 | | | WT Veh vs. WT ASO | >0.9999 | ns |
| | | | Mut Veh | 15 | 1.03 ± 0.22 | | | WT ASO vs. Mut Veh | 0.089 | ns |
| | | | Mut ASO | 16 | 8.14 ± 2.07 | | | Mut Veh vs. Mut ASO | 0.0004 | *** |
| | | Push-craw | WT Veh | 17 | 4.97 ± 0.91 | Group (Genotype+treatment): F (3, 59) = 3.261, P=0.0276 | | WT Veh vs. Mut Veh | 0.1405 | ns |
| | | | WT ASO | 15 | 4.77 ± 0.60 | | | WT Veh vs. WT ASO | >0.9999 | ns |
| | | | Mut Veh | 15 | 2.15 ± 0.62 | | | WT ASO vs. Mut Veh | 0.238 | ns |
| | | | Mut ASO | 16 | 5.79 ± 1.13 | | | Mut Veh vs. Mut ASO | 0.0262 | * |

| Extended Data Fig. 9b | Assay Performed | Parameter | Groups | N (animals) | Descriptive Statistics | Statistical Analysis | | Column statistics | | |
|---|---|---|---|---|---|---|---|---|---|---|
| | | | | | | One-Way ANOVA | | Bonferroni's multiple comparisons | | |
| | | | | | Mean ± SEM | Statistical Test | Significance | Comparison | P value | Significance |
| | Grooming | Grooming | WT Veh | 17 | 28.54 ± 4.79 | Group (Genotype+treatment): F (3, 59) = 7.766, P=0.0002 | | WT Veh vs. Mut Veh | 0.0017 | ** |
| | | | WT ASO | 15 | 26.54 ± 3.83 | | | WT Veh vs. WT ASO | >0.9999 | ns |
| | | | Mut Veh | 15 | 55.42 ± 6.02 | | | WT ASO vs. Mut Veh | 0.001 | *** |
| | | | Mut ASO | 16 | 27.13 ± 4.92 | | | Mut Veh vs. Mut ASO | 0.001 | ** |

| Extended Data Fig. 9c | Assay Performed | Parameter | Groups | N (animals) | Descriptive Statistics | Statistical Analysis | | Column statistics | | |
|---|---|---|---|---|---|---|---|---|---|---|
| | | | | | | One-Way ANOVA | | Bonferroni's multiple comparisons | | |
| | | | | | Mean ± SEM | Statistical Test | Significance | Comparison | P value | Significance |
| | EPM | Anxiety index | WT Veh | 17 | 65.48 ± 3.07 | Group (Genotype+treatment): F (3, 59) = 6.293, P=0.0009 | | WT Veh vs. Mut Veh | 0.0035 | ** |
| | | | WT ASO | 15 | 68.12 ± 2.81 | | | WT Veh vs. WT ASO | >0.9999 | ns |
| | | | Mut Veh | 15 | 81.20 ± 2.41 | | | WT ASO vs. Mut Veh | 0.0285 | * |
| | | | Mut ASO | 16 | 64.16 ± 3.74 | | | Mut Veh vs. Mut ASO | 0.0016 | ** |

| Extended Data Fig. 9d | Assay Performed | Parameter | Groups | N (animals) | Descriptive Statistics | Statistical Analysis | | Column statistics | | |
|---|---|---|---|---|---|---|---|---|---|---|
| | | | | | | One-Way ANOVA | | Bonferroni's multiple comparisons | | |
| | | | | | Mean ± SEM | Statistical Test | Significance | Comparison | P value | Significance |
| | MWM probe trial | Quadrant 1 | WT Veh | 17 | 13.09 ± 1.43 | Group (Genotype+treatment): F (3, 59) = 4.522, P=0.0064 | | WT Veh vs. Mut Veh | >0.9999 | ns |
| | | | WT ASO | 15 | 14.38 ± 1.95 | | | WT Veh vs. WT ASO | 0.0075 | ** |
| | | | Mut Veh | 15 | 24.05 ± 2.44 | | | WT ASO vs. Mut Veh | 0.0312 | * |
| | | | Mut ASO | 16 | 15.65 ± 3.10 | | | Mut Veh vs. Mut ASO | 0.078 | ns |
| | | Quadrant 2 | WT Veh | 17 | 25.22 ± 2.13 | Group (Genotype+treatment): F (3, 59) = 1.421, P=0.2456 | | WT Veh vs. Mut Veh | >0.9999 | ns |
| | | | WT ASO | 15 | 22.63 ± 3.27 | | | WT Veh vs. WT ASO | >0.9999 | ns |
| | | | Mut Veh | 15 | 27.92 ± 3.13 | | | WT ASO vs. Mut Veh | 0.9982 | ns |
| | | | Mut ASO | 16 | 20.71 ± 1.79 | | | Mut Veh vs. Mut ASO | 0.3446 | ns |
| | | Quadrant 3 (Target) | WT Veh | 17 | 42.36 ± 2.19 | Group (Genotype+treatment): F (3, 59) = 8.618, P<0.0001 | | WT Veh vs. Mut Veh | 0.0002 | *** |
| | | | WT ASO | 15 | 40.71 ± 3.01 | | | WT Veh vs. WT ASO | >0.9999 | ns |
| | | | Mut Veh | 15 | 25.87 ± 2.05 | | | WT ASO vs. Mut Veh | 0.0015 | ** |
| | | | Mut ASO | 16 | 41.20 ± 3.10 | | | Mut Veh vs. Mut ASO | 0.0008 | *** |
| | | Quadrant 4 | WT Veh | 17 | 19.32 ± 2.87 | Group (Genotype+treatment): F (3, 59) = 0.2889, P=0.8333 | | WT Veh vs. Mut Veh | >0.9999 | ns |
| | | | WT ASO | 15 | 22.28 ± 3.73 | | | WT Veh vs. WT ASO | >0.9999 | ns |
| | | | Mut Veh | 15 | 22.16 ± 2.76 | | | WT ASO vs. Mut Veh | >0.9999 | ns |
| | | | Mut ASO | 16 | 22.43 ± 1.78 | | | Mut Veh vs. Mut ASO | >0.9999 | ns |

| Extended Data Fig. 9e | Assay Performed | Parameter | Groups | N (animals) | Descriptive Statistics | Statistical Analysis | | Column statistics | | |
|---|---|---|---|---|---|---|---|---|---|---|
| | | | | | | One-Way ANOVA | | Bonferroni's multiple comparisons | | |
| | | | | | Mean ± SEM | Statistical Test | Significance | Comparison | P value | Significance |
| | MWM vision test | latency to platform | WT Veh | 17 | 9.26 ± 0.63 | Group (Genotype+treatment): F (3, 59) = 0.5431, P=0.6547 | | WT Veh vs. Mut Veh | >0.9999 | ns |
| | | | WT ASO | 15 | 10.88 ± 1.19 | | | WT Veh vs. WT ASO | >0.9999 | ns |
| | | | Mut Veh | 15 | 9.79 ± 1.05 | | | WT ASO vs. Mut Veh | >0.9999 | ns |
| | | | Mut ASO | 16 | 10.03 ± 0.80 | | | Mut Veh vs. Mut ASO | >0.9999 | ns |
